# Multi-influential genetic interactions alter behaviour and cognition through six main biological cascades in Down syndrome mouse models

**DOI:** 10.1101/2020.07.08.193136

**Authors:** Arnaud Duchon, Maria del Mar Muñiz Moreno, Sandra Martin Lorenzo, Márcia Priscilla Silva de Souza, Claire Chevalier, Valérie Nalesso, Hamid Meziane, Paulo Loureiro de Sousa, Vincent Noblet, Jean-Paul Armspach, Veronique Brault, Yann Herault

## Abstract

Down syndrome (DS) is the most common genetic form of intellectual disability caused by the presence of an additional copy of human chromosome 21 (Hsa21). To provide novel insights into genotype–phenotype correlations, we used standardized behavioural tests, magnetic resonance imaging (MRI) and hippocampal gene expression to screen several DS mouse models for the mouse chromosome 16 region homologous to Hsa21. First, we unravelled several genetic interactions between different regions of chromosome 21 and how they contribute significantly to altering the outcome of the phenotypes in brain cognition, function and structure. Then, in-depth analysis of misregulated expressed genes involved in synaptic dysfunction highlighted 6 biological cascades centred around DYRK1A, GSK3β, NPY, SNARE, RHOA and NPAS4. Finally, we provide a novel vision of the existing altered gene-gene crosstalk and molecular mechanisms targeting specific hubs in DS models that should become central to better understanding of DS and improving the development of therapies.

## Introduction

Down syndrome (DS) is the most common genetic form of intellectual disability and was first described as a disease by John Langdon Down in 1866. A century later, genetic studies demonstrated that DS is caused by the trisomy of human chromosome 21 (Hsa21) (1). People with DS have a wide range of phenotypic and physiological features with phenotypic variability, but they always present several disabling features like intellectual disability or Alzheimer’s disease (2). The leading cause of DS is the non-disjunction of chromosome 21 (3). However, in rare cases, people with partial Hsa21 duplications have been observed with a smaller spectrum of DS features. Studying these rare conditions increased our understanding of the genotype–phenotype correlations in DS (4–10): there is no single trisomic region responsible for all DS features, rather there are several susceptibility regions when presented in 3 copies that contributes to DS features in people with partial duplication of Hsa21. Consequently, this can induce a wide variety of features (5, 7). Nevertheless, several individuals displayed complex rearrangements such as contiguous or non-contiguous deletions or duplications, copy number variations of other regions or the duplication of genes located in the short arm of Hsa21. These rearrangements can potentially impact the phenotypic outcome of the Hsa21 duplication and add noise to the genetic dissection of clinical manifestations of human trisomy 21.

To circumvent the difficulties of studying DS in humans, several efforts have been made to generate DS mouse models (11). Indeed, there are three independent mouse regions homologous to Hsa21, carrying altogether 158 protein-coding homologous genes of the 218 protein-coding genes identified on Hsa21 (12). The largest region is found on mouse chromosome 16 (Mus musculus 16, noted as Mmu16) in a fragment of 22.42 Mb with 119 orthologous genes between *Lipi* and *Zbtb21* (13). The most telomeric part is split into two parts. The first part is found on mouse chromosome 17 (noted as Mmu17) with 19 homologous genes in the 1.1 Mb interval between *Umodl1* and *Hsf2bp*. Then, the second part is on mouse chromosome 10 (Mmu10) with 37 genes included in the 2.3 Mb *Cstb*-*Prmt2* genetic interval (11, 12, 14). Several DS mouse models have been generated over the years, most of them carried trisomy of the largest genetic region located on Mmu16 (11, 14). The Ts(17^16^)65Dn (noted Ts65Dn) model is the most widely used DS animal model and is quite unique, having a supplementary mini-chromosome obtained by x-ray irradiation of the male germline and containing the centromeric region of Mmu17, with genes from *Psid-ps2* to *Pde10a*, and the 13.5 Mb telomeric fragment of Mmu16 containing genes between *Mrpl39* and *Zbtb21* (15–18). Several models were made by chromosomal engineering (11) and carried the segmental duplication of Mmu16. The Dp(16*Lipi-Zbtb21*)1Yey (noted Dp1Yey) corresponds to the duplication of the entire Mmu16 region syntenic to Hsa21 (19). The Dp(16Cbr1-Fam3b)1Rhr (noted Dp1Rhr) model carries a duplication from *Cbr1* to *Fam3b* and demonstrates the contribution of the DS critical region (DCR) (20–23). All the DS mouse models displayed defects in behaviour and cognition which had been investigated in different laboratories with different protocols and environmental conditions, making inter-model comparison difficult (11).

To improve our knowledge of DS, we analysed seven DS mouse models that carry either large segmental duplication, like Dp1Yey, or a transgenic line overexpressing *Dyrk1a*, the main driver gene of the phenotype in mouse DS models, found on Mmu16 (24–28), and specific combinations of models (see Fig 1A). We used a unique and in-depth behaviour, morphological and transcriptomics pipeline in adult animals to dissect the contribution of the genes located on Mmu16 to DS mouse features. The behaviour pipeline relied on assessing specific hippocampal-dependent brain functions identified as altered in people with DS (29, 30). Thus, we performed standardized Y-maze, Open field (OF), Novel Object Recognition (NOR), Morris Water Maze (MWM) and contextual and cue Fear Conditioning (FC) tests. All the procedures were carried out in similar environmental conditions to reduce any impact of variation (31, 32). In addition, variations in specific brain regions have been observed in people with DS and mouse models (33–36). Neuroanatomical changes affect the whole brain volume or specific regions like the frontal region of the limbic lobe and the hippocampus in people with DS. Thus, we performed an in-depth morphological investigation of the brain by magnetic resonance imaging (MRI). Finally, whole gene expression was performed on hippocampi isolated from the six models to decipher the genes, functional pathways and biological cascades affected in the different DS mouse models.

**Figure 1.**
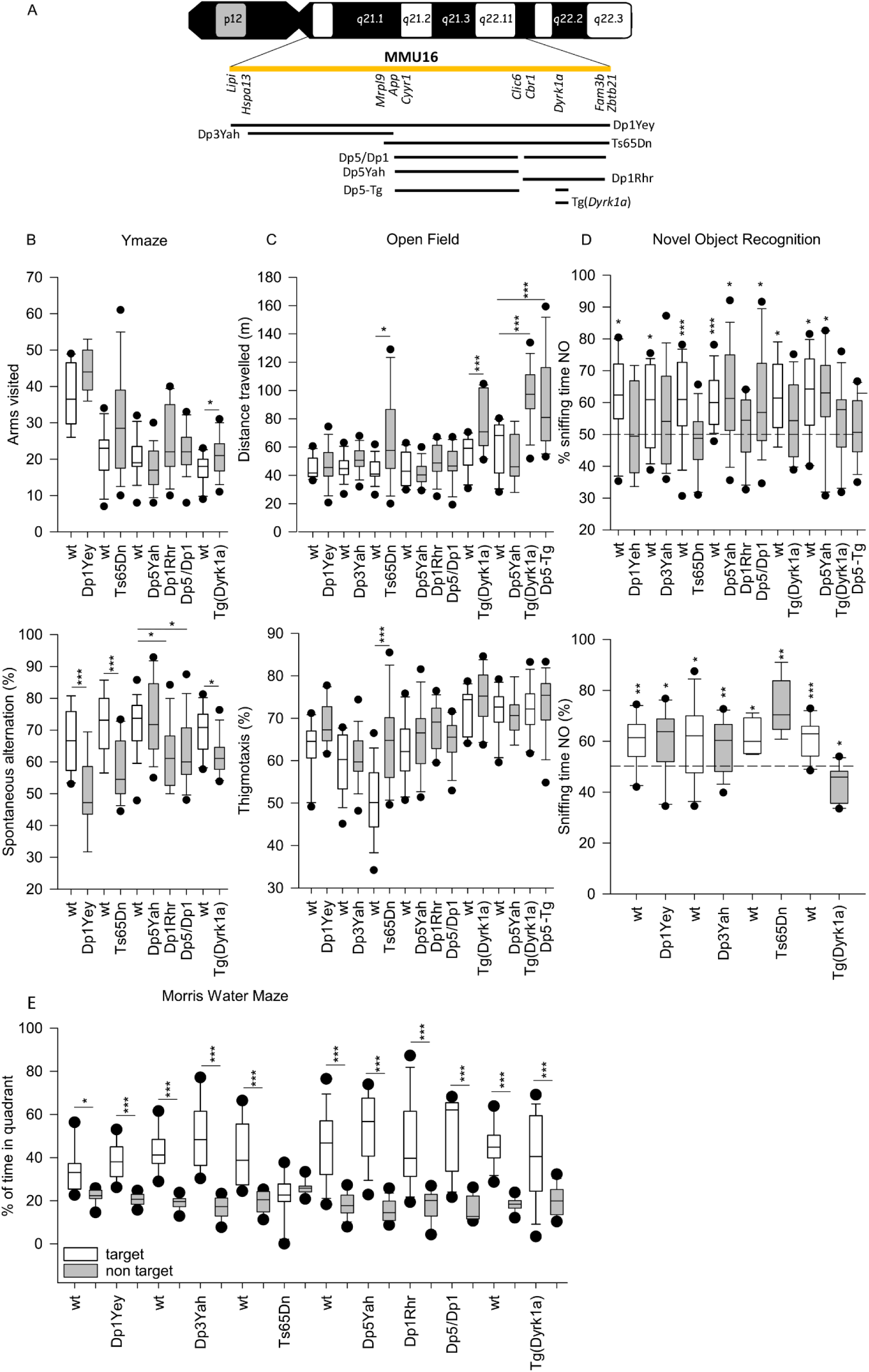
Down syndrome mouse models analysed in the study (A) and Standardized behavioural profiling (B-E). (A) In the upper part of the plot the human chromosome 21 is represented, in yellow we highlighted the Hsa21 syntenic region found in mouse from Lipi to Zbtb21 (known as Zfp295 previously). The eight models analysed on this study Dp1Yey, Dp3Yah, Ts65Dn, Dp5/Dp1, Dp5Yah, Dp1Rhr, Tg(Dyrk1a), Dp5yah crossed with Tg(Dyrk1a) (noted as Dp5-Tg)) trisomic chomosomal regions were draw in comparison with the Hsa21 region. (B) Y-maze spontaneous alternation. Arm visited (A upper panel) and alternation rate (A lower panel) are presented as box plots with the median and quartiles (upper and lower 90% confidence interval are indicated by a grey box). Only the Tg(*Dyrk1a*) mice were showed hyperactivity in this test with increased arms entries compared to the wild type (p=0,017). Alternation rate in Dp1Yey (p=0,002), Ts65Dn (p<0,001), Dp1Rhr (P=0,012), Dp5/Dp1Rhr (p=0,018) and Tg(*Dyrk1a*) (P=0,010) mice was significantly lower than respective wild-type mice (Dp1Yey n=10 wt and 10 Tg; Ts65Dn n=14 wt and 14 Tg, Dp5Yah/Dp1Rhr n=17 wt, 16 Dp5Yah, 15 Dp1Rhr and 17 Dp5Yah/Dp1Rhr; Tg(*Dyrk1a)* n=11 wt and 14 Tg). (C) Exploratory activity in a novel environment. Distance travelled (B upper panel) and % of distance travelled in peripheral zone recorded in the Open field arena (B lower panel). The total distance travelled was significantly higher in Ts65Dn (p=0,022), Tg(*Dyrk1a*) (p=0,008) and Dp5Yah/Tg(*Dyrk1a*) (p>0,001). Moreover, the % of distance in the peripheral zone was increased in Ts65Dn (p>0,001) mice comapred to wild type mice (Dp1Yey n=10 wt and 10 Tg; Dp3Yah n=15wt and 15 Tg; Ts65Dn n=14 wt and 14 Tg, Dp5Yah/Dp1Rhr n=17 wt, 16 Dp5Yah, 15 Dp1Rhr and 17 Dp5Yah/Dp1Rhr; Tg(*Dyrk1a*) n=11 wt and 14 Tg). (D) Novel Object Recognition with 24 hour (D upper panel) or 1 hour retention time (D lower panel). The results are presented as % of sniffing time (as box plots with the median and quartiles) for the novel object (NO). For 24 hours time laps, one sample t test vs 50% (hazard) showed that Dp1Yey (p=0,837), Dp3Yah (P=0,173), Ts65Dn (p=0;432), Dp1Rhr (p=0,492), Tg(*Dyrk1a*) (p=0,144) and Dp5Yah/Tg(*Dyrk1a*) (P=0,488) failed to recognize the new object. The Dp5Yah genomic fragment restored the capacity of the Dp1Rhr in the Dp5Yah/Dp1Rhr mice (p=0,0157; Dp1Yey n=10 wt and 10 Tg; Dp3Yah n=15 wt and 15 Tg; Ts65Dn n=14 wt and 14 Tg, Dp5Yah/Dp1Rhr n=17 wt, 16 Dp5Yah, 15 Dp1Rhr and 17 Dp5Yah/Dp1Rhr; Tg(*Dyrk1a*) n=11 wt and 14 Tg). For 1 hour retention time, all the mice were able to discriminate the NO except for the Tg(*Dyrk1a*) (p=0,011 preference for FO; Dp1Yey n=10 wt and 10 Tg; Dp3Yah n=15 wt and 15 Tg; Ts65Dn n=5 wt and 5 Tg; Tg(*Dyrk1a*) n=11 wt and 12 Tg). (E) Probe test session in Morris Water Maze. The results are presented as % of time in the target quadrant. All the mice have spent more time in the target quadrant versus non target excepted for the Ts65Dn mice (p=0,398) (* p<0.05, **p<0.01, ***p<0.001; Dp1Yey n=9 wt and 10 Tg; Dp3Yah n=15 wt and 15 Tg; Ts65Dn n=10 wt and 11 Tg, Dp5Yah/Dp1Rhr n=13 wt, 13 Dp5Yah, 13 Dp1Rhr and 13 Dp5Yah/Dp1Rhr; Tg(*Dyrk1a*) n=16 wt and 15 Tg).

## Results

### Dissecting the contribution of Mmu16 subregions to DS-related behavioral phenotypes in mouse models

We wanted to dissect the contribution of sub-regions located in the telomeric part of *Mus musculus* chromosome 16 (Mmu16), homologous to Hsa21 (37), to DS-related cognitive phenotypes. First, we selected 4 DS mouse models: Ts65Dn, the most commonly used DS model (15), and three additional models that carry segmental duplications of well-defined sub-regions located on Mmu16, Dp1Yey (19), Dp3Yah (38) and Dp1Rhr (20). In addition, we engineered a new sub-region, Dp5Yah, corresponding to three functional copies of the genes included in the genetic interval between *Cyyr1* to *Clic6*. This model was crossed with the Dp1Rhr sub-region to generate Dp5Yah/Dp1Rhr (noted Dp5/Dp1) compound transheterozygote carrying a trisomic gene content similar to Ts65Dn for the genes located on Mmu16. We also included a model carrying an additional copy of *Dyrk1a,* one of the driver genes for DS-related phenotypes (25, 35), and Tg(*Dyrk1a*) combined with the Dp5Yah model (noted Dp5-Tg) (See Fig 1A). We used standardized behavioral tests to study several aspects of learning and memory in mice, including the Y-maze (working memory), open field (exploration memory), novel object recognition (recognition memory), Morris water maze (spatial memory) and fear conditioning (associative memory) tests. For all the lines, independent cohorts of control and trisomic mouse littermates went through the pipeline at similar ages, after which the resulting data were processed using standard statistical analyses (see supplementary information for details). First, we assesed the potential existence of a background effect in the distribution of the measurements taken in the different tests. In our condition, the Q-Q plots with cumulative frequency were linear (see S1 table, S1 Fig) and thus, no notable difference between B6J or hybrid B6JC3B wild-type controls was observed.

Mouse activity and working memory were evaluated in the Y-maze (Fig 1B). The number of arm entries in the Y-maze showed that only the Tg(*Dyrk1a*) mutant line was hyperactive in this test while a deficit in spontaneous alternation was found in Dp1Yey (39), Ts65Dn (40), Dp1Rhr and Tg(*Dyrk1a*) (41). Dp5/Dp1 also showed a clear deficit in the percentage of spontaneous alternation in comparison to littermate controls while Dp5Yah trisomic animals showed normal performance. Thus the overexpression of *Dyrk1a* alone was sufficient to mimic the Y maze spontaneous alternation found in 3 overlapping DS models but we cannot rule out the possibility that another region is involved, as suggested by the work of Chang et al. (42).

Patterns of exploratory activity and anxiety were assessed in the open field (Fig 1C). Ts65Dn, Tg(*Dyrk1a*) and Dp5-Tg presented hyperactivity with an increased travelled distance compared to wild-type littermates, and support the results obtained in the Ymaze for the Tg(Dyrk1a) line. Thus the hyperactivity phenotype is linked to the Ts65Dn mouse model or only to the increase in Dyrk1a dosage.

The spatial reference memory was tested in the standard Morris water maze (MWM) task, in which mice have to escape from a circular pool of opaque water by localising a hidden platform at a single fixed location using distal spatial cues. We analysed the velocity, the distance travelled by the mice to reach the platform and thigmotaxis over-training (S2 Fig). The velocity of Dp1Yey and Dp5Yah was slightly lower than that of the wild-type mice (See S1 table, S2 FigB). As described previously (22, 43–49), Ts65Dn mice displayed a longer distance travelled to find the platform during all the sessions, compared to the wild type group (S2 FigA). Although Tg(*Dyrk1a*) mice were able to locate the platform, they also showed delayed acquisition compared to the control mice. Surprisingly, the Dp1Yey, Dp1Rhr, Dp5Yah and Dp3Yah mice completed this test without any difference with the wild type group. Interestingly, the Ts65Dn model, and to a lesser extent the Tg(Dyrk1a) model, presented marked thigmotaxic behaviour, spending a higher percentage of time in the peripheral zone in comparison to controls (S2 FigC). The retention of place location was evaluated during a single probe trial (PT) with no platform available, 24h after the last training session (Fig 1E). All the mouse strains except Ts65Dn remembered where the platform was located after the learning sessions. Finally, to check the visual ability of the mice, we performed a visual training version of the MWM during which the platform position was indicated by a flag. All the mice were able to find the visible platform without any significant difference with controls except for Tg(*Dyrk1a*) which presented a small delay in session 2 (S2 FigA).

We then evaluated non-spatial recognition memory using the novel object recognition (NOR) paradigm with a retention time of 24h. The percentage of sniffing time for the novel object was analysed and compared to 50% (hazard). This analysis showed that Dp1Yey, Dp3Yah, Ts65Dn, Dp1Rhr and Tg(*Dyrk1a*) were not able to discriminate between familiar and novel objects, unlike Dp5Yah and, more surpringly, Dp5/Dp1 (Fig 1D). To further characterize the effect/lack of effect of Dp5Yah mutation on novel object recognition, the Dp5Yah mouse line was crossed with the Tg(*Dyrk1a*) one and compared to new sets of wild-type, Dp5Yah and Tg(Dyrk1a) mice. Interestingly, we found that Dp5Yah/Tg(*Dyrk1a*) and, as expected, Tg(*Dyrk1a*), displayed altered novel object discrimination while Dp5Yah spent more time exploring the novel object than the familiar one. We also assessed the short term memory of Dp1Yey, Dp3Yah, Ts65Dn and Tg(*Dyrk1a*), by performing the NOR with a 1 hour delay between acquisition and retention; only the Tg(*Dyrk1a*) mice showed a deficit. Altogether, these data suggested that several loci are involved in the cognitive phenotype: one located in the Dp3Yah region and *Dyrk1a*; and presumably two interacting modifier loci: one located in the Dp5Yah region and another in the Dp1Rhr region.

All the trisomic lines were also tested for associative memory in the Pavlovian fear conditioning test. All the groups showed a higher percentage of freezing during the 2 min post-shock compared to the habituation session, indicating that all the groups developed fear responses during the training session (S3 Fig). When the animals were re-exposed 24 h later to the same context, the level of freezing in all the groups was increased compared to the habituation session (PRE2 and PRE4). However, the freezing time for Ts65Dn mice was lower compared to the respective control littermates. When we assessed cued fear conditioning in a new context, all the mice presented longer immobility time with a marked difference between pre-cue and cue periods (S3 Fig). In addition, Dp1Yey and, to a lesser extent, Ts65Dn showed lower freezing during the presentation of the cues in comparison to wildtype counterparts. As such, altered emotional associative memory was found in Ts65Dn and Dp1Yey but not in other DS models.Thus, depending on the behavioural traits observed, the overlapping DS models pointed to different regions with causative and modifier loci involved spread along the Mmu16 region homologous to Hsa21.

### Dissecting the contribution of Mmu16 sub-regions to the DS-related brain morphological phenotypes in mouse models

DS models have been reported to show brain morphological alterations of specific regions (35). Thus, we wondered if we could detect changes in brain morphology using MRI on these different partially overlapping trisomic mice models. Data were first analyzed using a volume approach and a brain region atlas. We confirmed that the brain of Tg(*Dyrk1a*) mice was larger (p<0.001) (35) and the brain of Dp1Yey mice was smaller than the respective wildtypes. Then, we analyzed different brain regions/structures taking into consideration the whole brain volume. Even with this correction, the Tg(*Dyrk1a*) mice were the most affected in terms of brain structure volume whereas, on the contrary, the Dp1Rhr mice did not show any significant variation compared to the wt mice. Several regions, such as the basal forebrain septum, central gray matter, the rest of the midbrain, and superior colliculi were significantly larger in the Tg(*Dyrk1a*), Ts65Dn and Dp1Yey DS models. Moreover, the cerebellum, hypothalamus, inferior colliculi and caudate putamen were significantly different in size for the Dp1Yey and Tg(*Dyrk1a*) carriers compared to controls (S2 table, S4 Fig), and a few additional areas were altered specifically in certain models (Amygdala, Globus pallidus, Hippocampus, Neo Cortex, and Thalamus for Tg(*Dyrk1a*); External capsule, Fimbria, and Ventricles for Dp1Yey). Altogether, this brain morphometric analysis showed greater similarity between the Dp1Yey and *Dyrk1a* overexpression transgenic models, with an intermediate overlap with the Ts65Dn mouse model.

### Dissecting the contribution of Mmu16 subregions to the DS-related transcriptome profiles in mouse models

Various studies have shown the consequences of trisomy on gene expression (50–59). Here we took the opportunity to dissect the alteration of gene expression and functional pathways in various DS trisomic models carrying different duplications of Mmu16. We analysed Ts65Dn, Dp1Yey, Dp3Yah, Dp5/Dp1, Dp5Yah, Dp1Rhr, and we included the trisomic model for *Dyrk1a* alone, Tg(*Dyrk1a*). Considering the hippocampal formation as a hub structure involved in learning and memory, we performed gene expression analysis in the adult hippocampus by comparing the DS models with their own littermate controls using a unique pipeline for all the models. For each DS model, we defined the expressed genes (noted as EGs) as the genes whose expression level was detected; the differentially expressed genes (noted as DEGs) as the genes whose expression level was found to be significantly altered in the trisomic model compared to the controls littermates; and then the trisomic expressed genes (TEGs) as the DEGs that are included within the duplicated chromosomal regions for each model (S3 table, Table 1).

**Table 1.** Differential expression analysis results of the seven models analysed. DEGs are Differential Expressed Genes and TEGs for Differential Expressed trisomic genes. * Analysis done with Mmu17 and 16 trisomic genes. GO are Go functional terms involved in cellular compartment (CC), molecular function (MF) and biological processes (BP).

| Mouse lines | Dp1yey | Dp3Yah | Ts65Dn<br>(Mmu16) | Dp5/Dp1 | Dp5Yah | Dp1Rhr | TgDyrk1A | Ts65Dn<br>(Mmu17) |
| --- | --- | --- | --- | --- | --- | --- | --- | --- |
| Nb of probes detected by the array |  |  |  |  | 35556 |  |  |  |
| Nb of annotated probes detected<br>(without control probes) |  |  |  |  | 27359 |  |  |  |
| Number of trisomic genes detected by<br>the array | 155 | 21 | 130 | 127 | 87 | 40 | 1 | 40 |
| Differential expressed genes (DEGs) | 711 | 826 | 1074 | 922 | 736 | 1306 | 850 |  |
| Differential expressed trisomic genes<br>(DETGs) | 66 | 13 | 64 | 54 | 39 | 18 | 1 | 28 |
| % of **DETGs | 43% | 62% | 49% | 43% | 45% | 45% | 100% | 70% |
| % of compensated trisomic genes<br>detected | 57% | 38% | 51% | 57% | 55% | 55% | 0% |  |
| Number of GAGE KEGG and GOs<br>(CC,BP,MF) terms disregulated in the<br>trisomic model FDR<0.1 | 244 | 67 | 12 | 111 | 318 | 225 | 231 |  |
| Number of GAGE KEGG and GOs<br>(CC,BP,MF) terms **upregulated** in the<br>trisomic model FDR<0.1 | 207 | 60 | 3 | 33 | 4 | 132 | 222 |  |
| Number of GAGE KEGG and GOs<br>(CC,BP,MF) terms **downregulated** in<br>the trisomic model FDR<0.1 | 37 | 7 | 9 | 78 | 314 | 93 | 9 |  |
| Number of GAGE KEGG and GOs<br>(CC,BP,MF) terms in the trisomic<br>model unique to each mice line<br>FDR<0.1 | 80 | 1 | 2 | 30 | 195 | 107 | 64 |  |

\*DEGs/DETs: Diferential expressed genes/ Diferential expressed \*\*DETGs/DETTs: Diferential expressed trisomic genes/

Although most of the genes in 3 copies were overexpressed in the relevant mouse model-derived hippocampi with a ratio around 1.5 (S5 Fig), from 38% to 57% of the trisomic genes showed a dosage compensation (Table 1, S3 table). While this compensation was expected, we noticed that most of the compensated genes behaved similarly in the different trisomic contexts, although the experiments were performed independently. As such, the genes from *Cldn17* to *Krtap11-1*, including the keratin cluster were not overexpressed when trisomic in any model (Dp1Yey, Ts65Dn, Dp5/Dp1 or Dp5Yah). This could have been due to the fact that this region seems to be under strong regulatory constraints, as on the borders two REST sites and a LaminB1 peak encompassing this region were found (UCSC browser), while *Btg3* (*BTG Anti-Proliferation Factor 3*) and C21orf91 (*Chromosome 21 open reading frame 91*, also known as *D16Ertd472e*), and *Mrpl39 (Mitochondrial Ribosomal Protein L39), Jam2 (Junctional Adhesion Molecule 2), Atp5J (ATP synthase peripheral stalk subunit F6), Gabpa (GA binding protein transcription factor subunit alpha)* and *App (amyloid beta precursor protein)* were overexpressed in various DS models. We also found that the genes located on the trisomic segment not homologous to Hsa21 on Mmu17 in Ts65Dn hippocampi, were overexpressed (S6 Fig). This was also observed in the Ts65Dn heart in a previous study (16). Looking in detail at the homologous Hsa21 region in Mmu17, we saw two main genetic effects due to the overdosage of the Mmu16 region homologous to Hsa21. Noticeably, *Cbs,* coding for the Cystathionine beta-synthase, another driver gene for DS cognitive phenotypes (60), was found to be down-regulated in all the models, except Dp3Yah and Tg(*Dyrk1a*), suggesting direct control of *Cbs* transcription by at least two loci, one located in Dp5Yah and another one, not due to *Dyrk1a* overdosage, in the Dp1Rhr trisomic region. Similarly, under-expression of the *glucagon like peptide 1 receptor (Glp1r)* was observed in Dp1Yey, Ts65Dn, Dp5/Dp1 and Dp5Yah. On the contrary, this gene was overexpressed in Dp1Rhr and not affected in Tg(*Dyrk1a*).

Here too, *Dyrk1a* dosage was not involved, but at least two loci controlling *Glp1r* expression with opposite and epistatic effects should be found respectively in the Dp5Yah and Dp1Rhr genetic intervals. Thus, a complex genetic interaction took place between different loci, controlling subsequent gene expression.

The analysis of DEGs in each model by principal of component analysis (PCA), t-SNE (Fig 2B) or OPLS techniques (see supplementary information) highlighted the capabilities to separate trisomic individuals from wild-type littermates (Fig 2A). Genome-wide misregulation was found independently of the model, as DEGs were spread in all the chromosomes (S3-S4 tables, S7 Fig), as shown previously (56), although with a stronger impact of the Dp1Rhr on the total number of DEGs detected.

**Figure 2:**
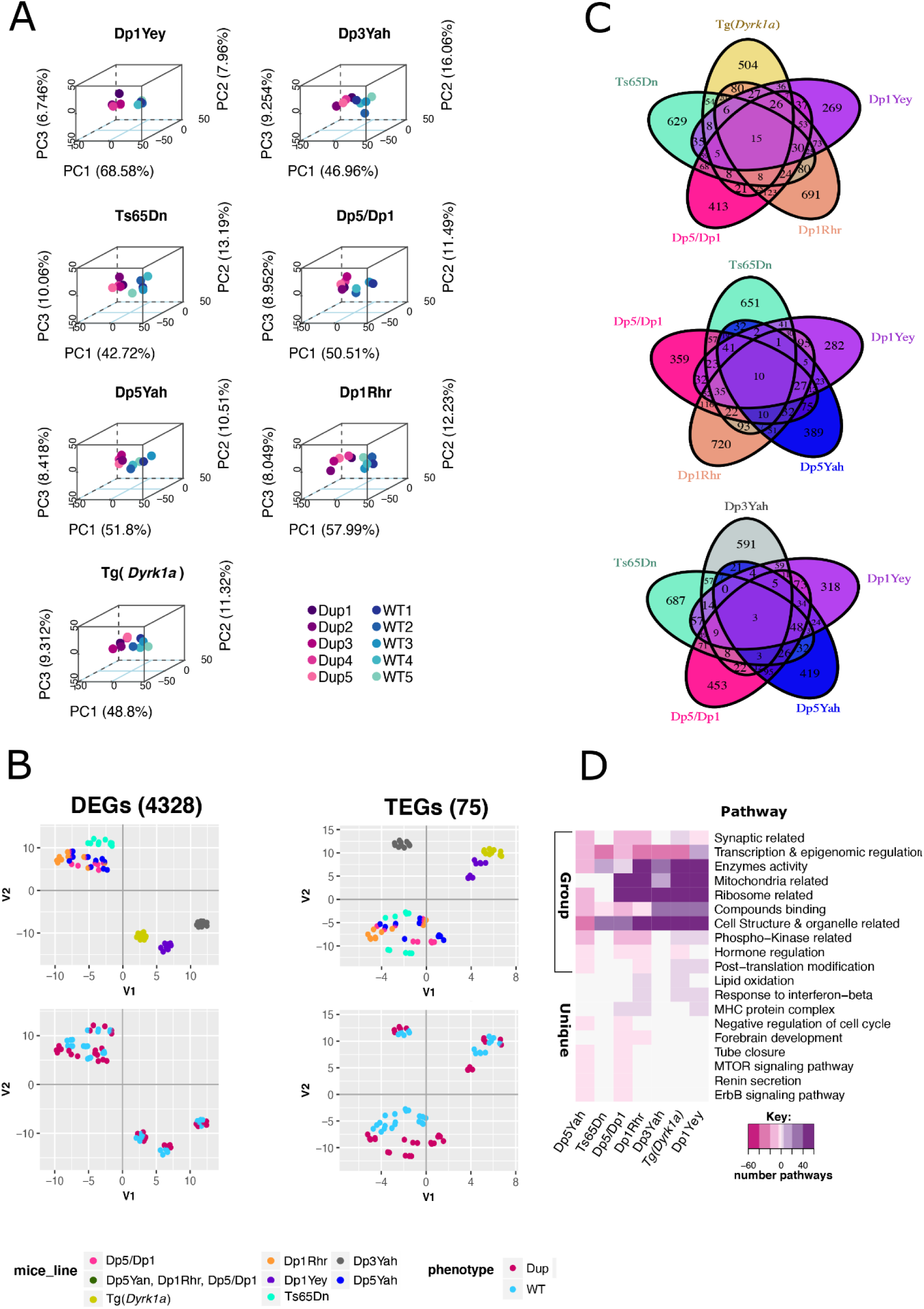
The differential expression analysis discriminates trisomic from disomic hippocampi and identifies common dysregulared genes and pathways. (A) 3D-PCA on the DEGs for each sample allows to separate trisomic (Dp) from disomic (wt) adult hippocampi. (B) Left column: two dimensional Principal Component Analysis (2D-PCA) on the 5599 transcripts of the 4328 DEGs over all the samples identified using fcros 0.025<α<0.975. Right column: 2D-PCA on the 75 trisomic genes with a measured expression in all the models. (C) Venn Diagrams showing the overlap in gene expression between the different mouse lines represented in different colours. (D) Heatmap representation of the number and regulation sense of the pathways shared at least by two mice lines identified using the genome expression for each mice line by GAGE R package and filtered by q-value cut off < 0.1. Grouped in the categories showed on the ordinate: synaptic related, synaptic related: representation of the pathways involved in Myelin sheath and SNARE complex formation, synaptic related: all the synaptic related pathways excluding myelin sheath and SNARE complex formation, Transcription & epigenomics regulation, Enzymes activity, Ribosome related, Mitochondria related, Cell Structure & organelle related, Phospho-kinase related… The color key breaks represents the number of pathways within the categories 60,40,20,5,0.5. The minus or pink color represents down regulated pathways, the white color represents no pathway found in the category and the purple or positive numbers stands for up regulated pathways respectively.

The most overexpressed genes in terms of log2 fold change (log2FC) of expression and significance in various genetic conditions were visualised using Volcano plots (S8 Fig, S4 table). For example, the *listerin E3 ubiquitin protein ligase 1* (*Ltn1*) gene, coding for a major component of ribosome quality control and causing neurodegeneration in mice (61), was found overexpressed in Dp1Yey, Ts65dn, Dp5/Dp1, and Dp5Yah or *Ifnar2*, coding for the Interferon Alpha and Beta Receptor Subunit 2, was overexpressed as expected in models that carried three copies of this gene (Dp1Yey, Ts65Dn, Dp5/Dp1 and Dp5Yah). Instead, a more controlled gene like the *neuronal acetylcholine receptor subunit alpha-3* (*Chrna3*), was found upregulated only in Dp1Rhr and Dp1/Dp5, certainly due to the overexpression of one gene from the *Cbr1-Fam3b* region but not *Dyrk1a*. Nevertheless, when we performed the intersection between the list of DEGs from the different models, we found only a few genes in common (Fig 2C, S4 table).

We decided to combine the analysis of all the lines together using PCA and t-SNE and revealed a strong clustering of models that shared at least a partially duplicated region (Fig 2B). t-SNE analysis, based on all the 4328 DEGs detected in each mouse model added together, showed different contributions of the various DS models to the transcriptome variation (Fig 2B, left panel) with 2 distinct groups: one encompassing four overlapping trisomies: Ts65Dn, Dp5/Dp1, Dp5, Dp1Rh and three isolated models: Dp1Yey, Dp3Yah and Tg(*Dyrk1a*) that were closer together, although Dp3Yah was clearly farthest from the other two. Similar distinct groups were seen when analysing the TEGs (Fig 2B, right panel) and overall, the trisomic and the wild-type individuals in each mouse line were nicely separated. As expected, the expression level of the TEGs and the DEGs in the different trisomic conditions were strongly correlated (S9 Fig). Interestingly, the 4328 DEGs showed a level of misregulation strongly correlated between Dp1Yey and Dp3Yah (33%), Dp5/Dp1 (50%), Dp1Rhr (40%) and Tg(*Dyrk1a*) (42%). Of the 75 genes detected and located on Mmu16 region homologous to Hsa21, the correlation of fold change was around 28% in the Dp1Yey and Tg(*Dyrk1a*) partial DS models. Thus, the correlation in gene deregulation showed that *Dyrk1a* overdosage is a key driver of transcriptome deregulation in the Dp1Yey and Dp1Rhr models.

Unexpectedly, the correlation of DEGs mis-expression level was lower between Ts65Dn and Dp1Yey (29%) or Dp1/Dp5 (28%). On the contrary, a large number of TEGs were misregulated in the same way between Ts65Dn and Dp1Yey (49%) and Dp1/Dp5 (52%; S9 Fig), suggesting that the other region found in 3 copies in the Ts65Dn over Mmu17 must affect the general DEG landscape.

Using qRT-PCR, we confirmed the mRNA overexpression of first, *Dyrk1a* and *Sod1* genes in the DS models where they were trisomic; second, of *Synaptojanin2 (Synj2*) and *T lymphoma invasion and metastasis inducing gene 2 (Tiam2*) located on the Mmu17 centromeric region in Ts65Dn, and third, of *Cholinergic Receptor Nicotinic Alpha 1 Subunit (Chrna1),* a gene misregulated in the Dp1Yey, Dp5/Dp1, Dp1Rhr and Ts65Dn models (S15 fig), and *Cbs* downregulated in all the models except Tg(*Dyrk1a*) and Dp3Yah,. We also confirmed alterations of the expression of immediate early-response genes *Arc, FosB, Fos* and *Npas4* that are important for cognition.

### Differential functional analysis unravels a few common altered pathways in DS models

To go further, we performed a differential functional analysis and found 12 to 318 misregulated pathways in the DS models (table 1, S5 table). Interestingly, the regulation of pathways is trisomic region-dependent, as the Dp5Yah (99%) region produced an overall downregulation whereas the Dp3Yah (89.5 %) and Dp1Rhr (56 %) trisomy together with the full Hsa21 syntenic model Dp1Yey (84.8 %) induced preferentially an upregulation.

To facilitate understanding, we clustered the broad functional dysregulation into 8 major functionality groups or meta-pathways. We found ribosomal components and mitochondrial process pathways altered in all the models, with many genes shared between models (S11 Fig). Cell structure and organelles, transcription and epigenetic regulation, interferon and synaptic meta-pathways were more affected in some models than in the others (S10 and S12 Fig). As such, we observed strong and connected effects in the control of transcription and epigenetic regulation, enzyme activity and cell structure, and cellular organelles involved in membrane and protein processing (endoplasmic reticulum, Golgi body, lysosome, peroxisome, etc.; Fig 2D) in the Dp1Yey, Dp5/Dp1, Dp1Rhr, and Tg(*Dyrk1a*) models, whereas the myelinization and 10 SNARE components, such as the *Synaptosome Associated Protein genes 25 and 23* (*Snap25* and *Snap23)*, were specifically dysregulated in the Dp1Yey Dp5Yah and Tg(*Dyrk1a*) models.

Interestingly, we saw many shared genes between these pathways and the models, giving rise to high pathway connectivity between models (see Materials and Methods). Considering the DEGs involved in brain synaptic pathways, with the DS synaptic MinPPINet (Fig 3A), we analysed the DS network topography and betweenness connectivity and found hubs and genes more central for network information flow. As expected from a PPI biological network, the likelihood of observing such connectivity in the DS network was more than one can expect by chance (P-value < 2e-16) and it showed a small world effect and scale-free topology. Using a network decomposition approach (see supplementary information), we highlighted 6 major subnetworks or biological cascades that strongly centralized 6 different proteins: DYRK1A, GSK3B, NPY, RHOA, SNARE and NPAS4 proteins (Fig 3B-C). As a summary, *Dyrk1a* was an upregulated in Dp1Yey,Ts65Dn, Dp5Dp1, Dp1Rhr and Tg(Dyrk1a) while *Npas4* was downregulated in Ts65Dn, Dp1Rhr and Tg(Dyrk1a), and *Npy* was upregulated in Dp1Yey, Dp5Dp1, Dp5Yah, Dp1Rhr, and Tg(Dyrk1a), and downregulated in Dp3Yah. Ten genes from the SNARE complex were dysregulated in some DS partial models; from these we validated the disregulated expression of *Snap25* and *Snap23* by qRT-PCR (Fig 4F). Gsk3b and Rhoa were not DEGs but these two were hubs interacting with many DEGs in the network.

**Figure 3:**
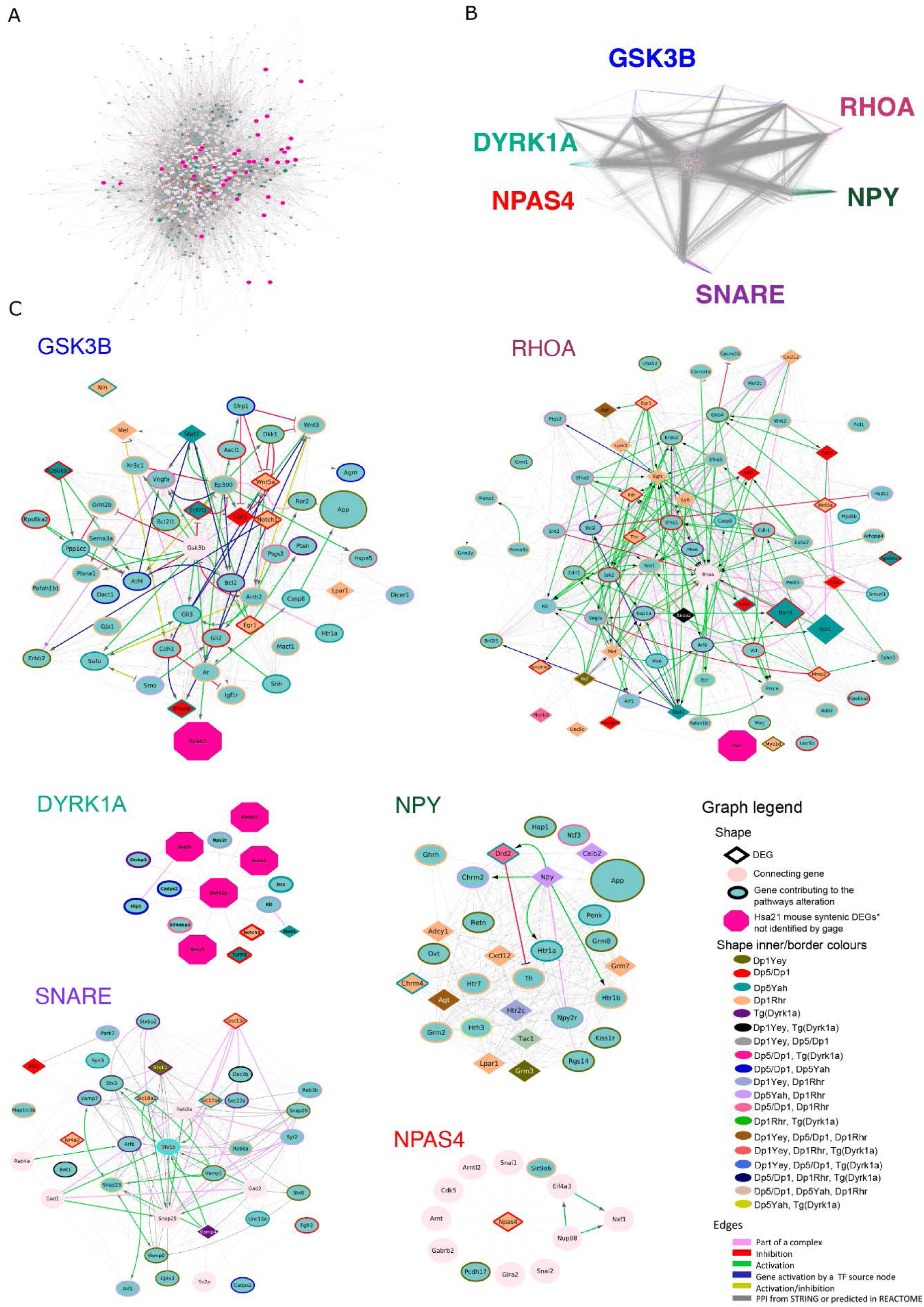
Protein-protein interaction networks involving DEGs linked to the synaptic function. (A) STRING04 MinPPINet of genes involved in synaptic function visualized using the edge weighed spring embedded layout by betwenness index in Cytoscape. The network was built by querying STRING and selecting the PPIs with a medium confidence score (CS=0.4) coming from all sources of evidence. The shapes of the nodes represent the following information: Shapes: i)Pallid pink ellipses: represent connecting proteins added to assure the full connectivity of the network; ii) pink octagons, represent HSA21 syntenic genes in mouse not identified as contributing to the meta-pathway dysregulation by GAGE; iii) green inner coloured ellipses, genes identified by GAGE after q-val <0.1 cut off to be contributing even slightly, to any pathway of those found dysregulated inside the meta-pathway. If the size is similar to the octagons, they are also HSA21 syntenic genes in mouse. Additionally, the border colour represents the mouse model multi group where those genes are found altered in; iv) diamonds, genes identified by GAGE after q-val <0.1 cut off and also by FCROS as DEGs. (B) Network Structure Decomposition of the STRING04 MinPPINet. Highlighting in different colors the interactions of GSK3B, NPY, SNARE proteins, DYRK1A and RHOA respectively. In the case of NPAS4, the interactions coloured correspond up to the first level interactions. (C) The six RegPPINets were extracted from the selection of each fo the following proteins and their 2^nd^ interactors from STRING04 MinPPINet: RHOA, DYRK1A, GSK3B, NPY, SNARE proteins and NPAS4. Then, those were further annotated with regulatory information using REACTOME (See Supplementary information). The shapes of the nodes represent the following information: Shapes: i)Pallid pink ellipses: represent connecting proteins added to assure the full connectivity of the network; ii) pink octagons, represent HSA21 syntenic genes in mouse not identified as contributing to the meta-pathway dysregulation by GAGE; iii) green inner coloured ellipses, genes identified by GAGE after q-val <0.1 cut off to be contributing even slightly, to any pathway of those found dysregulated inside the meta-pathway. If the size is similar to the octagons, they are also HSA21 syntenic genes in mouse. Additionally, the border colour represents the mouse model multi group where those genes are found altered in; iv) diamonds, genes identified by GAGE after q-val <0.1 cut off and also by FCROS as DEGs. The edges colored represent the type of interaction annotated by following the PathPPI classification (Tang *et al.* 2015),and ReactomeFIViz annotations as follows i) The GErel edges indicating expression were colored in blue and repression in yellow. ii) PPrel edges indicating activation were coloured in green, inhibition in red. Iii) Interactions between proteins known to be part of complexes in violet. Iv) Predicted interactions were represented in grey including the PPI interactions identified by STRING DB (Szklarczyk *et al.* 2017) after merging both networks.

**Figure 4:**
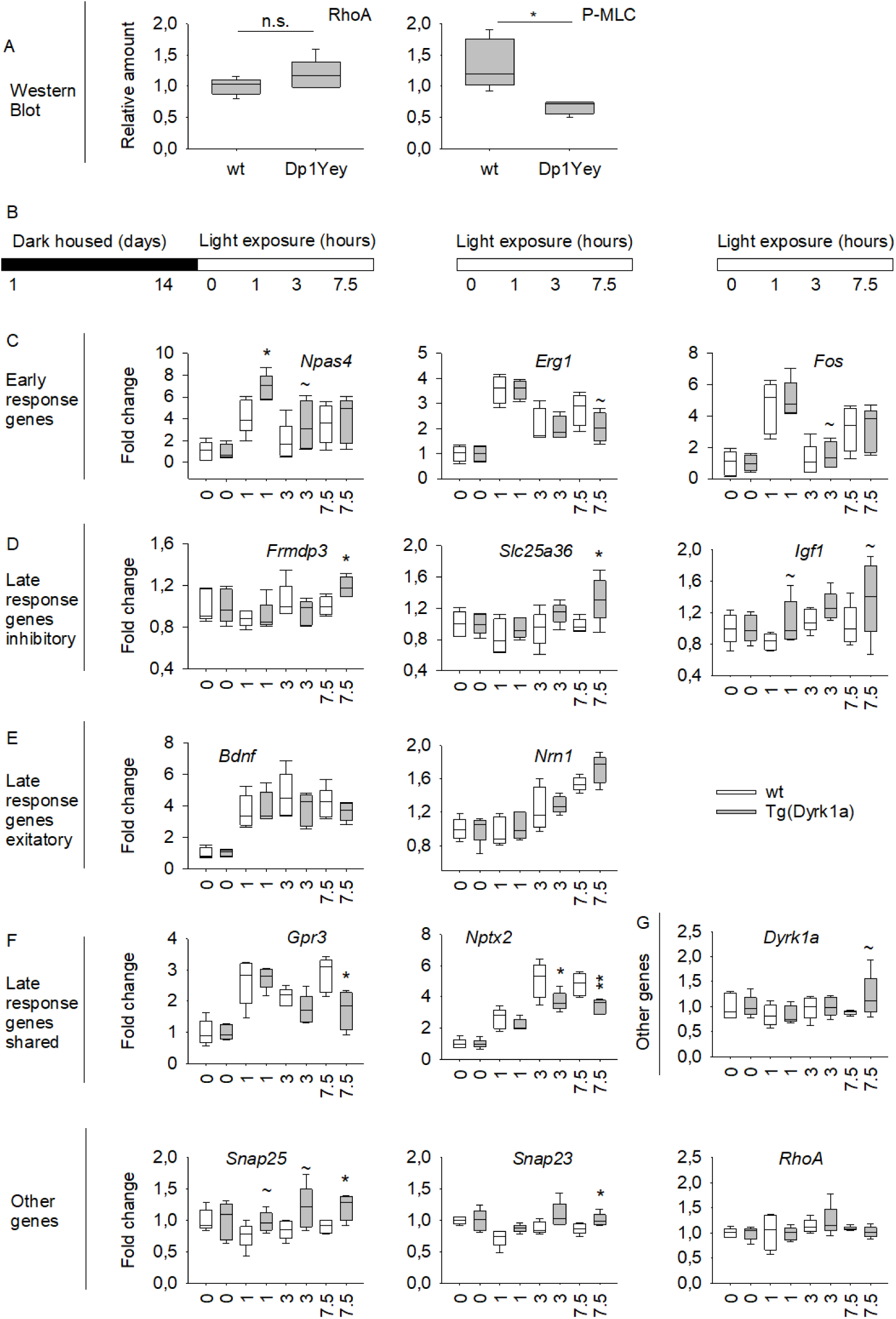
Evaluating Npas4 and RhoA pathways in DS models. (A) RHOA pathway was altered in the Dp1Yey. Western blot analysis was revealed no changes in the expression of RHOA but a significant decrease of the phosphorylation of the Myosin light chain (P-MLC) in the Dp1Yey hippocampi compared to control (n=5 per genotype). (B) Mice were housed in total darkness for 14 days and then were subsequently exposed to light for 0, 1, 3 or 7.5 h. Relative expression levels were determined, and fold change were calculated for each condition. Genotypes differences in fold change were assessed by T Test. (C) Only the fold change for early response genes *Npas4* was up-regulated in Tg(*Dyrk1a*) mice compared to wt at 1 hours of light induction. (D) The late responses genes specific to inhibitory neurons *Frmdp3*, *Slc25a36* and *Igf1* were up-regulated after 7.5 hours of light induction. (E) The fold change of late responses genes specific to excitatory neurons *Bdnf* and *Nrn1* were unchanged. (F) The fold change of late response genes shared by excitatory and inhibitory neurons *Gpr3* and *Nptx2* were downregulated after 3 and/or 7.5 hours of light induction. (G) *Dyrk1a* and *RhoA* showed a similar fold change along the different condition whereas *Snap25* and *Snap23* presented an increased enrichment for the 7.5 hours condition. Data are presented as box plots with the median and quartiles.

Overall, the network analysis of the DS synaptic MinPPINet showed that DYRK1A controls 42.3 % of the network nodes and 69.4 % of the network seeds via 2^nd^ level interactors. Hence, DYRK1A could control the DS synaptic network via PPI and regulatory interactions. Furthermore, the biological cascades centred on GSK3B, DYRK1A and RHOA are highly interconnected (S13 and S15 figs) and in fact several interactors of RHOA are connected and could somehow modulate a higher percentage (75 and 68.5%) of the nodes of the network and synaptic seeds (S6 table).

### Validating the newly identified RHOA, NPAS4 and SNARE pathways in DS Models

RHOA is a small GTPase protein acting through the activation of ROCK (RhoA Kinase) and the phosphorylation of the myosin light chain (MLC). Interestingly, *RhoA* was not found altered in the differential expression analysis; instead, it was a connecting node introduced to obtain a full connected PPI network o ascertain whether RHOA pathway was altered in Dp1Yey, we checked the expression levels of two proteins of the pathway by qRT-PCR and WB (Fig 4A). We found no changes in the expression of RHOA, converging with the transcriptomic analyses, but we detected a significant decrease of MLC phosphorylation (P-MLC) in the Dp1Yey hippocampi compared to control (S16 fig). Thus, the RHOA pathways appeared to be downregulated in the Dp1Yey DS mouse model.

In our transcriptomics and network analysis, we found that *Npas4* was misregulated in the Tg(*Dyrk1a*), Dp1Rhr and Ts65Dn models. We verified the downregulation of *Npas4* and several other immediate early genes (IEGs: *Arc*, *Fosb* and *Fos*) in DS mouse models (S15 Fig). It is known that these IEGs are activated when light exposure is induced after a long light deprivation period (62, 63). To confirm the impact of *Npas4* downregulation in Tg(*Dyrk1a*) mice, we performed qRT-PCR experiments to determine the specific early and late response genes altered in the visual cortex after light deprivation and de novo light exposure at 3 time points (1, 3 and 7.5 hours) (Fig 4B). The results showed that *Npas4* was clearly induced after light deprivation following 1 hour of light stimulation but the expression level of *Npas4* was higher in Tg(*Dyrk1a*) mice compared to control (Fig 4C). We also took the opportuntity to observe the expression of late response genes specific to inhibitory neurons (*Frmdp3*, *Slc25a36* and *Igf1,* Fig 4D) and late response genes (*Grp3* and *Nptx2,* Fig 4F). We found that gene expression was altered after 7.5 hours of light stimulation in Tg(*Dyrk1a*). Interestingly, late response genes specific to exitatory neurons (*Bdnf* and *Nrn1,* Fig 4E) were not affected. The *Snap25*, *Snap23* candidate genes found in our analysis showed an altered expression after 7.5 hours of light stimulation while *Dyrk1a* and *RhoA* levels were not affected (Fig 4F).

## Discussion

In this study, we explored five DS mouse models carrying 3 copies of the region homologous to Hsa21 found on Mmu16, to decode the DS genotype-phenotype relationships and further investigated genetic interactions between different regions. To this end, we also assessed a transgenic model overexpressing one copy of *Dyrk1a*, plus two combinations of models (Dp5/Dp1, Dp5-Tg, Fig 1), using a standardized behavioural pipeline focused on hippocampus-dependent memory processes; a process found impaired in people with DS.

In this parallel comparison, we observed several loci contributing to the alteration of different brain memory and control functions. We found that the spontaneus alternation in the Y-maze was altered in most of the models (except Dp5Yah alone; Fig 1B). The minimal common genetic part of these lines was the overexpression of *Dyrk1a* and the result observed for transgenic Tg(*Dyrk1a*). Altogether, our previous results support DYRK1A as being the main driver of working memory defect (25) although another region could be involved in controlling spontaneous alternation (42).

Similarly, DYRK1A overdosage is a major cause for increased activity in the open field in Dp5/Tg(*Dyrk1a*) and Tg(*Dyrk1a*), but not in other models. While the Dp1Rhr Ds model was not affected, it can be hypothesized that another locus, located in the *Cbr1-Fam3b* interval, inteferes with *Dyrk1a* overdosage in this model. The situation should be even more complex with additional genetic interaction. Indeed, no phenotype was observed for the distance travelled in the Dp1Yey, Dp5/Dp1, Dp5Yah, Dp1Rhr models. Thus the overexpression of Dyrk1a was able on its own, or combined with Dp5Yah, to induce increased activity while some loci that were not trisomic only in the Dp5/Dp1 model were able to suppress this effect, which can be reinduced by other trisomic loci specific to the Ts65Dn (see below). Altogether, our results suggest that many different genetic influences (at least 3 for the distance travelled) act on different behavioural variables in DS models.

Similar analysis of the NOR test with 2 distinct retention times highlighted several loci. The NOR test with 24h of retention unravelled deficiency in most of the models, except in Dp5Yah and Dp5/Dp1, suggesting that there are at least two causative and two modifier loci (Fig 5). Taken together, our results suggest that, depending on the variable observed in the behavioural test, several genetic interactions occur to build the link of behavioural phenotyping outcome in DS mouse models with loci, spread along Mmu16, including *Dyrk1a*. With 1h of retention time, the NOR test pointed only to Dyrk1a overexpression, with at least one suppressing loci in the Dp1Rhr trisomic region.

**Figure 5:**
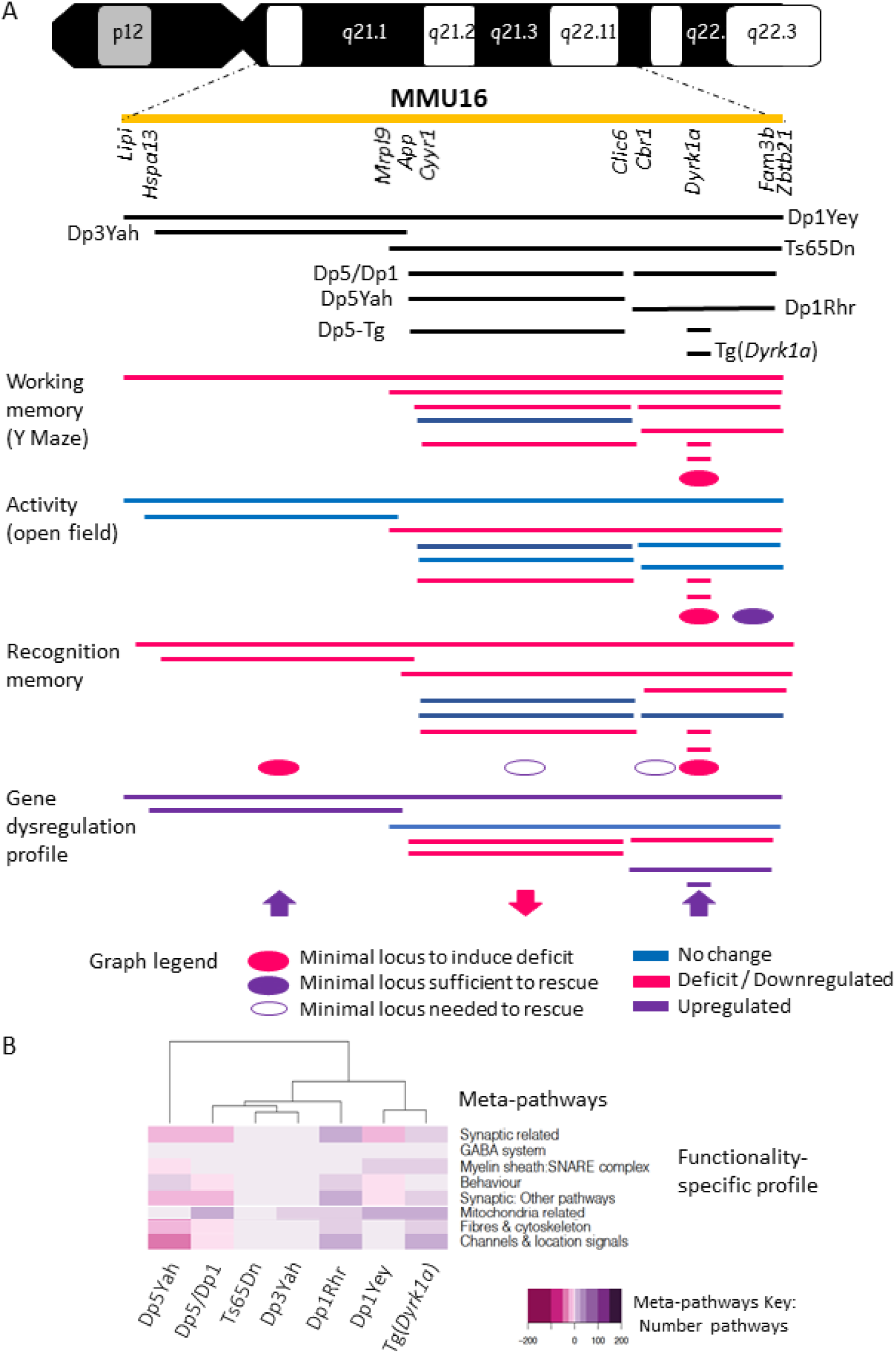
Genotype correlation associated to behaviour phenotype in partial trisomic DS model. Here we highlight the duplicated region carried on each model with the corresponding syntenic region in the human chromosome 21 together with the main behavioral and transcriptomics results pointing to the existence of region specific phenotypes and functional alteractions. The black lines represents the duplicated syntenic regions to human chromosome 21 on each model (represented in the yellow line). The blue lines represents the behavioral results where no alteration was found, instead the red lines identifed the tests with deficits. Over the transcriptomics meta-pathways fucntional profile summary picture, in purple is highlighted upregulation whereas in pink downregulation. The intensity of the color stands for the number of pathways included on each meta-pathway from the total number of pathways found altered on each model.

As expected, we found deficits in the Ts65Dn mice similar to those previously published in the Y-maze, the open field, NOR, MWM and contextual fear conditioning (40, 64) tests. Strikingly, thigmotaxis and time in the target quadrant in the probe test of the MWM were two variables modified only in the Ts65Dn trisomic model while Dp1Yey, which carries a duplication of the complete syntenic Mmu16 region, was less affected, as described before (65, 66). Remarkably, the Dp5Yah mice, with a duplication from *Cyyr1* to *Clic6*, displayed no deficits on their own in any of the tests performed and showed the lower number of DEGs in the hippocampi. Although several genes, *Sod1*, *Olig1*, *Olig2, Rcan1* and *Synj1*, from the region were proposed as inducing an early DS cognitive phenotype (67–70), our results indicated that the *Cyrr1-Clic6* region was not sufficient to induce by itself cognitive defects in DS models. Nevertheless, no defects were found in the Dp5/Dp1 mice, contrary to Ts65Dn in the open field, or they appeared less severe in the NOR test (see below). This indicates the existence of a key modulator in the *Cyyr1-Clic6* region. The major behavioural alterations found in Ts65Dn probably result from the influence of different genetic factors: first the overdosage of genes homologous to Hsa21; then the presence of the freely segregating mini-chromosome (71), and also the trisomy of about 60 Mmu17 centromeric genes, non homologous to Hsa21(16), and overexpressed in the hippocampus of Ts65Dn mice. In particular, the overdosage of *Tiam2* and *Synj2,* located on the Mmu17 centromeric region could exacerbate the effect of the overexpression of their respecitve paralogs *Tiam1* and *Synj1* (70, 72). Strikingly, the transcriptomic analysis showed a different global disruption of genome expression in the Ts65Dn hippocampi, compared to the other trisomic models with segmental duplications. This was emphasized by the low correlation of DEGs and deregulated pathways between Ts65Dn, Dp1Yey and Dp5/Dp1. Overall, we can hypothesize that the suppression effect seen in the Dp1Yey model compared to the Ts65Dn, is due to a suppression effect of genes overexpressed and located upstream of *Mrpl39*, or to an enhancing effect of genes located on the non-homologous region in the Ts65Dn minichromosome, or to the freely segregating minichromsome in the Ts65Dn. New models and further analysis will be needed to test these hypotheses.

Similar to cognition, brain morphology was affected differently in DS models. As observed in people with DS, a global decrease in brain size was observed in Dp1Yey, Ts65Dn and Tg(*Dyrk1a*) while Dp1Rhr showed an increase in size in many brain regions compared to the other DS models. In addition, specific changes were found in several regions including the basal forebrain septum, a predominant source of cortical cholinergic input with an early substantial loss of basal forebrain cholinergic neurons (BFCN); a constant feature of Alzheimer’s disease and other deficits in spatial learning and memory (73). Besides, only Ts65Dn presented an enlargement of the ventricles, which was previously associated with a decrease of cortical neurogenesis in the brains of the Ts1Cje and Ts2Cje mouse models (33).

Comparative genome wide expression profiling in the mouse hippocampus revealed that the entire dysregulation cannot be attributed to a single gene or region. The overall effect results from a complex interplay between certain trisomic overexpressed genes and other genes spread along the genome, evidenced by the fact that the majority of DEGs were not Hsa21 genes (S3 table). Additionally, we identified 34 trisomic DEGs (TEGs) with regulatory activity (Transcription factors, chromatin modellers, etc.) as *Mir99a, Usp16, Erg* or *Rcan1* that may be involved in the changed regulatory landscape of the models. Indeed, *RCAN1* and *USP16* were found to be upregulated in human brain datasets (cerebrum and cerebellum)(74, 75) and USP16 was also found as a DEG upregulated in heart and adult fibroblasts while MIR99A was found upregulated in adult fibroblasts (51). Nevertheless, the expression of DEGs was strongly correlated and conserved in Mmu16 based DS models. This is similar to the behavioural results obtained where related phenotypes were found in models carrying correlated partial duplications. Unexpectedly, Dp1Yey DEGs correlation was closer to Tg(*Dyrk1a*) than to Ts65Dn (42% against 25%) and there was a negative correlation between Dp3Yah & Dp5Yah (22%) and Tg(*Dyrk1a*) & Dp5Yah (13%). These correlations point to different gene dysregulations in these models and to the existence of epistasis with several regulatory trisomic genes countering the effect of genes in other trisomic regions.

We carried out an in depth functional annotation analysis to characterise 10 major meta-pathways with ribosome and mitochondrial functions, transcription & epigenomic regulation, and the synapse function categories highly affected. We also found a strong upregulation of genes involved in the Interferon beta pathway (S13 FigB) as some interferon receptors were found upregulated in Mmu16 DS models. As such *Ifnar2* and *Il10rb* were found upregulated in all the mice lines (except Tg(*Dyrk1a*)), pointing to a potentially critical role in interferon pathway dysregulation. The same phenomenon was observed with other genes like *Irgn1,Ifit1, Ifit2* or *Ndufa13*. This upregulation of the interferon beta pathways was previously reported in the Ts1Cje mouse model (76, 77) and linked to a possible increase of activity of the Jak-Stat signaling pathway, recorded here by the up regulation of *Stat1*.

The expression of genes involved in long-term synaptic potentiation (LTP) and synaptic plasticity were decreased in Dp1Yey, Dp5Yah, Dp1Rhr, Dp5/Dp1 models respectively, corroborating previous reports on different DS mouse models and in vitro studies. The only upregulated pathways were the myelin sheath and SNARE, both found in Dp1Yey and Tg(Dyrk1A) models. Interestingly, models carrying the Dp1Rhr region duplication showed dysregulation mainly in synapse transmission, plasticity and LTP, while models carrying the Dp5Yah region duplication showed dysregulation associated with genes involved in stemness and differentiation. Together, the models with both Dp5Yah & Dp1Rhr duplicated regions were involved in post-synapse modulation and transmission. Thus, there seems both region-specific effects and and region dependent effects.

Moreover, a high intermodel inter-pathway connectivity approach showed 6 major subnetwork biological cascades controlled by DYRK1A, GSK3B, NPY, SNARE complex, RHOA and NPAS4, which play a crucial role in the brain function. DYRK1A is a well recognized driver of DS phenotypes and the target of several therapeutic approaches (25, 39) which also interacts with GSK3B and NPY in DS models (25, 78–82). Thus, we were pleased to detect these 3 pathways and surprised to notice how closely interconnected those subnetworks were. We also described new pathways with SNARE, RHOA and NPAS4 in models based on Mmu16. Interestingly, SNARE complex proteins were also found modified in a DS model for the region homologous to Hsa21 on Mmu17 (60). RHOA is a member of the RHO GTPase involved in several intellectual disabilities that affect dendritic structure in adult neurons (83–86), a phenotype also described in certain DS models (87–89) or linked to DYRK1A (90, 91). Misregulation of *Npas4* and IEGs was found in various DS mouse models and can induce abnormal regulation when activated in specific biological processes, such as cognition in the Ts65Dn model (92). Furthermore, the network analyses highlighted NPAS4 as a potential modulator of synaptic dysfunction via well connected interactors. NPAS4 could affect the main altered biological cascades in addition to the GABA and NMDA receptors involved in the modulation of the excitatory / inhibitory balance of the brain (62).

The betweenness centrality index value is used to measure the interconnectivity of the network and showed that RHOA, DYRK1A, GSK3B, and their interactors were more closely knitted together and populated the central part of the network while SNARE, NPAS4 and NPY with their first- and second-layer interactors were more in the periphery of the network. This strong interconnectivity is of interest for two reasons: it makes the full network highly sensitive to targeted attacks against these proteins while, on the contrary, the network is robust against such attacks if they do not target several proteins simultaneously, for example during a drug trial. Thus, studying further closely connected altered genes and understanding their interactions could provide novel insights into the possible molecular mechanisms explaining why so many compounds, including DYRK1A specific kinase inhibitors, are capable of restoring learning and memory in DS models (39, 93–95). Additionally, these nodes show a large number of connections. Indeed, using the betweenness index, these nodes can be seen to orchestrate the network and their interactors occupy its centre, illustrating their extreme importance for its stability in terms of network theory. Moreover, similar to the observation in DS people, where the affectation/severity of gene dysfunction varies from one individual to another, we propose here that different DS mouse models show different changes in the 6 signalling interconnected cascades affecting brain dysfunction, leading to similar behavioural phenotypes. Our observation may support the developmental instability hypothesis, which postulates that the non specific triplication of a relatively small number of genes causes genetic imbalance with a significant impact on global gene expression. This hypothesis is in agreement with the study of rare individuals, carrying Hsa21 duplications, who displayed intellectual disabilities (5, 7) and should be taken into account when therapeutic assays are planned. We suggest that preclinical observation in one partial trisomic mouse model should be replicated in more genetically complex models to test potential genetic influences (39, 60, 96). This is probably the limit of the model, since although behaviour and memory mechanisms are common between mice and humans, the complexity of the model is lower. Conducting the same studies in more complex animal models carrying all the trisomic genes homologous to Has21, would definitely permit better deciphering of genes having an impact on cognitive behaviour.

Taking advantage of DS mouse models, we investigated behaviour and cognition, brain morphology and hippocampal gene expression in a standardized and controlled manner. Our results with the partial duplication of the Mmu16 region homologous to Hsa21 are in agreement with human genetic analysis (4–10) and showed how multiple genetic interactions between different regions of chromosome 21 contribute towards altering the outcome of the behavioural, morphological and molecular/pathway phenotypes. We are now faced with the challenge of carefully dissecting all these genetic interactions. Nonetheless, we found that overlapping DS models show convergence in the biological cascades altered, observed via building protein-protein interaction and regulatory networks, and centred on 6 main hubs: DYRK1A, GSK3β, NPY, SNARE, RHOA and NPAS4. We propose to name these hubs the center of the DS biological cascade. Some of them have already been described as altered in certain DS models, and we validated two additional ones, RHOA and NPAS4. Thus, we have built a novel vision of existing altered gene-gene crosstalk and molecular mechanisms, with 6 specific highly interconnected DS hubs in mouse models. They may well prove essential in improving our understanding of DS neurobiology and making progress in therapy development.

## Material and Methods

### Mouse lines

The duplications of different Mmu 16 regions (Dp(16*Lipi*-*Zbtb21*)1Yey (or Dp1Yey), the Dp(16*Hspa13*-*App*)3Yah (noted Dp3Yah), the Dp(16*Cbr1*-*Fam3b*)1Rhr (or Dp1Rhr)) and BAC transgenic mice for *Dyrk1a* (Tg(*Dyrk1a*)) models were described previously (19, 20, 24, 38). The genetic background of the DS lines carrying each duplication was pushed toward the C57BL/6J (B6J) genetic background for more than 7 generations of backcrossing. The only exception was the trisomic Ts65Dn (Ts(17^16^)65Dn) mice, initially obtained from the Jax, which were kept on a F1 B6C3B genetic background (with the C3B line as a C_3_H/HeH congenic line for the BALB/c allele at the *Pde6b* locus (99)). The Dp(16*Cyyr1*-*Clic6*)5Yah (noted Dp5Yah) was generated by the *in vivo* TAMERE technology inserting loxP sites in *App* and *Runx1* (see Supplementary information). In the Dp3Yah and Dp5Yah models, only 2 complete copies of *App* and *Runx1* genes were expressed. The Dp5Yah line was crossed with the Dp1Rhr line in order to generate Dp5Yah/Dp1Rhr (also noted Dp5/Dp1) compound transheterozygote carrying a similar trisomic Mmu16 gene content to that of the Ts65Dn. Indeed, only 15 Hsa21 homologous genes (*Mrpl39, Jam2, Atp5j, Gabpa, App, Cyyr1, Runx1, Setd4, Mx2, Tmprss2, Ripk4, Prdm15, C2cd2 and Zbpb21*) out of 174 are not in 3 copies in Dp5/Dp1 compared to Ts65Dn. In addition 46 protein-coding genes located on the Mmu17 centromeric region in the Ts65Dn minichromosome (13, 16) are not trisomic in the Dp5/Dp1. The Dp5Yah model was also combined with Tg(*Dyrk1a*) by crossing Dp5Yah/+ and Tg(*Dyrk1a*)/0 animals and generating the four genotypes (Dp1Yah, Dp5Yah, Tg(*Dyrk1*a) and [Dp5Yah; Tg(*Dyrk1a*)] noted here Dp5-Tg), to test specific interaction between Dp5Yah and *Dyrk1a* overdosage.

All the lines were maintained under specific pathogen-free (SPF) conditions and were treated in compliance with the animal welfare policies of the French Ministry of Agriculture (law 87 848), and the phenotyping procedures were approved by our local ethical committee (Com’Eth, no. 17, APAFIS no. 2012-069).

### Behaviour pipeline

A series of behavioural experiments were conducted in male mice with a age-range starting at 2.5 up to 7 months for the last test, as described in supplementary information. The tests were administered in the following order: Y-maze, open field, novel object recognition (24h), Morris water maze and fear conditioning (contextual and cue). Behavioural experimenters were blinded to the genetic status of the animals. Separate groups of animals were composed for each line (as indicated in S1 table). Several mouse models found defective for the NOR performed with 24h of retention memory were also tested after 1h of retention. The Dp5Yah crossed with Tg(*Dyrk1a*) was tested for Y-maze and NOR at 24h. All the standard operating procedures for behavioural phenotyping have been already described (96, 100–103) and are detailed in the supplementary information.

### Magnetic Resonance Imaging

A dedicated cohort of animals at the age 102 +/− 7 days was anesthetized and perfused with 30 ml of room temperature 1X Phosphate Buffer Saline (PBS) complemented with 10% (% w/v) heparine and 2mM of ProHance Gadoteridol (Bracco Imaging, Courcouronnes, France) followed by 30ml of 4% PFA complemented with 2mM of ProHance Gadoteridol. Then the brain structure was dissected and kept in PFA 4% 2mM ProHance overnight at 4°C. The next day, each specimen was transferred into 1X PBS 2mM ProHance until imaging.

Just prior to imaging, the brains were removed from the fixative and placed into plastic tubes (internal diameter 1 cm, volume 13 mL) filled with a proton-free susceptibility-matching fluid (Fluorinert® FC-770, Sigma-Aldrich, St. Louis, MO). Images of excised brains were acquired on a 7T BioSpec animal MRI system (Bruker Biospin MRI GmbH, Ettlingen, Germany). Images were reconstructed using ParaVision 6.0. An actively decoupled quadrature-mode mouse brain surface coil was used for signal reception and a 72-mm birdcage coil was used for transmission, both supplied by Bruker. The first protocol consisted of a 3D T2-weighted rapid-acquisition with relaxation enhancement (RARE). The second imaging protocol consisted of a 3D T2*-weighted Fast Low Angle (FLASH) sequence. The image matrix for both sequences was 195 x 140 x 90 over a field of view 19.5 x 14.0 x 9.0 mm^3^ yielding an isotropic resolution of 100 μm and treated and analysed for anatomical parameters, as detailed in the supplementary information.

### Gene expression assay

Hippocampuses were isolated from DS trisomic models and their littermate controls (N = 5 per group) and flash frozen in liquid nitrogen. Total RNA was prepared using the RNA extraction kit (Qiagen, Venlo, Netherlands) according to the manufacturer’s instructions. Sample quality was checked using an Agilent 2100 Bioanalyzer (Agilent Technologies, Santa Clara, California, USA). Gene expression analysis was carried out using GeneChip® Mouse Gene 1.0 ST arrays (Affymetrix, Santa Clara, CA). All the procedures and analyses are detailed in supplementary information.

### Availability of data and materials

All the mouse lines are available in the Jax or the EMMA/Infrafrontier repository. Raw microarray data and re-analysed data have been deposited in GEO (Accession No. GSE149470).

### Bioinformatic analysis

The gene expression profile of the mouse hippocampi isolated from Dp1Yey, Dp3Yah, Ts65Dn, Dp5/Dp1, Dp5Yah, Dp1Rhr and Tg(*Dyrk1a*) trisomic mouse models was analysed with a specific bioinformatics pipeline and controlled for quality prior to and after data pre-processing and normalization (see supplementary information in the detailed material and methods section). The differentially expressed genes (DEGs) were identified using a method based on fold change rank ordering statistics (FCROS)(104). In the FCROS method, k pairs of test/control samples are used to compute fold changes (FC). For each pair of test/control samples, the FCs obtained for all genes are ranked in increasing order. Resulting ranks are associated with genes. Then, the k ranks of each gene are used to calculate a statistic and resulting probability (f-value) used to identify the DEGs after fixing the error level at 5% False Discovery Rate (FDR).

We performed the functional differential analysis using GAGE (105) and grouped all the pathways into 10 functional categories (noted meta-pathways). Functional intermodel meta-pathway connectivity was studied by identifying the genes shared between pathways and models inside the same meta-pathway. Then, to assess gene connectivity we built a minimum fully connected protein-protein interaction (PPI) network (noted MinPPINet) of genes known to be involved in synaptic function as they were associated with synaptic pathways via GO (106) and KEGG databases (107). Furthermore, regulatory information was added to build the final RegPPINet. We used the betweenness centrality analysis to identify hubs, keys for maintaining the network communication flow. The relevance of the connecting nodes was further predicted by the machine learning algorithm Quack (108). Finally, we computed 100000 random networks with a similar degree, to assess if the likelihood of observing such connectivity in the DS network was more than one can expect by chance using statnet and sna R packages (https://cran.r-project.org/web/packages/statnet/index.html; https://cran.r-project.org/web/packages/sna/index.html). The full list of R packages used can be found in table S7.

### Western blot

The expression levels of the RHOA protein and Myosin Light Chain phosphorylation by the Myosin Light Chain Kinase part of the RHOA pathway were analysed using Western Blot in 5 animals Dp1Yey and 5 control (wt) littermates (See Fig 4G, Supplementary fig 16, and supplementary information). We used the following primary antibodies: anti-RHOA (2117, Cell Signaling, USA, 1:1.000), anti-pMLC (Thr18/Ser19 #3674, Cell signalling, Boston, MA, USA, 1:1.000) and mouse monoclonal Anti-β-Actin−Peroxidase antibody (A3854 Sigma, 1:150.000); and HRP conjugated Goat anti-Rabbit IgG secondary antibody (A16096, Invitrogen, France). Protein signals were visualized with Amersham™ Imager 600 and were quantified using ImageJ and statistical analysis using Sigma Plot. The relative amount of RHOA and p-MLC proteins was calculated as the ratio of the signal detected for each protein of interest compared to the β-actin signal detected and normalized by the mean signal of the wt samples.

### Visual stimulation

Mice raised in a standard light cycle were housed in constant darkness for two weeks. Then, animals in the light-exposed condition group, were consecutively exposed to light for 0, 1, 3, and 7.5 h before being sacrificed. The animals belonging to the dark-housed condition group were sacrificed in the dark. After euthanasia, their eyes were enucleated before visual cortex dissection in the light and flash frozen in liquid nitrogen. cDNA and quantitative PCR were performed as indicated in supplementary information. The Ct values were transformed to quantities by using the comparative Ct method. Hence, all data were expressed relative to the expression of the most expressed gene. These relative expression levels, were normalized with Genorm by keeping the more stable reference genes (109). To calculate fold-induction, the relative quantity of gene expression at each time point was divided by the mean of the relative level of gene expression of dark-housed mice for the corresponding genotype. The mean and standard error were calculated at each time point from these fold-induction values.

### Statistical analysis

All data are expressed as mean group value ± standard error of the mean (SEM) or as box plots with the median and quartiles. For each data set, we analysed if the data were normally distributed using the Shapiro–Wilk test and Quantile-Quantile plots (S2 Fig) and the homogeneity of variances by the Brown-Forsy test. Differences between groups were inferred by one-way ANOVA (Open field) and ANOVA for repeated measures, or we performed the Kruskal-Wallis non-parametric test in the case of datasets where the assumptions of normality or homogeneity of variances were not fulfilled. The post hoc tests (Fisher LSD Method) were conducted only if the F parameter in ANOVA achieved a level of 0.05. All the behavioral analysis results can be found in table S1. For the MRI data, intergroup comparisons on region-based data were conducted on the normalized volumes (i.e. ratio between the volume of the structure and the whole brain volume) of each segmented structure using the Student t-test while correcting by multiple testing setting up an FDR correction.

## Acknowledgements

We would like to thank members of the research group, of the IGBMC laboratory and of the ICS. We are grateful to the IGBMC microarray and Sequencing platform and particularly C. Thibault-Carpentier, for providing us access to microarray. We extend our thanks to the animal care-takers of the ICS who are in charge of the mice wellness. We also thank F.Riet, C. Mittelhaeuser, A. Lux and V. Alunni and D. Dembele for expert technical assistance and useful discussion.We would like to aknowledge that Maria del Mar Muñiz Moreno was an IGBMC International PhD Programme fellow supported by LabEx INRT funds [ANR-10-LABX-0030-INRT].

This work has been supported by the National Centre for Scientific Research (CNRS), the French National Institute of Health and Medical Research (INSERM), the University of Strasbourg (Unistra), the French state funds through the “Agence Nationale de la Recherche” under the frame programme Investissements d’Avenir [ANR-10-IDEX-0002-02, ANR-10-LABX-0030-INRT, ANR-10-INBS-07 PHENOMIN to YH]. The funders had no role in study design, data collection and analysis, decision to publish, or preparation of the manuscript.

## Conflicts of Interest Statement

The authors declare no conflict of interest

## Notes

### Competing Interest Statement

The authors have declared no competing interest.

### Summary of Updates

We improve the manuscript, change some figure to facilitate the understanding of the models used in the study

## References

1. Lejeune, J., Turpin, R. and Gautier, M. (1959) [Mongolism; a chromosomal disease (trisomy).]. Bull Acad Natl Med, 143, 256–265.

2. Antonarakis, S.E., Lyle, R., Dermitzakis, E.T., Reymond, A. and Deutsch, S. (2004) Chromosome 21 and down syndrome: from genomics to pathophysiology. Nat Rev Genet, 5, 725–738.

3. Oliver, T.R., Feingold, E., Yu, K., Cheung, V., Tinker, S., Yadav-Shah, M., Masse, N. and Sherman, S.L. (2008) New insights into human nondisjunction of chromosome 21 in oocytes. PLoS Genet, 4, e1000033.

4. McCormick, M.K., Schinzel, A., Petersen, M.B., Stetten, G., Driscoll, D.J., Cantu, E.S., Tranebjaerg, L., Mikkelsen, M., Watkins, P.C. and Antonarakis, S.E. (1989) Molecular genetic appoach to the characterization of the &quot;Down syndrome region&quot; of chromosome 21. Genomics, 5, 325–331.

5. Korbel, J.O., Tirosh-Wagner, T., Urban, A.E., Chen, X.N., Kasowski, M., Dai, L., Grubert, F., Erdman, C., Gao, M.C., Lange, K. et al. (2009) The genetic architecture of Down syndrome phenotypes revealed by high-resolution analysis of human segmental trisomies. Proceedings of the National Academy of Sciences of the United States of America, 106, 12031–12036.

6. Korenberg, J.R. (1990) Molecular mapping of the Down syndrome phenotype. Prog Clin Biol Res, 360, 105–115.

7. Lyle, R., Béna, F., Gagos, S., Gehrig, C., Lopez, G., Schinzel, A., Lespinasse, J., Bottani, A., Dahoun, S., Taine, L. et al. (2009) Genotype-phenotype correlations in Down syndrome identified by array CGH in 30 cases of partial trisomy and partial monosomy chromosome 21. Eur J Hum Genet, 17, 454–466.

8. Delabar, J.M., Theophile, D., Rahmani, Z., Chettouh, Z., Blouin, J.L., Prieur, M., Noel, B. and Sinet, P.M. (1993) Molecular mapping of twenty-four features of Down syndrome on chromosome 21. Eur J Hum Genet, 1, 114–124.

9. Korenberg, J.R., Chen, X.N., Schipper, R., Sun, Z., Gonsky, R., Gerwehr, S., Carpenter, N., Daumer, C., Dignan, P. and Disteche, C. (1994) Down syndrome phenotypes: the consequences of chromosomal imbalance. Proc Natl Acad Sci U S A, 91, 4997–5001.

10. Rahmani, Z., Blouin, J.L., Creau-Goldberg, N., Watkins, P.C., Mattei, J.F., Poissonnier, M., Prieur, M., Chettouh, Z., Nicole, A. and Aurias, A. (1989) Critical role of the D21S55 region on chromosome 21 in the pathogenesis of Down syndrome. Proc Natl Acad Sci U S A, 86, 5958–5962.

11. Herault, Y., Delabar, J.M., Fisher, E.M.C., Tybulewicz, V.L.J., Yu, E. and Brault, V. (2017) Rodent models in Down syndrome research: impact and future opportunities. Disease Models & Mechanisms, 10, 1165–1186.

12. Gupta, M., Dhanasekaran, A.R. and Gardiner, K.J. (2016) Mouse models of Down syndrome: gene content and consequences. Mammalian Genome, 27, 538–555.

13. Muñiz Moreno, M.D.M., Brault, V., Birling, M.C., Pavlovic, G. and Herault, Y. (2020) Modeling Down syndrome in animals from the early stage to the 4.0 models and next. Prog Brain Res, 251, 91–143.

14. Herault, Y., Duchon, A., Velot, E., Marechal, D., Brault, V., Dierssen, M. and DeLaTorre, R. (2012) The in vivo Down syndrome genomic library in mouse. Down Syndrome: From Understanding the Neurobiology To Therapy, 197, 169–197.

15. Reeves, R.H., Irving, N.G., Moran, T.H., Wohn, A., Kitt, C., Sisodia, S.S., Schmidt, C., Bronson, R.T. and Davisson, M.T. (1995) A MOUSE MODEL FOR DOWN-SYNDROME EXHIBITS LEARNING AND BEHAVIOR DEFICITS. Nature Genetics, 11, 177–184.

16. Duchon, A., Raveau, M., Chevalier, C., Nalesso, V., Sharp, A.J. and Herault, Y. (2011) Identification of the translocation breakpoints in the Ts65Dn and Ts1Cje mouse lines: relevance for modeling down syndrome. Mammalian Genome, 22, 674–684.

17. Reinholdt, L.G., Ding, Y.M., Gilbert, G.T., Czechanski, A., Solzak, J.P., Roper, R.J., Johnson, M.T., Donahue, L.R., Lutz, C. and Davisson, M.T. (2011) Molecular characterization of the translocation breakpoints in the Down syndrome mouse model Ts65Dn. Mammalian Genome, 22, 685–691.

18. Davisson, M.T., Schmidt, C., Reeves, R.H., Irving, N.G., Akeson, E.C., Harris, B.S. and Bronson, R.T. (1993) Segmental trisomy as a mouse model for Down syndrome. Prog Clin Biol Res, 384, 117–133.

19. Li, Z., Yu, T., Morishima, M., Pao, A., LaDuca, J., Conroy, J., Nowak, N., Matsui, S.-I., Shiraishi, I. and Yu, Y.E. (2007) Duplication of the entire 22.9 Mb human chromosome 21 syntenic region on mouse chromosome 16 causes cardiovascular and gastrointestinal abnormalities. Hum Mol Genet, 16, 1359–1366.

20. Olson, L.E., Richtsmeier, J.T., Leszl, J. and Reeves, R.H. (2004) A chromosome 21 critical region does not cause specific Down syndrome phenotypes. Science, 306, 687–690.

21. Olson, L.E., Roper, R.J., Baxter, L.L., Carlson, E.J., Epstein, C.J. and Reeves, R.H. (2004) Down syndrome mouse models Ts65Dn, Ts1Cje, and Ms1Cje/Ts65Dn exhibit variable severity of cerebellar phenotypes. Dev Dyn, 230, 581–589.

22. Olson, L.E., Roper, R.J., Sengstaken, C.L., Peterson, E.A., Aquino, V., Galdzicki, Z., Siarey, R., Pletnikov, M., Moran, T.H. and Reeves, R.H. (2007) Trisomy for the Down syndrome ‘critical region’ is necessary but not sufficient for brain phenotypes of trisomic mice. Hum Mol Genet, 16, 774–782.

23. Belichenko, N.P., Belichenko, P.V., Kleschevnikov, A.M., Salehi, A., Reeves, R.H. and Mobley, W.C. (2009) The “Down syndrome critical region” is sufficient in the mouse model to confer behavioral, neurophysiological, and synaptic phenotypes characteristic of Down syndrome. J Neurosci, 29, 5938–5948.

24. Guedj, F., Pereira, P.L., Najas, S., Barallobre, M.J., Chabert, C., Souchet, B., Sebrie, C., Verney, C., Herault, Y., Arbones, M. et al. (2012) DYRK1A: a master regulatory protein controlling brain growth. Neurobiol Dis, 46, 190–203.

25. Duchon, A. and Herault, Y. (2016) DYRK1A, a Dosage-Sensitive Gene Involved in Neurodevelopmental Disorders, Is a Target for Drug Development in Down Syndrome. Frontiers in Behavioral Neuroscience, 10.

26. De la Torre, R., De Sola, S., Pons, M., Duchon, A., Martinez de Lagran, M., Farre, M., Fito, M., Benejam, B., Langohr, K., Rodriguez, J. et al. (2014) Epigallocatechin-3-gallate, a DYRK1A inhibitor, rescues cognitive deficits in Down syndrome mouse models and in humans. Molecular Nutrition &amp; Food Research, 58, 278–288.

27. Altafaj, X., Martin, E.D., Ortiz-Abalia, J., Valderrama, A., Lao-Peregrin, C., Dierssen, M. and Fillat, C. (2013) Normalization of Dyrk1A expression by AAV2/1-shDyrk1A attenuates hippocampal-dependent defects in the Ts65Dn mouse model of Down syndrome. Neurobiology of Disease, 52, 117–127.

28. Garcia-Cerro, S., Martinez, P., Vidal, V., Corrales, A., Florez, J., Vidal, R., Rueda, N., Arbones, M.L. and Martinez-Cue, C. (2014) Overexpression of Dyrk1A Is Implicated in Several Cognitive, Electrophysiological and Neuromorphological Alterations Found in a Mouse Model of Down Syndrome. Plos One, 9.

29. Pennington, B.F., Moon, J., Edgin, J., Stedron, J. and Nadel, L. (2003) The neuropsychology of Down syndrome: evidence for hippocampal dysfunction. Child Dev, 74, 75–93.

30. Nadel, L. (2003) Down’s syndrome: a genetic disorder in biobehavioral perspective. Genes Brain Behav, 2, 156–166.

31. Brown, S.D.M., Hancock, J.M. and Gates, H. (2006) Understanding mammalian genetic systems: The challenge of phenotyping in the mouse. Plos Genetics, 2, 1131–1137.

32. Mandillo, M., Tucci, V., Holter, S.M., Meziane, H., Al Banchaabouchi, M., Kallnik, M., Lad, H.V., Nolan, P.M., Ouagazzal, A.M., Coghill, E.L. et al. (2010) Reliability, robustness, and reproducibility in mouse behavioral phenotyping: a cross-laboratory study. (vol 34, pg 243, 2008). Physiological Genomics, 40, 217–217.

33. Ishihara, K., Amano, K., Takaki, E., Shimohata, A., Sago, H., Epstein, C.J. and Yamakawa, K. (2010) Enlarged Brain Ventricles and Impaired Neurogenesis in the Ts1Cje and Ts2Cje Mouse Models of Down Syndrome. Cerebral Cortex, 20, 1131–1143.

34. Raveau, M., Nakahari, T., Asada, S., Ishihara, K., Amano, K., Shimohata, A., Sago, H. and Yamakawa, K. (2017) Brain ventriculomegaly in Down syndrome mice is caused by Pcp4 dose-dependent cilia dysfunction. Human Molecular Genetics, 26, 923–931.

35. Guedj, F., Pereira, P.L., Najas, S., Barallobre, M.J., Chabert, C., Souchet, B., Sebrie, C., Verney, C., Herault, Y., Arbones, M. et al. (2012) DYRK1A: A master regulatory protein controlling brain growth. Neurobiology of Disease, 46, 190–203.

36. Guedj, F., Sebrie, C., Rivals, I., Ledru, A., Paly, E., Bizot, J.C., Smith, D., Rubin, E., Gillet, B., Arbones, M. et al. (2009) Green Tea Polyphenols Rescue of Brain Defects Induced by Overexpression of DYRK1A. Plos One, 4, 8.

37. Gardiner, K., Fortna, A., Bechtel, L. and Davisson, M.T. (2003) Mouse models of Down syndrome: how useful can they be? Comparison of the gene content of human chromosome 21 with orthologous mouse genomic regions. Gene, 318, 137–147.

38. Brault, V., Duchon, A., Romestaing, C., Sahun, I., Pothion, S., Karout, M., Borel, C., Dembele, D., Bizot, J.C., Messaddeq, N. et al. (2015) Opposite phenotypes of muscle strength and locomotor function in mouse models of partial trisomy and monosomy 21 for the proximal Hspa13-App region. PLoS Genet, 11, e1005062.

39. Nguyen, T.L., Duchon, A., Manousopoulou, A., Loaëc, N., Villiers, B., Pani, G., Karatas, M., Mechling, A.E., Harsan, L.A., Limanton, E. et al. (2018) Correction of cognitive deficits in mouse models of Down syndrome by a pharmacological inhibitor of DYRK1A. Dis Model Mech, 11.

40. Faizi, M., Bader, P.L., Tun, C., Encarnacion, A., Kleschevnikov, A., Belichenko, P., Saw, N., Priestley, M., Tsien, R.W., Mobley, W.C. et al. (2011) Comprehensive behavioral phenotyping of Ts65Dn mouse model of Down Syndrome: Activation of pradrenergic receptor by xamoterol as a potential cognitive enhancer. Neurobiology of Disease, 43, 397–413.

41. Souchet, B., Guedj, F., Sahun, I., Duchon, A., Daubigney, F., Badel, A., Yanagawa, Y., Barallobre, M.J., Dierssen, M., Yu, E. et al. (2014) Excitation/inhibition balance and learning are modified by Dyrk1a gene dosage. Neurobiology of Disease, 69, 65–75.

42. Chang, P., Bush, D., Schorge, S., Good, M., Canonica, T., Shing, N., Noy, S., Wiseman, F.K., Burgess, N., Tybulewicz, V.L.J. et al. (2020) Altered Hippocampal-Prefrontal Neural Dynamics in Mouse Models of Down Syndrome. Cell Rep, 30, 1152–1163.e1154.

43. Escorihuela, R.M., Fernández-Teruel, A., Vallina, I.F., Baamonde, C., Lumbreras, M.A., Dierssen, M., Tobeña, A. and Flórez, J. (1995) A behavioral assessment of Ts65Dn mice: a putative Down syndrome model. Neurosci Lett, 199, 143–146.

44. Martinez-Cue, C., Baamonde, C., Lumbreras, M., Paz, J., Davisson, M.T., Schmidt, C., Dierssen, M. and Florez, J. (2002) Differential effects of environmental enrichment on behavior and learning of male and female Ts65Dn mice, a model for Down syndrome. Behavioural Brain Research, 134, 185–200.

45. Martinez-Cue, C., Rueda, N., Garcia, E., Davisson, M.T., Schmidt, C. and Florez, J. (2005) Behavioral, cognitive and biochemical responses to different environmental conditions in male Ts65Dn mice, a model of Down syndrome. Behavioural Brain Research, 163, 174–185.

46. Moran, T.H., Capone, G.T., Knipp, S., Davisson, M.T., Reeves, R.H. and Gearhart, J.D. (2002) The effects of piracetam on cognitive performance in a mouse model of Down’s syndrome. Physiol Behav, 77, 403–409.

47. Rueda, N., Florez, J. and Martinez-Cue, C. (2008) Effects of chronic administration of SGS-111 during adulthood and during the pre- and post-natal periods on the cognitive deficits of Ts65Dn mice, a model of Down syndrome. Behavioural Brain Research, 188, 355–367.

48. Rueda, N., Florez, J. and Martinez-Cue, C. (2008) Chronic pentylenetetrazole but not donepezil treatment rescues spatial cognition in Ts65Dn mice, a model for Down syndrome. Neuroscience Letters, 433, 22–27.

49. Seo, H. and Isacson, O. (2005) Abnormal APP, cholinergic and cognitive function in Ts65Dn Down’s model mice. Exp Neurol, 193, 469–480.

50. Lockstone, H.E., Harris, L.W., Swatton, J.E., Wayland, M.T., Holland, A.J. and Bahn, S. (2007) Gene expression profiling in the adult Down syndrome brain. Genomics, 90, 647–660.

51. Prandini, P., Deutsch, S., Lyle, R., Gagnebin, M., Vivier, C.D., Delorenzi, M., Gehrig, C., Descombes, P., Sherman, S., Bricarelli, F.D. et al. (2007) Natural gene-expression variation in Down syndrome modulates the outcome of gene-dosage imbalance. American Journal of Human Genetics, 81, 252–263.

52. Ait Yahya-Graison, E., Aubert, J., Dauphinot, L., Rivals, I., Prieur, M., Golfier, G., Rossier, J., Personnaz, L., Creau, N., Blehaut, H. et al. (2007) Classification of human chromosome 21 gene-expression variations in Down syndrome: impact on disease phenotypes. American journal of human genetics, 81, 475–491.

53. Laffaire, J., Rivals, I., Dauphinot, L., Pasteau, F., Wehrle, R., Larrat, B., Vitalis, T., Moldrich, R.X., Rossier, J., Sinkus, R. et al. (2009) Gene expression signature of cerebellar hypoplasia in a mouse model of Down syndrome during postnatal development. Bmc Genomics, 10.

54. Sultan, M., Piccini, I., Balzereit, D., Herwig, R., Saran, N.G., Lehrach, H., Reeves, R.H. and Yaspo, M.L. (2007) Gene expression variation in ‘Down syndrome’ mice allows to prioritize candidate genes. Genome Biol, 8, R91.

55. Mao, R., Zielke, C.L., Zielke, H.R. and Pevsner, J. (2003) Global up-regulation of chromosome 21 gene expression in the developing Down syndrome brain. Genomics, 81, 457–467.

56. Olmos-Serrano, J.L., Kang, H.J., Tyler, W.A., Silbereis, J.C., Cheng, F., Zhu, Y., Pletikos, M., Jankovic-Rapan, L., Cramer, N.P., Galdzicki, Z. et al. (2016) Down Syndrome Developmental Brain Transcriptome Reveals Defective Oligodendrocyte Differentiation and Myelination. Neuron, 89, 1208–1222.

57. Guedj, F., Pennings, J.L., Massingham, L.J., Wick, H.C., Siegel, A.E., Tantravahi, U. and Bianchi, D.W. (2016) An Integrated Human/Murine Transcriptome and Pathway Approach To Identify Prenatal Treatments For Down Syndrome. Sci Rep, 6, 32353.

58. Guedj, F., Pennings, J.L., Wick, H.C. and Bianchi, D.W. (2015) Analysis of adult cerebral cortex and hippocampus transcriptomes reveals unique molecular changes in the Ts1Cje mouse model of down syndrome. Brain Pathol, 25, 11–23.

59. Aziz, N.M., Guedj, F., Pennings, J.L.A., Olmos-Serrano, J.L., Siegel, A., Haydar, T.F. and Bianchi, D.W. (2018) Lifespan analysis of brain development, gene expression and behavioral phenotypes in the Ts1Cje, Ts65Dn and Dp(16)1/Yey mouse models of Down syndrome. Dis Model Mech, 11.

60. Marechal, D., Brault, V., Leon, A., Martin, D., Pereira, P.L., Loaëc, N., Birling, M.C., Friocourt, G., Blondel, M. and Herault, Y. (2019) Cbs overdosage is necessary and sufficient to induce cognitive phenotypes in mouse models of Down syndrome and interacts genetically with Dyrk1a. Hum Mol Genet.

61. Joazeiro, C.A.P. (2019) Mechanisms and functions of ribosome-associated protein quality control. Nature Reviews Molecular Cell Biology, 20, 368–383.

62. Mardinly, A.R., Spiegel, I., Patrizi, A., Centofante, E., Bazinet, J.E., Tzeng, C.P., Mandel-Brehm, C., Harmin, D.A., Adesnik, H., Fagiolini, M. et al. (2016) Sensory experience regulates cortical inhibition by inducing IGF1 in VIP neurons. Nature, 531, 371–375.

63. Spiegel, I., Mardinly, A.R., Gabel, H.W., Bazinet, J.E., Couch, C.H., Tzeng, C.P., Harmin, D.A. and Greenberg, M.E. (2014) Npas4 Regulates Excitatory-Inhibitory Balance within Neural Circuits through Cell-Type-Specific Gene Programs. Cell, 157, 1216–1229.

64. Smith, G.K., Kesner, R.P. and Korenberg, J.R. (2014) Dentate Gyrus Mediates Cognitive Function in the Ts65Dn/DnJ Mouse Model of Down Syndrome. Hippocampus, 24, 354–362.

65. Belichenko, P.V., Kleschevnikov, A.M., Becker, A., Wagner, G.E., Lysenko, L.V., Yu, Y.E. and Mobley, W.C. (2015) Down Syndrome Cognitive Phenotypes Modeled in Mice Trisomic for All HSA 21 Homologues.

66. Yu, T., Liu, C.H., Belichenko, P., Clapcote, S.J., Li, S.M., Pao, A.N., Kleschevnikov, A., Bechard, A.R., Asrar, S., Chen, R.Q. et al. (2010) Effects of individual segmental trisomies of human chromosome 21 syntenic regions on hippocampal long-term potentiation and cognitive behaviors in mice. Brain Research, 1366, 162–171.

67. Chakrabarti, L., Best, T.K., Cramer, N.P., Carney, R.S.E., Isaac, J.T.R., Galdzicki, Z. and Haydar, T.F. (2010) Olig1 and Olig2 triplication causes developmental brain defects in Down syndrome. Nature Neuroscience, 13, 927–U939.

68. Xu, R., Brawner, A.T., Li, S., Liu, J.J., Kim, H., Xue, H., Pang, Z.P., Kim, W.Y., Hart, R.P., Liu, Y. et al. (2019) OLIG2 Drives Abnormal Neurodevelopmental Phenotypes in Human iPSC-Based Organoid and Chimeric Mouse Models of Down Syndrome. Cell Stem Cell, 24, 908–926 e908.

69. Ermak, G., Cheadle, C., Becker, K.G., Harris, C.D. and Davies, K.J.A. (2004) DSCR1 (Adapt78) modulates expression of SOD1. Faseb Journal, 18, 62–69.

70. Voronov, S.V., Frere, S.G., Giovedi, S., Pollina, E.A., Borel, C., Zhang, H., Schmidt, C., Akeson, E.C., Wenk, M.R., Cimasoni, L. et al. (2008) Synaptojanin 1-linked phosphoinositide dyshomeostasis and cognitive deficits in mouse models of Down’s syndrome. Proceedings of the National Academy of Sciences of the United States of America, 105, 9415–9420.

71. Antonarakis, S.E. (2016) Down syndrome and the complexity of genome dosage imbalance. Nat Rev Genet.

72. Chen, C.-K., Bregere, C., Paluch, J., Lu, J.F., Dickman, D.K. and Chang, K.T. (2014) Activity-dependent facilitation of Synaptojanin and synaptic vesicle recycling by the Minibrain kinase. Nature Communications, 5.

73. Dunnett, S.B., Everitt, B.J. and Robbins, T.W. (1991) THE BASAL FOREBRAIN CORTICAL CHOLINERGIC SYSTEM - INTERPRETING THE FUNCTIONAL CONSEQUENCES OF EXCITOTOXIC LESIONS. Trends in Neurosciences, 14, 494–501.

74. Mao, R., Wang, X., Spitznagel, E.L., Jr., Frelin, L.P., Ting, J.C., Ding, H., Kim, J.W., Ruczinski, I., Downey, T.J. and Pevsner, J. (2005) Primary and secondary transcriptional effects in the developing human Down syndrome brain and heart. Genome Biol, 6, R107.

75. Lockstone, H.E., Harris, L.W., Swatton, J.E., Wayland, M.T., Holland, A.J. and Bahn, S. (2007) Gene expression profiling in the adult Down syndrome brain. Genomics, 90, 647–660.

76. Lee, H.C., Tan, K.L., Cheah, P.S. and Ling, K.H. (2016) Potential Role of JAK-STAT Signaling Pathway in the Neurogenic-to-Gliogenic Shift in Down Syndrome Brain. Neural Plast, 2016, 7434191.

77. Ling, K.H., Hewitt, C.A., Tan, K.L., Cheah, P.S., Vidyadaran, S., Lai, M.I., Lee, H.C., Simpson, K., Hyde, L., Pritchard, M.A. et al. (2014) Functional transcriptome analysis of the postnatal brain of the Ts1Cje mouse model for Down syndrome reveals global disruption of interferon-related molecular networks. BMC Genomics, 15, 624.

78. Woods, Y.L., Cohen, P., Becker, W., Jakes, R., Goedert, M., Wang, X. and Proud, C.G. (2001) The kinase DYRK phosphorylates protein-synthesis initiation factor eIF2Bepsilon at Ser539 and the microtubule-associated protein tau at Thr212: potential role for DYRK as a glycogen synthase kinase 3-priming kinase. Biochem J, 355, 609–615.

79. Hong, S.H., Lee, K.S., Kwak, S.J., Kim, A.K., Bai, H., Jung, M.S., Kwon, O.Y., Song, W.J., Tatar, M. and Yu, K. (2012) Minibrain/Dyrk1a regulates food intake through the Sir2-FOXO-sNPF/NPY pathway in Drosophila and mammals. PLoS Genet, 8, e1002857.

80. Hernández-González, S., Ballestín, R., López-Hidalgo, R., Gilabert-Juan, J., Blasco-Ibáñez, J.M., Crespo, C., Nácher, J. and Varea, E. (2015) Altered distribution of hippocampal interneurons in the murine Down Syndrome model Ts65Dn. Neurochem Res, 40, 151–164.

81. Shukkur, E.A., Shimohata, A., Akagi, T., Yu, W.X., Yamaguchi, M., Murayama, M., Chui, D., Takeuchi, T., Amano, K., Subramhanya, K.H. et al. (2006) Mitochondrial dysfunction and tau hyperphosphorylation in Ts1Cje, a mouse model for Down syndrome. Human Molecular Genetics, 15, 2752–2762.

82. King, M.K., Pardo, M., Cheng, Y.Y., Downey, K., Jope, R.S. and Beurel, E. (2014) Glycogen synthase kinase-3 inhibitors: Rescuers of cognitive impairments. Pharmacology & Therapeutics, 141, 1–12.

83. Khelfaoui, M., Pavlowsky, A., Powell, A.D., Valnegri, P., Cheong, K.W., Blandin, Y., Passafaro, M., Jefferys, J.G., Chelly, J. and Billuart, P. (2009) Inhibition of RhoA pathway rescues the endocytosis defects in Oligophrenin1 mouse model of mental retardation. Hum Mol Genet, 18, 2575–2583.

84. da Silva, J.S. and Dotti, C.G. (2002) Breaking the neuronal sphere: regulation of the actin cytoskeleton in neuritogenesis. Nat Rev Neurosci, 3, 694–704.

85. Ramakers, G.J.A. (2002) Rho proteins, mental retardation and the cellular basis of cognition. Trends in Neurosciences, 25, 191–199.

86. Billuart, P., Bienvenu, T., Ronce, N., des Portes, V., Vinet, M.C., Zemni, R., Carrié, A., Beldjord, C., Kahn, A., Moraine, C. et al. (1998) Oligophrenin 1 encodes a rho-GAP protein involved in X-linked mental retardation. Pathol Biol (Paris), 46, 678.

87. Belichenko, P.V., Kleschevnikov, A.M., Salehi, A., Epstein, C.J. and Mobley, W.C. (2007) Synaptic and cognitive abnormalities in mouse models of down syndrome: Exploring genotype-phenotype relationships. Journal of Comparative Neurology, 504, 329–345.

88. Belichenko, P.V., Masliah, E., Kleschevnikov, A.M., Villar, A.J., Epstein, C.J., Salehi, A. and Mobley, W.C. (2004) Synaptic structural abnormalities in the Ts65Dn mouse model of Down Syndrome. J Comp Neurol, 480, 281–298.

89. Haas, M.A., Bell, D., Slender, A., Lana-Elola, E., Watson-Scales, S., Fisher, E.M., Tybulewicz, V.L. and Guillemot, F. (2013) Alterations to dendritic spine morphology, but not dendrite patterning, of cortical projection neurons in Tc1 and Ts1Rhr mouse models of Down syndrome. PLoS One, 8, e78561.

90. Ori-McKenney, K.M., McKenney, R.J., Huang, H.H., Li, T., Meltzer, S., Jan, L.Y., Vale, R.D., Wiita, A.P. and Jan, Y.N. (2016) Phosphorylation of β-Tubulin by the Down Syndrome Kinase, Minibrain/DYRK1a, Regulates Microtubule Dynamics and Dendrite Morphogenesis. Neuron, 90, 551–563.

91. Thomazeau, A., Lassalle, O., Iafrati, J., Souchet, B., Guedj, F., Janel, N., Chavis, P., Delabar, J. and Manzoni, O.J. (2014) Prefrontal deficits in a murine model overexpressing the down syndrome candidate gene dyrk1a. J Neurosci, 34, 1138–1147.

92. Braudeau, J., Dauphinot, L., Duchon, A., Loistron, A., Dodd, R.H., Hérault, Y., Delatour, B. and Potier, M.C. (2011) Chronic Treatment with a Promnesiant GABA-A α5-Selective Inverse Agonist Increases Immediate Early Genes Expression during Memory Processing in Mice and Rectifies Their Expression Levels in a Down Syndrome Mouse Model. Adv Pharmacol Sci, 2011, 153218.

93. de la Torre, R., de Sola, S., Hernandez, G., Farre, M., Pujol, J., Rodriguez, J., Espadaler, J.M., Langohr, K., Cuenca-Royo, A., Principe, A. et al. (2016) Safety and efficacy of cognitive training plus epigallocatechin-3-gallate in young adults with Down’s syndrome (TESDAD): a double-blind, randomised, placebo-controlled, phase 2 trial. Lancet Neurology, 15, 801–810.

94. Nakano-Kobayashi, A., Awaya, T., Kii, I., Sumida, Y., Okuno, Y., Yoshida, S., Sumida, T., Inoue, H., Hosoya, T. and Hagiwara, M. (2017) Prenatal neurogenesis induction therapy normalizes brain structure and function in Down syndrome mice. Proc Natl Acad Sci U S A, 114, 10268–10273.

95. Herault, Y., Delabar, J.M., Fisher, E.M.C., Tybulewicz, V.L.J., Yu, E. and Brault, V. (2017) Rodent models in Down syndrome research: impact and future opportunities. Dis Model Mech, 10, 1165–1186.

96. Marechal, D., Lopes Pereira, P., Duchon, A. and Herault, Y. (2015) Dosage of the Abcg1-U2af1 region modifies locomotor and cognitive deficits observed in the Tc1 mouse model of Down syndrome. PLoS One, 10, e0115302.

97. Busti, I., Allegra, M., Spalletti, C., Panzi, C., Restani, L., Billuart, P. and Caleo, M. (2020) ROCK/PKA Inhibition Rescues Hippocampal Hyperexcitability and GABAergic Neuron Alterations in a Oligophrenin-1 Knock-Out Mouse Model of X-Linked Intellectual Disability. J Neurosci, 40, 2776–2788.

98. Meziane, H., Khelfaoui, M., Morello, N., Hiba, B., Calcagno, E., Reibel-Foisset, S., Selloum, M., Chelly, J., Humeau, Y., Riet, F. et al. (2016) Fasudil treatment in adult reverses behavioural changes and brain ventricular enlargement in Oligophrenin-1 mouse model of intellectual disability. Human Molecular Genetics, 25, 2314–2323.

99. Hoelter, S.M., Dalke, C., Kallnik, M., Becker, L., Horsch, M., Schrewe, A., Favor, J., Klopstock, T., Beckers, J., Ivandic, B. et al. (2008) “Sighted C3H” mice - a tool for analysing the influence of vision on mouse behaviour? Frontiers in Bioscience, 13, 5810–5823.

100. Dubos, A., Meziane, H., Iacono, G., Curie, A., Riet, F., Martin, C., Loaec, N., Birling, M.C., Selloum, M., Normand, E. et al. (2018) A new mouse model of ARX dup24 recapitulates the patients’ behavioural and fine motor alterations. Hum Mol Genet.

101. Arbogast, T., Iacono, G., Chevalier, C., Afinowi, N.O., Houbaert, X., van Eede, M.C., Laliberte, C., Birling, M.C., Linda, K., Meziane, H. et al. (2017) Mouse models of 17q21.31 microdeletion and microduplication syndromes highlight the importance of Kansl1 for cognition. PLoS Genet, 13, e1006886.

102. Ung, D.C., Iacono, G., Méziane, H., Blanchard, E., Papon, M.A., Selten, M., van Rhijn, J.R., Montjean, R., Rucci, J., Martin, S. et al. (2017) Ptchd1 deficiency induces excitatory synaptic and cognitive dysfunctions in mouse. Mol Psychiatry.

103. Arbogast, T., Ouagazzal, A.M., Chevalier, C., Kopanitsa, M., Afinowi, N., Migliavacca, E., Cowling, B.S., Birling, M.C., Champy, M.F., Reymond, A. et al. (2016) Reciprocal Effects on Neurocognitive and Metabolic Phenotypes in Mouse Models of 16p11.2 Deletion and Duplication Syndromes. PLoS Genet, 12, e1005709.

104. Dembele, D. and Kastner, P. (2014) Fold change rank ordering statistics: a new method for detecting differentially expressed genes. Bmc Bioinformatics, 15.

105. Luo, W., Friedman, M.S., Shedden, K., Hankenson, K.D. and Woolf, P.J. (2009) GAGE: generally applicable gene set enrichment for pathway analysis. BMC Bioinformatics, 10, 161.

106. Ashburner, M., Ball, C.A., Blake, J.A., Botstein, D., Butler, H., Cherry, J.M., Davis, A.P., Dolinski, K., Dwight, S.S., Eppig, J.T. et al. (2000) Gene ontology: tool for the unification of biology. The Gene Ontology Consortium. Nat Genet, 25, 25–29.

107. Esling, P., Lejzerowicz, F. and Pawlowski, J. (2015) Accurate multiplexing and filtering for high-throughput amplicon-sequencing. Nucleic Acids Res, 43, 2513–2524.

108. Desai, A.P., Razeghin, M., Meruvia-Pastor, O. and Pena-Castillo, L. (2017) GeNET: a web application to explore and share Gene Co-expression Network Analysis data. PeerJ, 5, e3678.

109. Vandesompele, J., De Preter, K., Pattyn, F., Poppe, B., Van Roy, N., De Paepe, A. and Speleman, F. (2002) Accurate normalization of real-time quantitative RT-PCR data by geometric averaging of multiple internal control genes. Genome Biology, 3.

