## Supplemental information for "Multi-influential genetic interactions alter behaviour and cognition through six main biological cascades in Down syndrome mouse models"

### Detailed Materials and Methods

All experiments were performed in accordance with the Directive of the European Parliament: 2010/63/EU, revising/replacing Directive 86/609/EEC and with French Law (Decret n° 2013-118 01 and its supporting annexes entered into legislation 01February 2013) relative to the protection of animals used in scientific experimentation. YH was the principal investigator of this study (accreditation 67-369) in our animal facility (Agreement C67-218-40). The mouse lines are available through the INFRAFRONTIER/European Mouse Mutant Archive.

To generate the Dp5Yah duplication, we used LoxP sites inserted at the *App* and *Runx1* loci as described previously (1-3) and using an *in vivo* TAMERE strategy (4, 5). Dp5Yah mice carry a duplication of the Mmu16 region located between 75752482 and 92698235bp (GRCm38/mm10) (Supplementary fig 1). Mouse lines were genotyped according to published protocols (6-9). The Dp5Yah mice were genotyped by PCR with common primer App_wt/rec3Up (5’GGCTCATCCTTGTTCTTTGTT3’) and two specific primers App_wt2Dw (5’AGCAGAGCACCTTCGTGTTT3’) and App_Rec3Dw (5’CGTCTTGCCTGGAGTTATTTT3’). For Dp3Yah mice, the PCR genotyping strategy used three pairs of primers, Stch_wt/recDw (5'GGCTTGGCTCCAAAACTACA3'), Stch_wtUp (5'TGGGCCATACTAGCATCAACT3') and Stch_recUp (5'GAGTCAGTGAGCGAGGAAGC3'). MBACtgDyrk1a transgenic line was maintained on a C57BL/6J background and genotyped by PCR with primers Dyrk-L (5′-TGGGCCAAGCAGTTAGGAGTTT-3′) and Bac11-R (9) (5′-CCATGATTACGCCAAGCTATTTAGG-3′).

#### Behavioural analysis

We generated experimental animal cohorts by selecting mice from litters containing a minimum of two male pups. After weaning, animals were sorted by litters into 39 x 20 x 16 cm cages (Green Line, Techniplast, Italy) where they had free access to purified water and food (D04 chow diet, Safe, Augy, *France*). The temperature was maintained at 23±1 °C, and the light cycle was controlled as 12 h light and 12 h dark (lights on at 7 am). On testing days, animals were transferred to the antechambers of the experimental room 30 min before the start of the experiment. All experiments were performed between 8:00 AM and 4:00 PM. A resting period of 2 days to 1 week was used between two consecutive tests.

##### Y-maze

Short-term memory was assessed by recording spontaneous alternation in the Y-maze test (10). The Y-maze test is based on the innate preference of animals to explore an arm that has not been previously explored, a behaviour that, if occurred with a frequency greater than 50%, is called spontaneous alternation behaviour (SAB). The maze was made of three enclosed plastic arms, each 40x9x16cm, set at an angle of 120° to each other, in the shape of a Y. The wall- of each arm have a different pattern to encourage SAB. Animals were placed at the end of one arm (this initial arm was alternated within the group of mice to prevent bias of arm placement), facing away from the centre, and allowed to freely explore the apparatus for 8 min under moderate lighting conditions (70 lux in the centre-most region). The time sequences of entries in the 3 arms were recorded, (considering the mouse enters an arm when all four paws were inside the arm). Alternation was determined from successive entries to the three arms on overlapping triplet sets in which three different arms are entered. The number of alternations was then divided by the number of alternation opportunities namely, total arm entries minus one. In addition, total entries were scored as an index of locomotor activity.

##### Open field (OF)

The open field test measures a combination of locomotor activity, exploratory drive and some aspects of anxiety and fear in mice. The output of the various interacting drives is locomotion, which is the direct measure obtained. The OF was performed in a white circular arena (55-cm diameter) placed in a dimly lit testing room (40 lux). The mice were introduced for 15 min in this arena. During those sessions, mice were monitored using a video tracking system (Ethovision, Wageningen, The Netherlands), and the total distance travelled in the different zones of the arena (centre, intermediate and peripheral) was recorded.

##### Novel Object Recognition (NOR)

The novel object recognition task is based on the innate tendency of rodents to explore novel objects over familiar ones (11). This test was done 24 hours after the OF session performed in the same arena. On Day 1, mice were free to explore 2 identical objects for 10 min. After this acquisition phase, mice returned to their home cage for 24 hours retention interval. In order to test their memory, on day 2, one familiar object (of those already experienced during the acquisition phase) and one novel object were placed in the arena and mice were free to explore the two objects for a 10 min period. Between trials and subjects, the different objects were cleaned with 70° ethanol in order to reduce olfactory cues. To avoid a preference for one of the two objects, the new one was different and counterbalanced between the different animal groups and genotypes. Similarly, to avoid a location preference, the emplacement of the novel object compared to the familiar one (left or right) was counterbalanced too. Object exploration was manually scored and defined as the orientation of the nose to the object allowing a distance < 1 cm. For the retention phase, the percent of time exploring familiar vs novel objects was calculated to assess memory performance.

Morris Water Maze (MWM)

Spatial learning can be analysed using the MWM task. This test was designed for the animals to learn to navigate a swimming tank (150 cm of diameter) filled with opaque water, following the most direct path to the hidden submerged platform when started from different, random locations around the perimeter of the tank. The use of distal cues provides the most effective strategy to accomplish this and escape the aversive effect of the cold water (22°C). This ability is controlled by hippocampal-dependent spatial cognition. The partial trisomic mice were trained in the standard version of the water maze as previously described (12, 13). This standard version contains 3 different phases, a learning phase (using 5 to 8 acquisition sessions, depending on the time taken by the mice to reach the plateau), a probe test (to evaluate memory performance) and a cue session (to validate the test). Two principal axes of the maze were defined, each line bisecting the maze perpendicular to one another to create an imaginary '+'. The end of each line demarcates four cardinal points: North (N), South (S), East (E) and West (W) and four quadrants (NE, NW, SE, and SW). Each acquisition session contained 4 trials in which mice were placed at one of the starting locations in random order (N, S, E, and W) and were allowed to swim until they located the platform situated in the target quadrant. Mice failing to find the platform within 60 s were gently guided and placed on it for 20 s (the same period of time as the successful animals). At the end of the learning phase, when the mice reached the platform, the probe test was done, with the platform removed and the time spent in the target and non-target quadrants as well as the number of platform annulus crossings during 60 s were recorded. After 24 hours, the cued session was performed to test the swimming speed and visual ability using the visible platform, clearly indicated by a visible cue (black flag). All the trials were recorded with a video tracking system (Ethovision, Wageningen, Netherlands).

Fear conditioning (Contextual and Cued)

To further challenge hippocampus mediated cognitive behaviours we used the fear conditioning test. Fear conditioning (FC), is an associative learning paradigm for measuring aversive learning and memory where a neutral conditioned stimulus (CS) such as light and tone are paired with an aversive unconditioned stimulus (US) such as mild shock in the paw. Concomitantly, animals associate the spatial context cues with the CS. After conditioning, the CS or the spatial context elicits a central state of fear in the absence of the US, translated in a reduced locomotor activity or total lack of movement (freezing) response. Thus, immobility time is used as a measure of learning/memory performances (14, 15). Experiments were conducted in four operant chambers (28 × 21 × 22 cm) with a metal bar floor linked to a shocker (Coulbourn Instruments, Allentown US). Chambers were dimly lit with a permanent house-light and equipped with a speaker for tone delivery and infra-red activity monitor. The experimental procedure encompassed 3 sessions over 2 days where the activity/inactivity behaviour was monitored continuously and the duration of inactivity per 2 s was collected. In day 1, for the conditioning session, the mouse was allowed to acclimate for 4 min, then a light/tone (10 kHz, 80-dB) CS was presented for 20s and terminated by a mild shock in the paw, US (1sec, 0.4 mA). After the paw shock, animals were left in the chamber for another 2 minutes. We defined total freezing time in first 2min and 4min and 2min immediately after paw shock as PRE1, PRE2 and POST, respectively. In Day 2, the fear to context was tested by bringing back the mouse into the same chamber and allowing it to explore for 6 minutes without presentation of the light/auditory CS. The movements of the animal were monitored to detect freezing behaviour consequence of recognizing the chamber as the spatial context (contextual learning). The total freezing time was calculated per 2min time block as CONT2, CONT4 and CONT6. Finally, the cue testing was performed 5 hours after the context testing. Animals were tested in modified conditioning chambers with walls and floor of different colour and texture. The mouse could habituate for 2 minutes to the chamber and then it was subjected to light and auditory cues for 2 minutes to evaluate conditioning fear. The total freezing time was calculated by 2min block as PRECUE1, CUE1, PRECUE2 and CUE2.

**Statistical analysis of behavioral results**

A summary of all the behavioural tests performed is included in Supplementary table 1. For each data set, we analysed if the data were normally distributed by Shapiro–Wilk test and Quantile-Quantile plots (Supplementary fig S1) and the homogeneity of variances by Brown-Forsythe test. Statistical significance of differences between groups was inferred by one-way ANOVA (Open field) and ANOVA for repeated measures (MWM, FC) or in case of datasets where the assumptions of normality or homogeneity of variances were not fulfilled, we performed the Kruskal-Wallis non-parametric test. For NOR and Y-maze, we performed a one sample t-test on the percentage of novel object sniffing time or on the percentage of spontaneous alternation versus 50% (hazard). The post hoc tests (Fisher LSD Method) were conducted only if F in ANOVA achieved 0.05 level. In addition, for the Y-maze and the NOR, we performed a one sample t test versus 50% (the randomness probability) for the % of alternation and the sniffing time for the NO, for the two genotypes.

**MRI capture, processing and analysis**

Images were acquired using MRI on fixed brain samples with an actively decoupled quadrature-mode mouse brain surface coil for signal reception and a 72-mm birdcage coil for transmission, both supplied by Bruker. Two imaging protocols were used. The first protocol consisted of a 3D T2-weighted rapid-acquisition with relaxation enhancement (RARE). The parameters for this sequence were: TR 325 ms, TE 32 ms, Rare factor = 6, interecho spacing 10.667 ms, 92 kHz bandwidth. The second imaging protocol consisted of a 3D T2*-weighted Fast Low Angle (FLASH) sequence with the following parameters: TR 50 ms, TE 25 ms, FA 50 degrees, 28 kHz bandwidth. The output image matrixes for both sequences were 195 x 140 x 90 over a field of view 19.5 x 14.0 x 9.0 mm3 yielding an isotropic resolution of 100 µm and were reconstructed using ParaVision 6.0.

Morphological MRI images were compared across groups according a region-based analysis. In order to benefit from the information from both imaging modalities, each modality was processed separately, and the resulting region-based volume estimations were averaged out for each animal before the statistical analysis.

Each MRI image was segmented into twenty anatomical structures according to a multi-atlas label propagation framework. To this end, the ten manually segmented in-vitro MR images from the MRM NeAt Mouse Brain Database (http://brainatlas.mbi.ufl.edu/) were considered (16). The image processing pipeline consisted of the following steps: (i) a skull-stripping step was first performed using the tissue brain segmentation method provided in SPMMouse (http://www.spmmouse.org) (17); (ii) each MR image was then corrected for bias field in homogeneity using N4ITK (18); (iii) the ten anatomically annotated images from the MRM NeAt Mouse Brain Database were registered in a deformable way on each mouse image using the ANTs registration toolbox (http://stnava.github.io/ANTs/) (19); (iv) anatomical labels were finally fused using the simultaneous truth and performance level estimation (STAPLE) (20).

By this way, the volumes of the twenty anatomical structures as well as the whole brain were computed for each image modality and each mouse. Finally, the volumes computed from the two images modalities were averaged out to obtain the final volume associated to each mouse.

#### Whole-genome expression arrays

Biotinylated single strand cDNA targets were prepared, starting from 150 ng of total RNA, using the Ambion WT Expression Kit (Cat # 4411974, Ambion, Life Technologies, Carlsbad, California, USA) and the Affymetrix GeneChip® WT Terminal Labeling Kit (Cat # 900671) according to Affymetrix recommendations. Following fragmentation and end-labelling, 1.9 µg of cDNAs were hybridized for 16 hours at 45°C on GeneChip® Mouse Gene 1.0 ST arrays (Affymetrix, Santa Clara, CA) interrogating 28.853 genes represented by approximately 27 probes spread across the full length of the gene. The chips were washed and stained in the GeneChip® Fluidics Station 450 (Affymetrix) and scanned with the GeneChip® Scanner 3000 7G (Affymetrix) at a resolution of 0.7 µm. Raw data (.CEL Intensity files) were extracted from the scanned images using the Affymetrix GeneChip® Command Console (AGCC) version 4.0 and are stored for public use in the NCBI GEO database (GSE149470).

#### Transcriptomics analysis

The gene expression profile of the following DS trisomic mouse models, Dp1Yey, Dp3Yah, Ts65Dn, Dp5Yah, Dp1Rhr, Dp5Yah/Dp1Rhr (noted also as Dp5/Dp1 or Dp1Rhr_Dp5Yah), and Tg(*Dyrk1a*) was analysed using a bioinformatics pipeline, developed over the R environment (21) using several publicly available packages. The full list of R packges used and their references is in Supplementary table 6. The R and Linux bash scripts were implemented in both an Ubuntu pc (62,8 GiB memory, and with a processor Intel® Xeon(R) CPU E3-1270 v5 @ 3.60GHz × 8), and a MacBook Pro (Retina, 2,6 GHz Intel Core i5, 8 GB 1600 MHz DDR3). The pipeline consisted in seven major steps: Quality control (QC), pre-processing and normalization of the raw data, analysis of the structure and homogenicity of the data, differential expression analysis (DEA), differential functional analysis (DFA), Network connectivity analyses of pathways and genes and finally network topology and centrality analyses.

**Data quality control:** The quality control (QC) workflow was implemented in two steps. The first step consisted in analysing the quality of the raw data, and the second in the analysis of the quality after the pre-processing and normalization steps. To assess the quality of the raw data we analysed the following parameters. First, the signal distribution using both boxplots of log-intensity and density histograms. The boxplots of log-intensity allow inter-array comparison and to check the effect of the normalization as the probe-set log-intensities of all the arrays are more similar to each other than the raw log-intensities. Instead, the density histograms show the intensity distribution of each array and plotted in a unique plot, allows a better comparison between arrays, the identification of odd distributions and to check the effect of the normalization on the intensity distribution. Second, the Intensity-dependent bias using MA-plots. MA-plots are used to compare each Affymetrix array to a pseudo-array computed as the median intensity of each probe over all arrays. These plots evaluate the dependency of the measured variability in expression levels with the expression level. As a result, the difference between the intensity of a probe on one array and the median intensity of that probe over all arrays (M) should be similar across the arrays. Besides, the average of the intensity of a probe on one array and the median intensity of that probe over all arrays (A), should be centred around the blue line (M=0). Instead when (A) increases, the loess curve (red line) moves away from M=0. Therefore, to remove as much as possible this dependency, the data was normalized. Third, the spatial bias was considered using 2D images, pseudo-images and residuals plots (not shown). The pseudo-image plots assess the spatial distribution of the data on the arrays to identify the existence of abnormal patterns or spatial quality problems may affect our downstream analysis. The external control .CEL files are available in ArrayExpress and GEO database, under the identifiers E-GEOD-62538 and GSE62538 respectively. In comparison with the control arrays, we did not find any worrisome quality problem in the spatial bias assessment of the arrays even though a few small spots and edge effects in some of the arrays were present.

**Data pre-processing and annotation:** The raw intensity measurements of the arrays where normalized using limma RMA normalization algorithm (22) producing the summaries at the probe/transcript level**.** The probes were gene and transcript annotated using the specific R package pd.mogene.1.0.st.v1 (23) and further annotated using ENSEMBL BioMart databases from biomaRt R package (24, 25) updated on 12/02/2018 against the mm10 genome assembly. The non-annotated probes and control probes were discharged from the analysis. Of 35556 probes, 27359 were annotated.

**Data structure and homogeneity:** We performed a study of i) the gene expression correlation across models and ii) the structure of the data and the separation between experimental groups using hierarchical clustering, PCA, tSNE (26) and OPLS. First, the pairwise correlation coefficients were computed and shown on a matrix giving an idea of the arrays homogeneity. Low coefficients indicate important differences between array intensities. (Supplementary figs 9 A and B). We computed the correlation considering all the expressed genes identified (EGs) (data not shown), the differential expressed genes in all models (4328 DEGs) as shown in Supplementary fig 9A or uniquely the differential expressed trisomic genes (75 TEGs) whose expression was captured in all the arrays (Supplementary fig 9 B). The plots were created in R using the pairs.panel function from Psych R CRAN package (27). Each row represents each model pairwise correlation with the rest of the models. The density and histograms distribution of the gene expression values for each model is found in the medial diagonal (Black for DEGs, pink for TEGs). The lower part of the matrix provides a visual representation of each gene expression level over each pairwise comparison of models, whereas the upper part of the matrix shows the numeric correlation calculated. The LOESS smoothed fit is represented by a red line.

Second, to study the structure of the data we performed several analyses such as principal component analysis (PCA). PCA, is a well-known dimension reduction statistical method based on the orthogonal transformation of the observations of could be correlated variables into a list of values of linearly uncorrelated variables, called the principal components (PCs). Afterwards PCA-kmeans, Orthogonal Projections to Latent Structures Plots (OPLS) and t-Distributed Stochastic Neighbor Embedding (t-SNE) plots (26) (See Fig 2 B) were computed to further assess the structure of the data and homogeneity. These methods gave us similar profiles on the overall structure of the data although in some cases increasing the resolution.

**Differential expression analysis (DEA):** We carried on the differential expression analysis in R environment using FCROS (28, 29). FCROS is a fold change (FC) rank based method that works well with noisy datasets and gives strong reproducible results. FCROS computes for each pair of test/control samples (K pairs), a statistic associated with the k ranks of the FC values for each gene, and the obtained probability (f-value) is used to identify the differential expressed genes (DEGs) within an error level fixed by the analyst. Our fixed error level corresponds to a 5% False Discovery Rate (FDR), and is attained by setting the $\alpha$ parameter within this range $0.025<\alpha>0.975$. The summary of the results obtained is shown in Supplementary table 2

Afterwards, to highlight both the number of genes identified as DEGs and the ones with higher log2 fold-change (log2 FC) values for each mouse model we used Volcano plots of the log2 FC versus −log10 of the FDR corrected p-value (Supplementary fig 8). Each gene was classified into one of the following four groups, not significant, significant, FC and significant, or FC as follows: a) Not significant: genes not identified by FCROS as DEGs based on the alpha cut off; b) Significant: DEGs identified by FCROS based on the alpha cut off, whose FDR corrected p-values are < 0.05; c) FC: All the expressed genes whose abs (FC) value > 1.4. d) FC: DEGs identified by FCROS based on the alpha cut off, whose FDR corrected p-values are < 0.05 and whose abs (FC) value > 1.4.

Identification of common and unique DEGs across models

We studied the number and name of the common and unique DEGs per model identified by the FCROS method, performing the intersection between the lists of DEGs from the different models and using VennDiagram (30) for the visualization of up to 5 models (See Fig 2 C). Vennerable was used for drawing the 7 models together in the R environment (31).

FC expression profile of the genes included in the duplicated region:

To assess the gene expression variation on the genes included on each chromosome 16 (chr16) trisomic region, we plotted the FC values along the genes ordered by chromosomal location (Supplementary fig 5). Additionally, a similar plot was built of the chromosome 17 (chr17), region as the Ts65Dn model have a break point and carries a duplication of part of this chromosome not homologous to Hsa21. (See Supplementary fig 6)

**Differential functional analysis (DFA):** To identify the altered functions due to the changes in gene expression and possibly get new insights into the mechanistic changes along the pathways between the two conditions trisomic vs. wild type on each mouse model, we used both GSEA Desktop Application (32) and generally applicable gene set enrichment for pathway analysis (GAGE) R package (33) to carry on the functional expression analysis. Both methods check for the existence of differences in the expression over gene sets (functional related genes) instead of focusing on identifying changes in individual genes based on prior pathway knowledge and your resulting list of DEGs identified.

The use of GAGE has several advantages over other typical GSEA methods, as the authors described on the paper (33). First GAGE accounts for both gene set specific variance and background variance. Secondly, GAGE is able to adjust for different experimental designs and sample sizes by dividing group-on-group comparisons into one-on-one comparisons between samples from different groups to finally compute a global p-value using the Stouffer's summarization on the p-values from these comparisons for each gene set, this is an advantage over the classical GSEA. Last, GAGE separates experimentally probed perturbed gene sets from canonical pathways increasing the robustness and test power in identifying meaningful biological differences and also provides the framework to conduct a unified and rigorous FDR procedure for gene sets of different sizes.

We developed and run the gage workflow in R, following the joint workflow with limma as described over the gage manual.

1) The Gene-set data, was compiled from mouse KEGG Onthology and the different GO (Gene Ontology) terms, in real time on the 12/02/2018.

2) We used our already RMA normalized counts and the raw counts normalized following the recommended protocol by GAGE on the datasets with annotated gene Names (27359 probes) to assess the effect of the normalization protocol on the results.

3) The probes were annotated with the ENTREZ Gene ID

4) The gage function was run, using compare=”as.group”, use.stouffer=TRUE and same.dir = TRUE for KEGG, and run both with same.dir = TRUE and FALSE for all the GO gene sets ( The GO terms are subdivided in three categories: cellular compartment (noted CC), biological processes (noted as BP) and molecular functions (noted as MF)).

5)We selected as Differentially altered pathways those KEGG and GO terms whose q-value <0.1 as recommended by the authors of GAGE (33).

6) Two functions were used to generate a report file. First, EssGene function, select genes with above-background expression changes in gene sets or pathways. Secondly, the geneData function generate an output report file per each combination of mouse model, regulation sense of the identified pathway and the name of the pathway. These final report files include the following information: MiceLineName (name of the mouse model), PathwayReg (Regulation sense of the pathway as upregulated or downregulated), PathwayName (name of the pathway identified as altered), GeneENTREZ_ID (ENTREZ id code), and the expression levels for both wt and trisomic samples. Then we combined all the reports using an *in-house* bash script, imported it to R and re annotated the gene name using the ENTREZ IDs names.

7) The pathways and GO terms shared between at least two models were grouped into 8 higher categories or meta-pathways we defined to be able to shrink the broad dysregulation into few mayor functionality groups. Those meta-pathways were thus, higher functional groups formed

by the inclusion of pathways involved in similar functionalities defined as follows: i) synaptic related, ii) transcription & epigenomics regulation, iii) enzymes activity, iv) ribosome related, v) mitochondria related, vi) cell structure & organelles, vii) phospho-kinase related and viii) compounds binding, ix) Hormone regulation, x) Post-translation modification. The classification of the individual pathways into those meta-pathways was performed using natural language processing (NLP) by defining associative common patterns found in the KEGG and GO terms. The pathways not fitting in any group where left ungrouped and labelled with the unique pathway name. The visual representation of the functional dysregulation found in each model over the meta-groups and individual pathways was visualised using a heatmap (See Fig 2 D). Finally, we split the synaptic related group into four sub-meta-pathways: GABA system, Myelin sheath: SNARE complex pathways, Behaviour involving pathways, and the rest. The main reason for this distinction was due to the observation of a different regulatory profile of Dp5Yah and Dp1Yey over some of those sub-groups (See Fig 2 B). The heatmap representation in both cases, reflects the overall up or down regulated profile for each meta-pathway or ungrouped pathway (rows) per mouse model (columns), showing first the meta-pathways and after those, the ungrouped pathways names. The colour scale deriving from pink to purple represents up regulated and down regulated pathways respectively, the crossing over in white-grey represent that none pathway was found to be included on that meta-pathway. The intensity of the colour corresponds to the number of pathways inside of those meta-pathways. The heatmap colour key breaks were defined representing this number of pathways 200, 150, 100, 50, 25 and 5 up regulated (purple) and down regulated (pink).

GAGE was able to achieve a better performance than GSEA, and we were able to identify up and down regulated gene sets and pathways, meaningful and correlated with the samples source after q-val correction. Thus, we carried on further analyses using uniquely GAGE functional results.

**Network connectivity analyses of pathways and genes:** We performed two different network analyses, one at the pathways level and another at the gene level to assess on the first case inter-pathway connectivity and gene redundancy, and on the second, the gene connectivity and crosstalk.

Computing grouped weighted networks:

We generated functional weighted networks for each meta-pathway, considering only the relationships between the pathways included on one meta-pathway at a time. All the analysis was done in the R environment using in house scripts. We defined the weights as the number of altered genes identified by GAGE shared between the pathways and each mouse model or combination of mouse models (mouse model multi groups) to analyse the inter-pathway relationships and gene redundancy. The strength of the pathways connectivity within each meta-group was represented by the thickness of each edge connecting the pathways (nodes) in a weighted network built specifically for each meta-group. The different coloured edges joining each pathway and edges thickness represents on each specific mouse model multi groups (coloured edges), the unique number of genes connecting those pathways on each specific mouse model multi groups (edges thickness) (See Supplementary figs 10, 11, 12, 14)

Building up the Protein-protein interaction and regulatory gene connectivity networks (MinPPINet and RegPPINets)

To further characterize how the different models show conserved or distinct altered genes on each shared meta-pathway we built gene connectivity networks using the protein-protein interaction information store in STRING database (34) and the regulatory information contained in REACTOME (35). Those networks could reveal the central genes and biological cascades affected, linked to connected functionalities. Furthermore, including regulatory information may give some insights into the altered gene and transcription factor dynamics to identify druggable hubs to further study. We used NetworkAnalyst (36) to generate a preliminary minimum network based on the PPI selected from STRING DB (34) by fixing different confidence thresholds to allow their inclusion on the network building process. We selected different confidence levels 700 (medium-high confidence), 500 (medium) and 400 (medium-low) to select the PPI. Both forcing and not forcing all the PPI considered to have being experimental validated. In the end the minimum network was built (MinPPINet) removing all the nodes that were not on the shortest paths connecting the seeds and it was exported to Cytoscape 3.5.1 (37). The regulatory network was built using ReactomeFIViz app in Cytoscape. The interactions reported by the app are classified following the KEGG Markup Language (KGML) hierarchical structure into four categories: ECrel, PCrel, GErel and PPrel interactions (edges). Each interaction also contains a name attribute specifying the type of interaction. Following PathPPI classification (38), we considered only GErel edges, indicating the source node to be acting like a transcription factor and the target node as a repressed or expressed gene and PPrel edges (protein protein activation or/and inhibition relationships), plus the FI annotated interactions labelled as complexes and predicted to highlight in our networks using different colours for the edges i) The GErel edges indicating expression were colored in blue and repression in yellow. ii) PPrel edges indicating activation were coloured in green, inhibition in red. Iii) Interactions between proteins known to be part of complexes in violet. Iv) Predicted interactions were represented in grey including the PPI interactions identified by STRING DB. Then both the PPI and regulatory networks were merged to create the final Protein-protein interaction and regulatory gene connectivity networks (RegPPINets) (See Fig 4 and Supplementary fig 14 A and B). Furthermore, the nodes where further annotated using an in-house R script to generate the proper annotations for each node attribute as specified below:

For the nodes involved in the synaptic meta-pathway:

If the node is a DEG it appears with a pyramid shape, and the colour of the shape represents the mouse models where it was found as DEG;

If the node is a DEG and part of the HSA21 synteny region in mouse it appears in a bigger height and width than the rest for the nodes.

If the node is not DEG but involved in the dysregulation of the pathways in some specific mouse models, the border colour represents the mouse models where this gene was found involved in the dysregulation by GAGE even if not identified as DEG by FCROS. Additionally, we wanted to further characterise the relationship of the genes involved in the pathways dysregulation with other DEGs in the HSA21 synteny region in mouse (that were not identified by GAGE as being contributing to those pathways). Those genes appear with a magenta octagonal shape with increased height and width. Finally, the connecting nodes incorporated to build a fully connected network but unknown by Go and KEGG gene: pathways associations to be involved in brain synaptic dysfunction are represented with a ellipse shape and soft pink colour. These nodes are necessary to maintain the integrity of the network and are novel genes that can be linked to brain synaptic dysfunction.

**Network topology and centrality analyses:** First, we analyzed different network topography parameters (shortest path length distribution, average clustering coefficient distribution, in-degree, out-degree, … ) and betweenness centrality of the synaptic MinPPINet, (Fig 4 A) to identify hubs and genes central for the flow of information along the network. Using a network decomposition approach, we were able to identify 5 mayor subnetworks that strongly centralized in five different proteins DYRK1A, GSK3B, RHOA, NPY and SNARE complex (See Fig 4 B, C). After subtracting from the network the first and second interactors of those proteins, no other sub-cluster of DEGs was left. Then, for each subnetwork a RegPPINet was built and annotated to provide further insights in the functional alteration and gene-gene gene-protein regulatory crosstalk. Furthermore, we identified a strong connectivity and lack of independency of three of those five main cascades represented by RHOA, GSK3B and DYRK1A (Supplementary fig 14 A) at the second interactors levels. We addressed how closely connected are those subnetworks by identifying i) the number of shared altered genes or “seeds” linked to synaptic function ii) the number of shared nodes (Supplementary fig 13 A) and the overall relevance of the seeds to the network connectivity by addressing how many seeds were shared compared to network nodes ( considering both seeds and connecting proteins) (Supplementary fig 13 B) iiI) the number of seeds or total nodes of the Synaptic RegPPINet included on each subnetwork (S6 table). Finally, we further characterized NPAS4 effect over the DS brain synaptic dysfunction network both experimentally and computationally, by analyzing the sub-network formed by extracting NPAS4 1$^{st}$ level interactors (Fig 3 C, bottom right) and the 3$^{rd}$ level interactors (Supplementary fig 14 B).

#### QRT-PCR

cDNA synthesis was performed using the SuperScript® VILO™ cDNA Synthesis Kit (Invitrogen, Carlsbad, CA). PCR were performed with TaqMan® Universal Master Mix II and pre-optimized TaqMan® Gene Expression assays (Applied Biosystems, Waltham, Massachusetts, USA), consisting of a pair of unlabelled PCR primers and a TaqMan® probe with an Applied Biosystems™ FAM™ dye label on the 5’ end and minor groove binder (MGB) and nonfluorescent quencher (NFQ) on the 3’ end (listed in Supplementary Table 4). mRNA expression profiles were analysed by real-time quantitative PCR using TaqMan TM universal master mix II with UNG in a realplex II master cycler Eppendorf (Hambourg, Germany). The complete reactions were subjected to the following program of thermal cycling: 1 cycle of 2 minutes at 50°C; 1 cycle of 10 minutes at 95°C; 40 cycles of 15 seconds at 95°C and 1 minutes at 60°C. Efficiencies of the TaqMan assays were checked using cDNA dilution series from extracts of hippocampal sample. Normalization was performed by carrying out in parallel the amplification of 4 housekeeping genes (*Gnas, Pgk1, Actb* and *Atp5b*) and using the GeNorm procedure in order to correct the variations of the amount of source RNA in the starting material (39). All the samples were tested in triplicate (Supplementary fig 15).

#### Western Blot

Fresh hippocampal tissues were isolated by rapid decapitation/dissection of 6-7 month-old mice and snap frozen. Samples were then lysed in ice-cold sonication buffer supplemented with

Complete™ Protease Inhibitor Cocktail (Roche). Individual samples were disaggregated, centrifuged at 4°C for 30 minutes at 14000 rpm, diluted in 4X Laemmli sample buffer containing β-mercaptoethanol (Bio-Rad, France) and incubated at 95 °C for 5 min. Protein concentration was determined by PierceTM BCA Protein Assay Kit (23225, Thermo Fisher Scientific, Strasbourg). The samples were further diluted in buffer to a concentration of 5ug/uL, 30 µg of protein was loaded per well onto a 15% polyacrylamide gel. The proteins were transferred to nitrocellulose membranes by Trans-Blot® Turbo™ Transfer System (BioRad, France) through MIXED MW Bio-Rad Pre-programmed Protocol. Then, the blots were incubated with the following blocking solution: 5% BSA, 1 X Tris-buffered saline, 0.1% Tween 20 (TBS-T) during 5 minutes and incubated with 1:1000 primary antibody diluted in blocking solution during 10 minutes. Blots were washed in TBS-T and incubated with 1:5.000 HRP conjugated Goat anti-Rabbit IgG secondary antibody (A16096, Invitrogen, France) diluted in blocking solution during 10 minutes. All incubations were made using the SNAP i.d.® 2.0 Protein Detection System (C73105, Merck). This apparatus has a vacuum-driven technology and a built-in flow distributor that actively drives reagents through the membrane.Proteins were visualized with Amersham™ Imager 600. Signals were quantified using ImageJ and the statistical analysis using Sigma Plot. We used the following primary antibodies: anti-RHOA (2117, Cell Signaling, USA, 1:1.000), anti-pMLC (Thr18/Ser19 #3674, Cell signalling, Boston, MA, USA, 1:1.000) and mouse monoclonal Anti-β-Actin−Peroxidase antibody (A3854 Sigma, 1:150.000). The relative amount of RHOA and p-MLC proteins was calculated as the ratio of the signal detected for each proteins of interest compared to the β-actin signal detected and normalized by the mean signal of the wt samples (Supplementary fig 16).

### Supplementary Figures

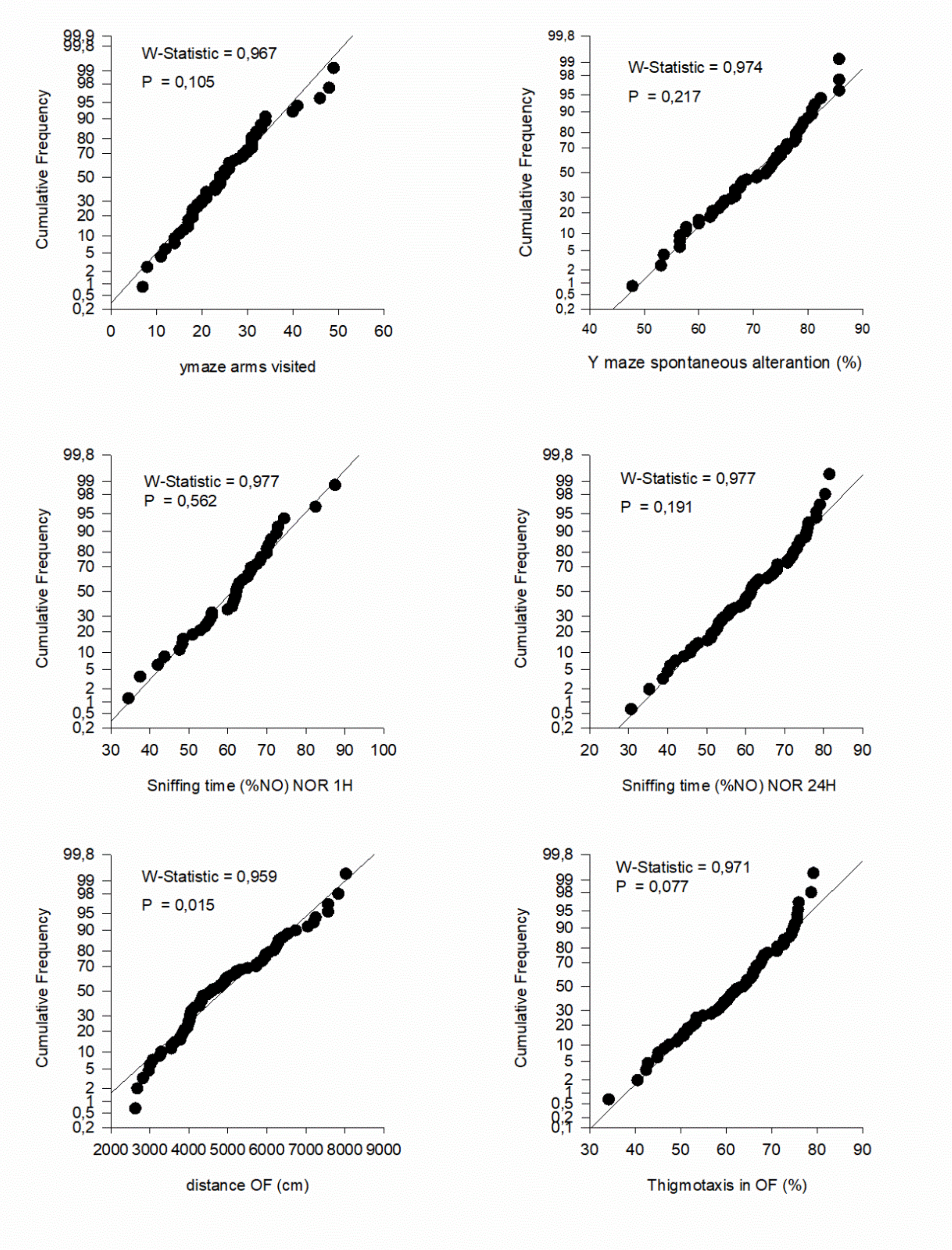


**S1 Fig. Quantile-Quantile plots to assess Normal distribution.** The quantile-quantile plots (Q-Q plots) and Shapiro-Wilk test for the following behavioural datasets: Y-maze, NOR and OF were analysed to study how well our data could be modelled following a normal distribution. Only the distance travelled in OF did not reach the statistical significance level in the Shapiro-Wilk test.

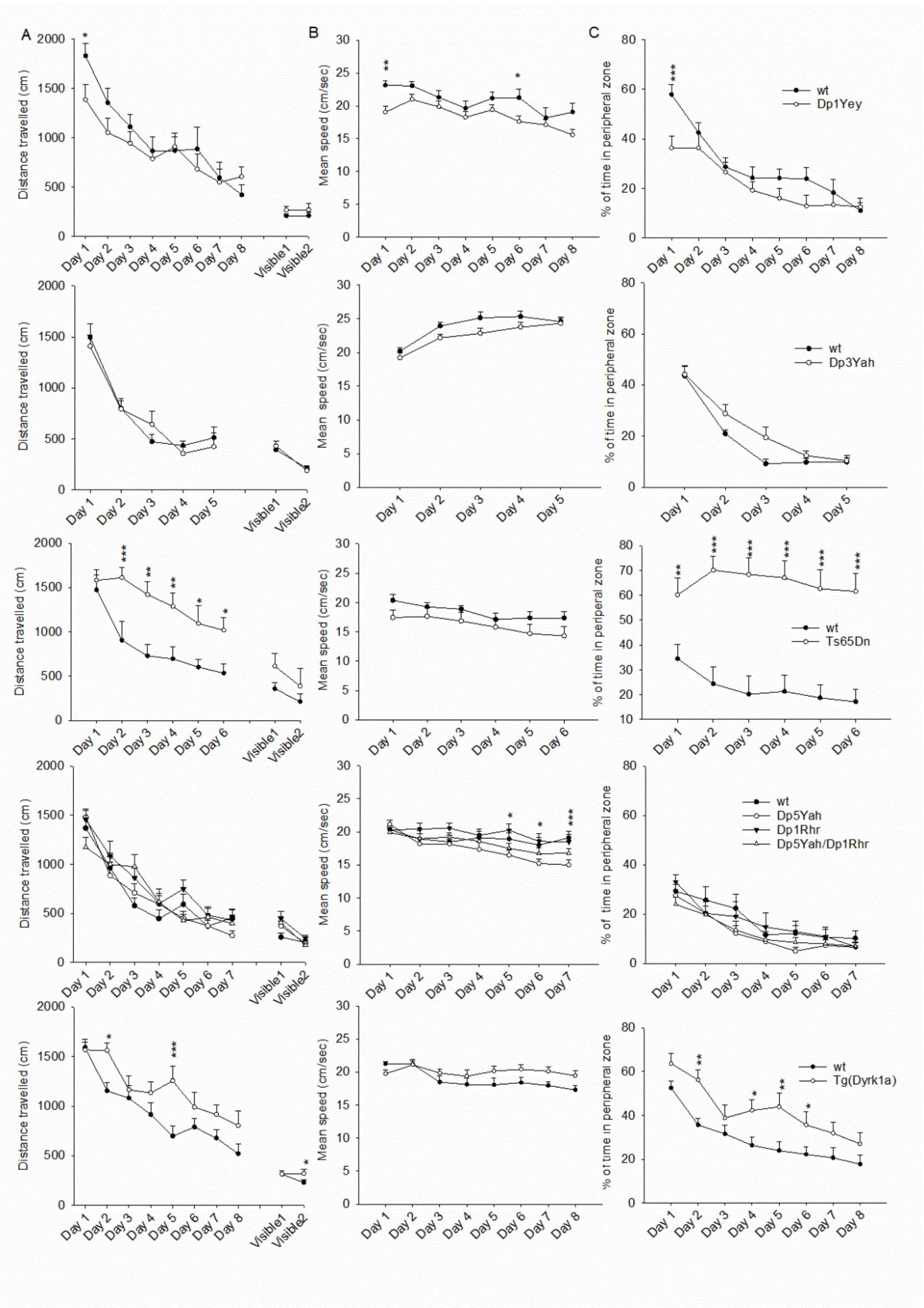


**S2 Figure. DS mouse models and Morris water maze.** The MWM results are presented as distance travelled to reach the platform (A), speed (B) and % of distance travelled in the peripheral zone (C, Mean ± SEM). Ts65Dn mice were drastically impaired with an increase in distance travelled to reach the platform and in the distance travelled along the peripheral zone. Less drastically Tg(*Dyrk1a*) mice showed an increased distance travelled to escape the pool and in the peripheral zone. The velocity of Dp1Yey and Dp5Yah, was slightly lower than the wild-type mice (* $p<0.05$, ** $p<0.01$, *** $p<0.001$; Dp1Yey n=9 wt and 10 Dp; Dp3Yah n=15 wt and 13 Dp; Ts65Dn n=10 wt and 11 Ts; Dp5Yah/Dp1Rhr n=13 wt, 13Dp5, 13 Dp1, 13 Dp5/Dp1; Tg(*Dyrk1a*) n=16wt and 15Tg).

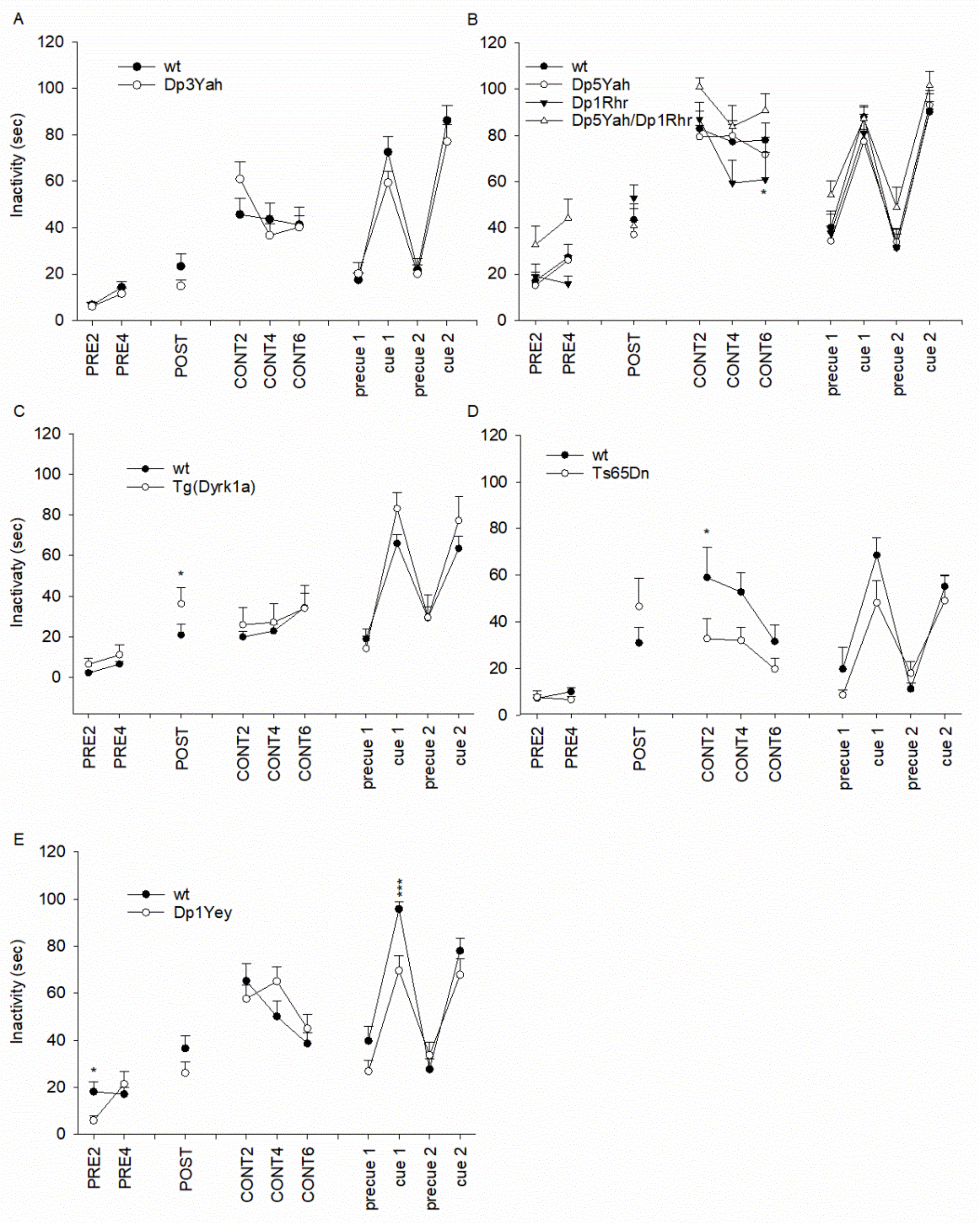


**S3 Fig. DS mouse models and context and cue fear conditioning.** The fear conditioning results are presented as inactivity periods in seconds (Mean±SEM): (A) Dp3Yah (n=15 wt and 15 Dp3Yah); (B) Dp5Yah/Dp1Rhr (n=13 wt, 13 Dp5Yah, 13 Dp1Rhr, and n= 13Dp5Yah/Dp1Rhr); (C) Tg(*Dyrk1a*) (n=12wt and 8 Tg(*Dyrk1a*)) ; (D) Ts65Dn (n=10 wt and 10 Ts65Dn); (E) Dp1Yey (n=21 wt and 20 Tg). Only Ts56Dn (D) mice present an impairment in context fear conditioning (* $p<0.05$, ** $p<0.01$, *** $p<0.001$.

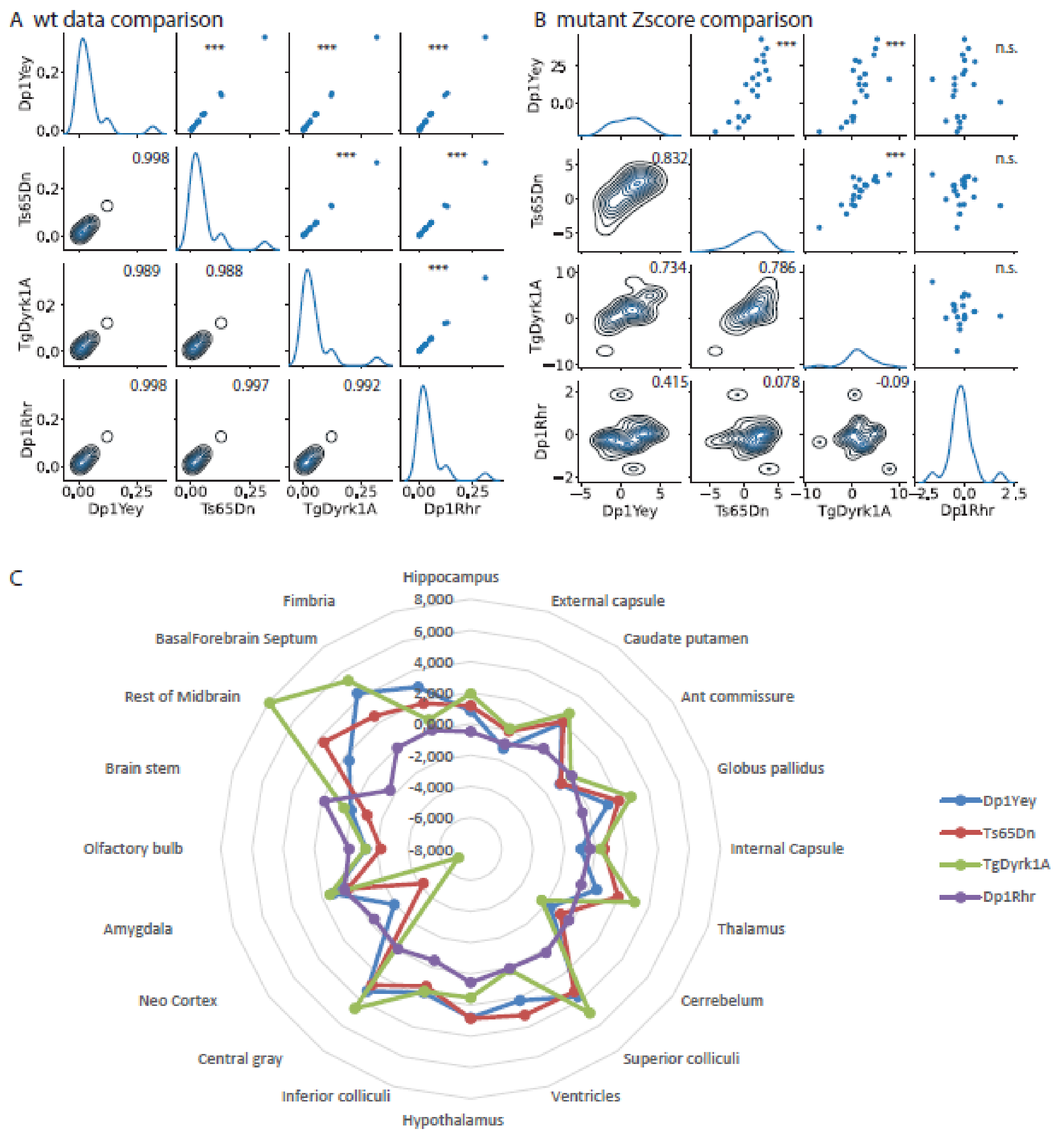


**S4 Fig. Supplementary Figure 4. IRM brain volume analysis.**

(A and B): upper and lower map of the grid represent respectively scatterplot and contour plot of two dimensional Kernel Density Estimate (KDE). The spearman corelation test value and significance level are indicate. Diagonal plot represent only one-dimensional KDE. (A) Analysis of the correlation based on the data of the wt in the different mouse line. Overall, the correlation was strong between all the wt without major consequences of the genetic background that was different in the Ts65Dn compared to the other lines. (B) Z-score analysis of the different mouse lines. The Z-score was calculated as the mean of wt mice minus the mean of transgenic divided by each type of wt and transgenic (Tg)/trisomic (Ts)/duplicated (Dp) line, per brain structures, per mouse lines. A strong correlation was observed between changes observed in the Dp1Yey, Ts65Dn and Tg(*Dyrk1a)* while no correlation was found with the Dp1Rhr line. (C): radar plot representation of the Z-score representation. The structures affected in Tg(Dyrk1a) were also affected in Dp1Yey and Ts65Dn, whereas no difference was observed for the Dp1Rhr mice. (Dp1Yey n=6wt and 6 Dp ; Ts65Dn n=5 wt and 6 Ts ; Tg(Dyrk1a) n=8 wt and 5 Tg ; Dp1Rhr n=7 wt and 7 Dp; * $p<0.05$, ** $p<0.01$, *** $p<0.001$).

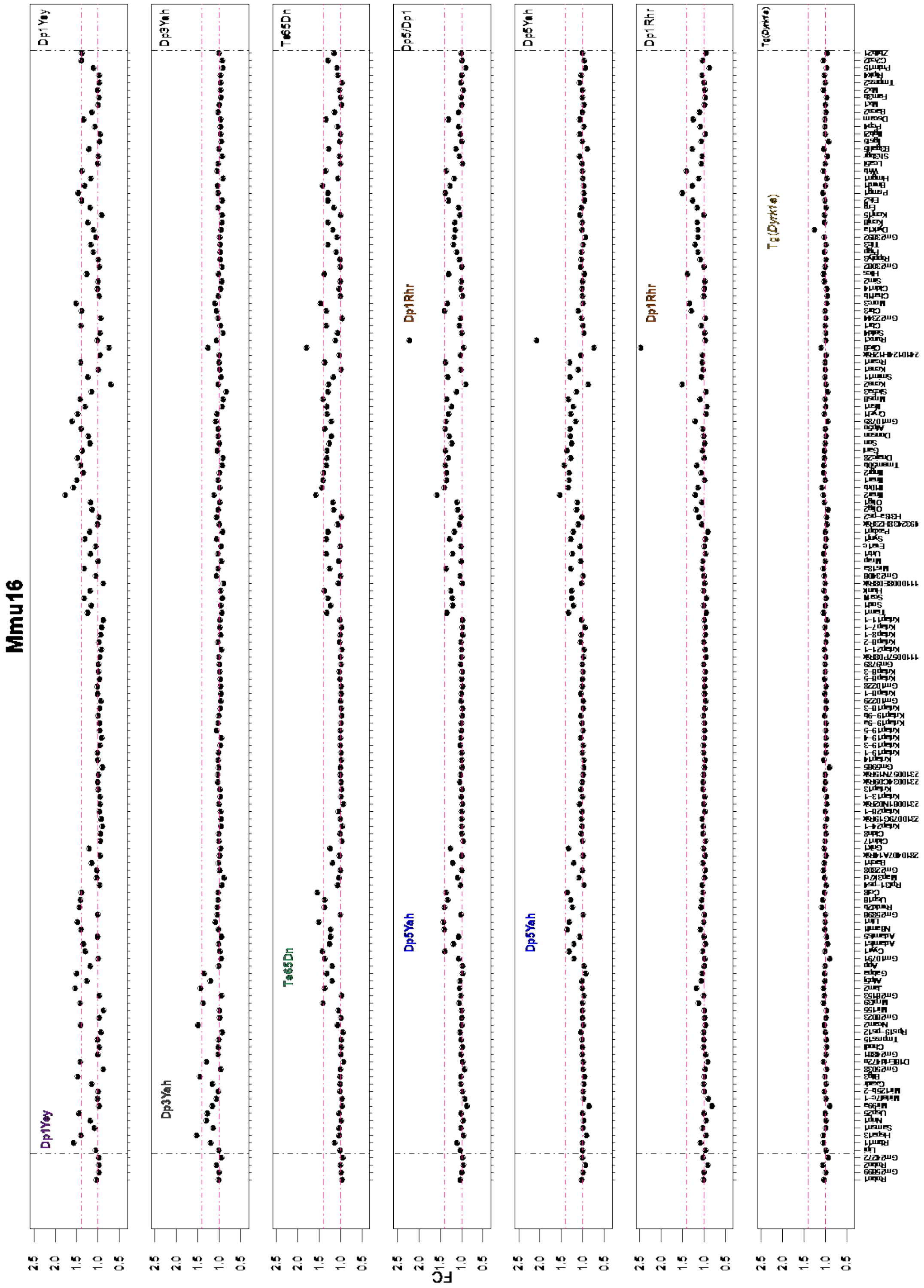


**S5 Fig. Fold change expression levels of the genes analysed by the microarrays homologous to the Hsa21 mouse chromosome 16 (mm16).** The genes are displayed following the order of their genomic start site coordinates. The duplicated areas for each mouse model appear shaded in the following colours, purple, grey, aquamarine, red, blue and golden for Dp1Yey, Dp3Yah, Ts65Dn, Dp5/Dp1, Dp5Yah, Dp1Rhr, and Tg(*Dyrk1a*) respectively.

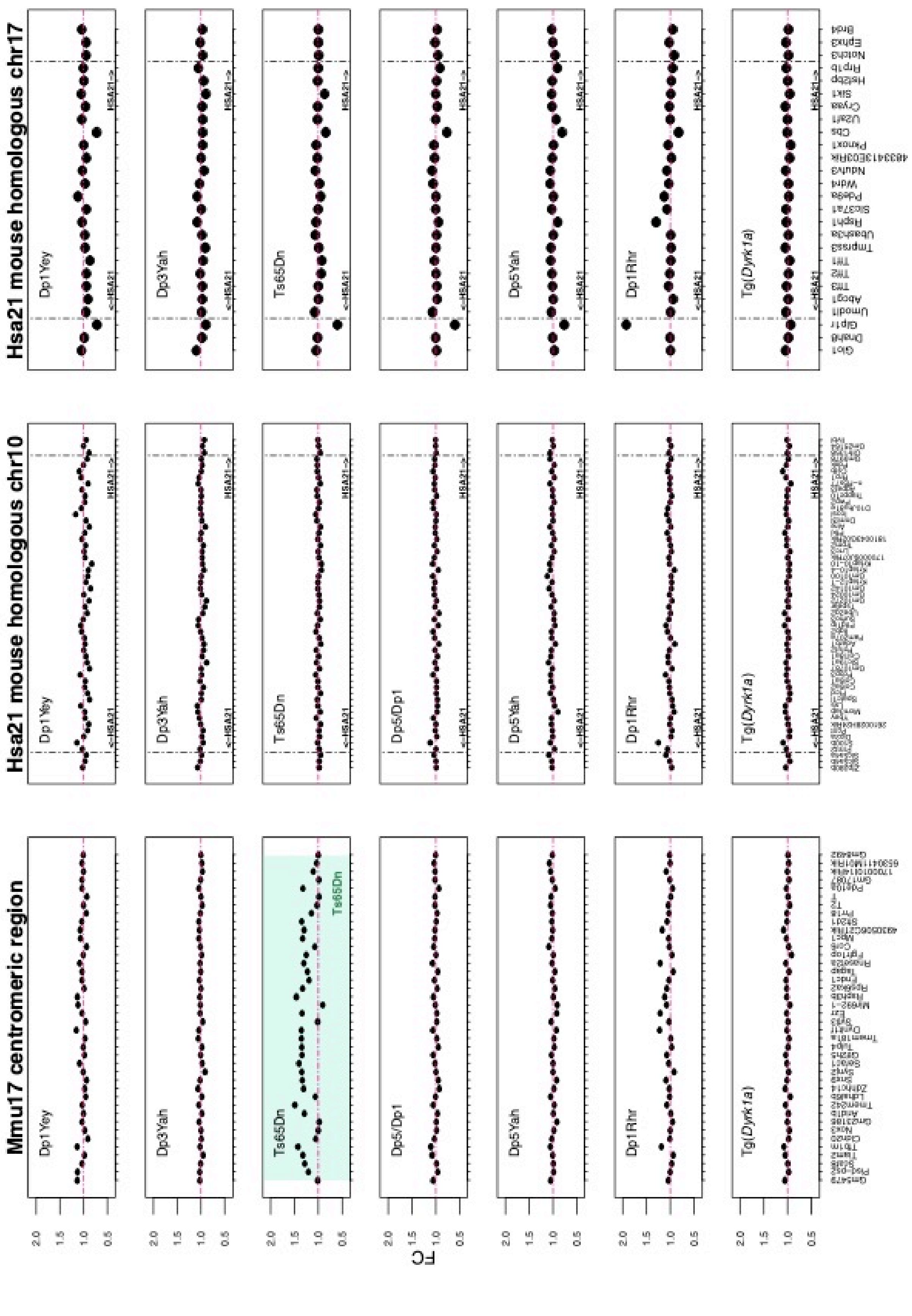

Mmu17 centromeric region
Hsa21 mouse homologous chr10
Hsa21 mouse homologous chr17
Dp1Yey
Dp3Yah
Ts65Dn
Dp5/Dp1
Dp5Yah
Dp1Rhr
Tg(Dyrk1a)
FC
<-HSA21
HSA21->
2.0
1.5
1.0
0.5

**S6 Fig. Fold change expression levels of genes analyzed by the microarrays.** (A) Fold change expression levels of the genes analysed by the microarrays part of the centromeric region of the mouse chromosome 17. The centromeric region of the mouse chromosome 17, is defined by the UCSC table browser as the region between the coordinates chr17:110000-3000000.The genes are displayed following the order of their start site genomic coordinates. The region duplicated in the Ts65Dn model non homologous to the hsa21, appears shaded in aquamarine. (B) Fold change expression levels of the genes homologous to the Hsa21 in mouse chromosome 10 (mm10) and chromosome 17 (mm17) analysed by the microarrays. The genes are displayed following the order of their start site genomic coordinates.

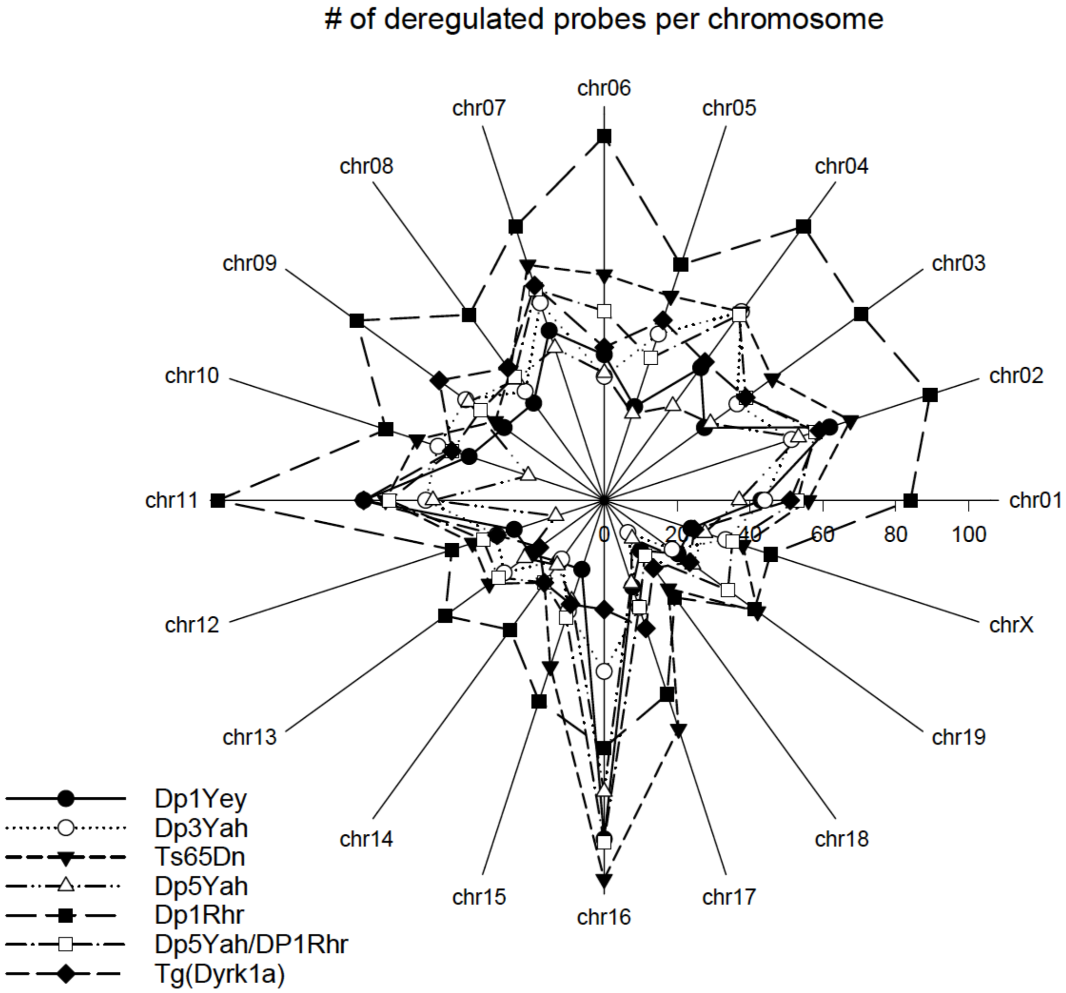


**S7 Fig. Number of deregulated probes per chromosomes per models**. Trisomic mice have similar patterns with a pic of deregulated genes located on mouse chromosome 16. Tg Dyrk1a at contrary, present a global pattern of deregulated probes.

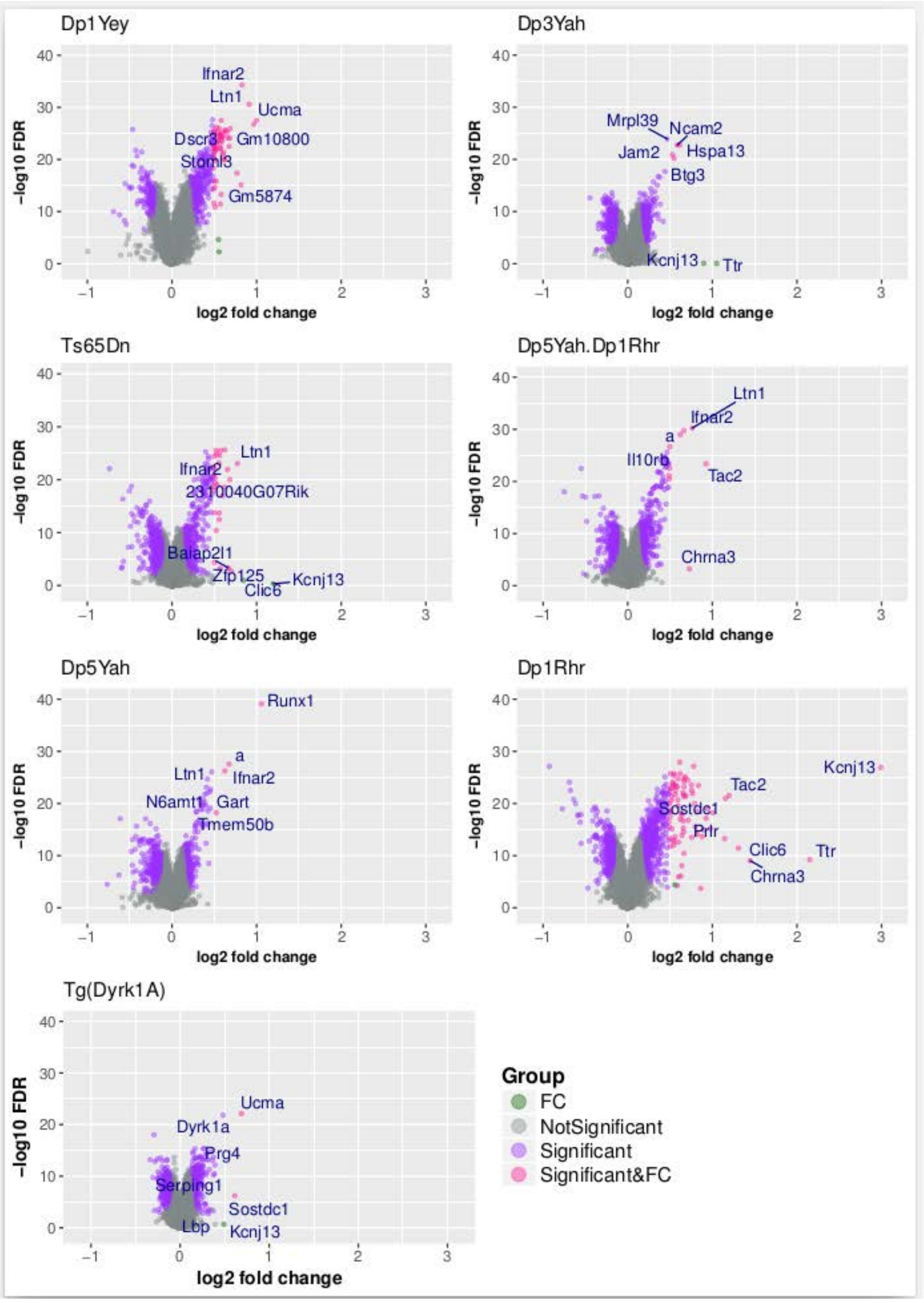


**S8 Fig. Overview of the top 7 dysregulated genes (higher significance and fold change difference) for each mouse model over the global transformed expression profiles illustrated over volcano plots.** The x axis and y axis represent the log2 fold change of the genes expression and the log transformed FDR values respectively. The genes whose fcros FDR value <0.05 are considered significant. These genes are represented in purple or pink and labelled as "Significant". Instead the genes whose abs (FC)> 1.4 are represented in green or pink and labelled "FC". Finally, the genes that don't comply these conditions are considered not significant.

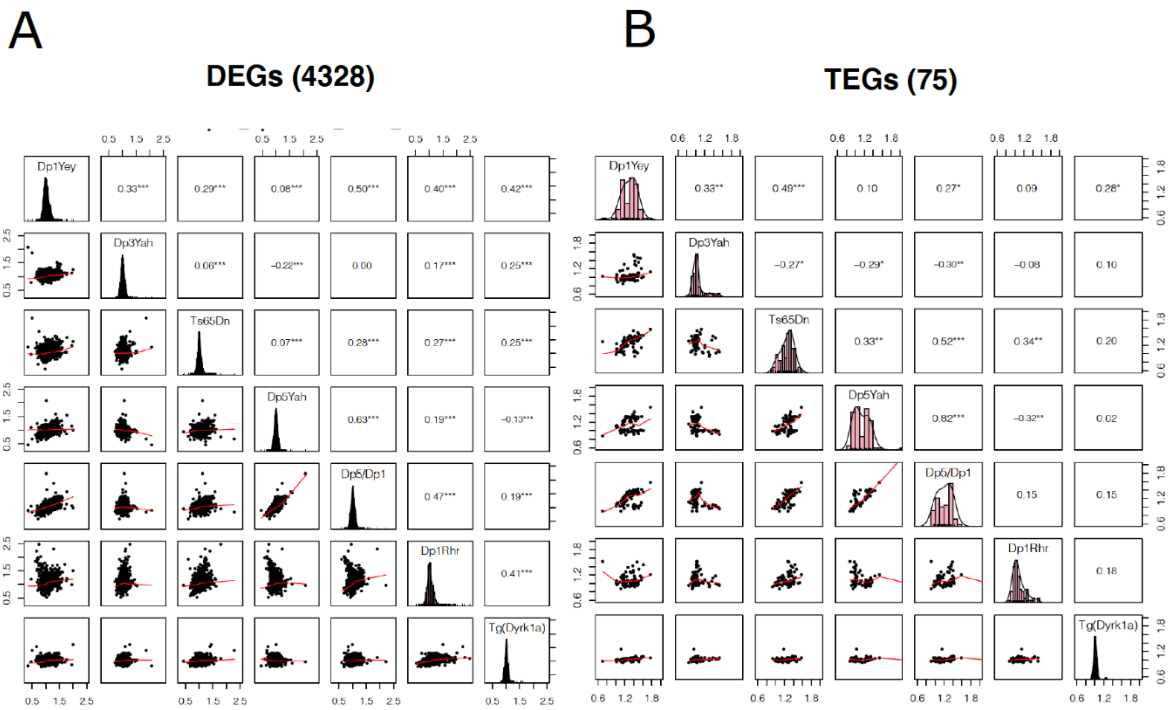


**S9 Fig. Gene expression correlation analyses. (**A) Correlation of the 4328 differentially expressed genes (DEGs) expression levels on the different models. B) Correlation of the 75 differentially expressed trisomic genes (TEGs) expression levels on the different models. Each row represents a model pairwise correlation with the rest of the models. The density and histograms distribution of the gene expression values for each model is found in the medial diagonal (Black for DEGs, and pink for TEGs). The LOESS smoothed fit is represented by a red line. The plots were created in R using the pairs.panel function from Psych R CRAN package (27).

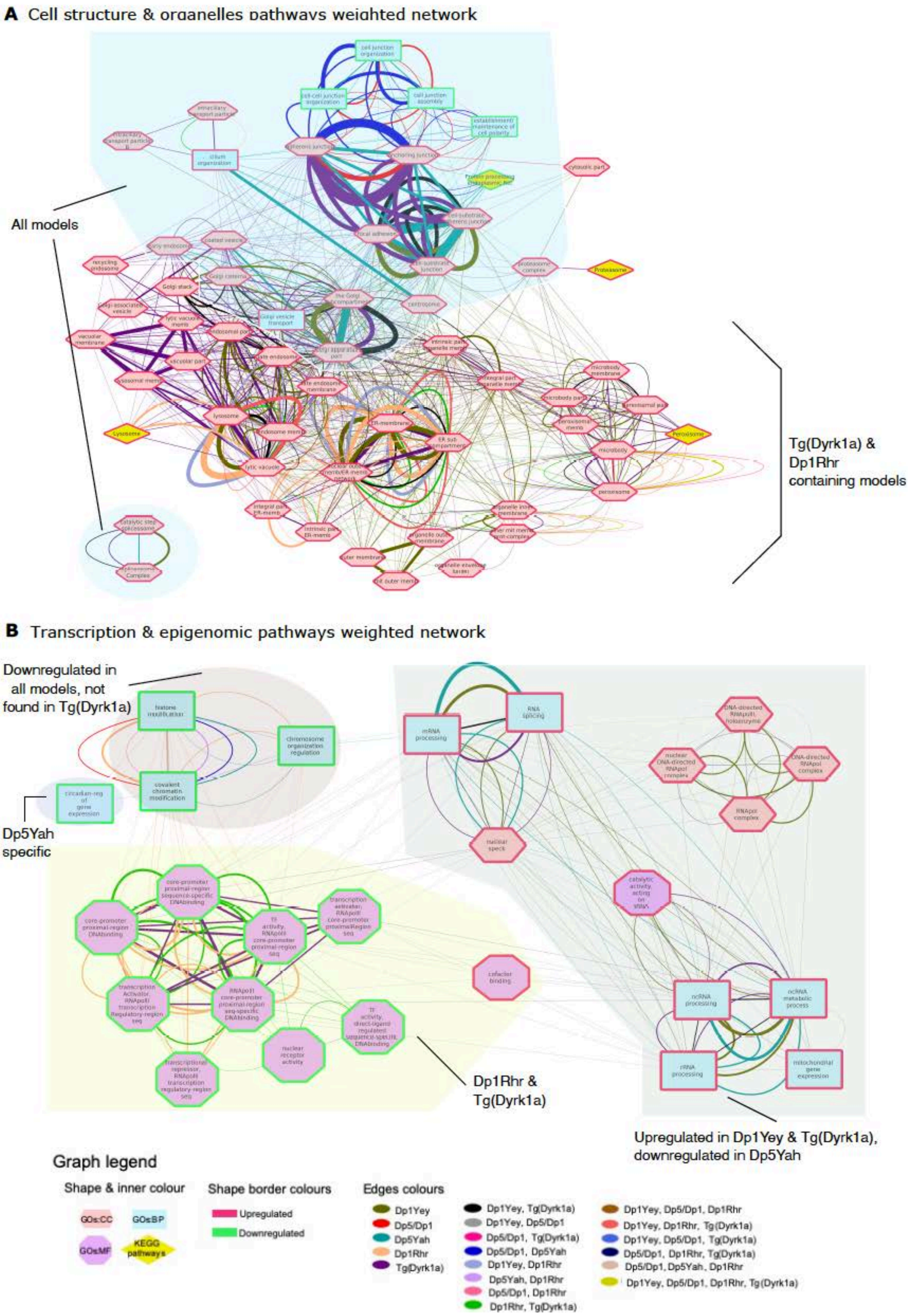


**S10 Fig. Weighted networks representing the connectivity of the pathways included in the Cell structure & organelles, and the Transcription & epigenomic regulation meta-pathways.** (A) Representation of the strength connecting the pathways included on the Cell

Structure & organelle meta-pathway. (B) Representation of the strength connecting the pathways included on the Transcription & epigenomics regulation meta-pathway. The pathways incorporated in each meta-pathway were identified in the differential functional analysis (DFA) by GAGE after imposing a q-val <0.1 cut off. The connectivity between the pathways, (the edges weight or strength connecting the pathways) is represented by the number of altered genes identified by GAGE shared by each two pathways and is calculated based on the number of common genes within the pathways shared inter and intra mouse model (by each mouse model multi group defined) also grouped by regulation sense using the following formula* and represented by the thickness of the edges connecting the nodes (pathways) of the network. Each edges colour depends on the mouse models where those genes were identified (mouse model multi groups). Formula* = number of genes shared by each pathway on each model on each sense of regulation * factorRes with factorRes= 0.6.

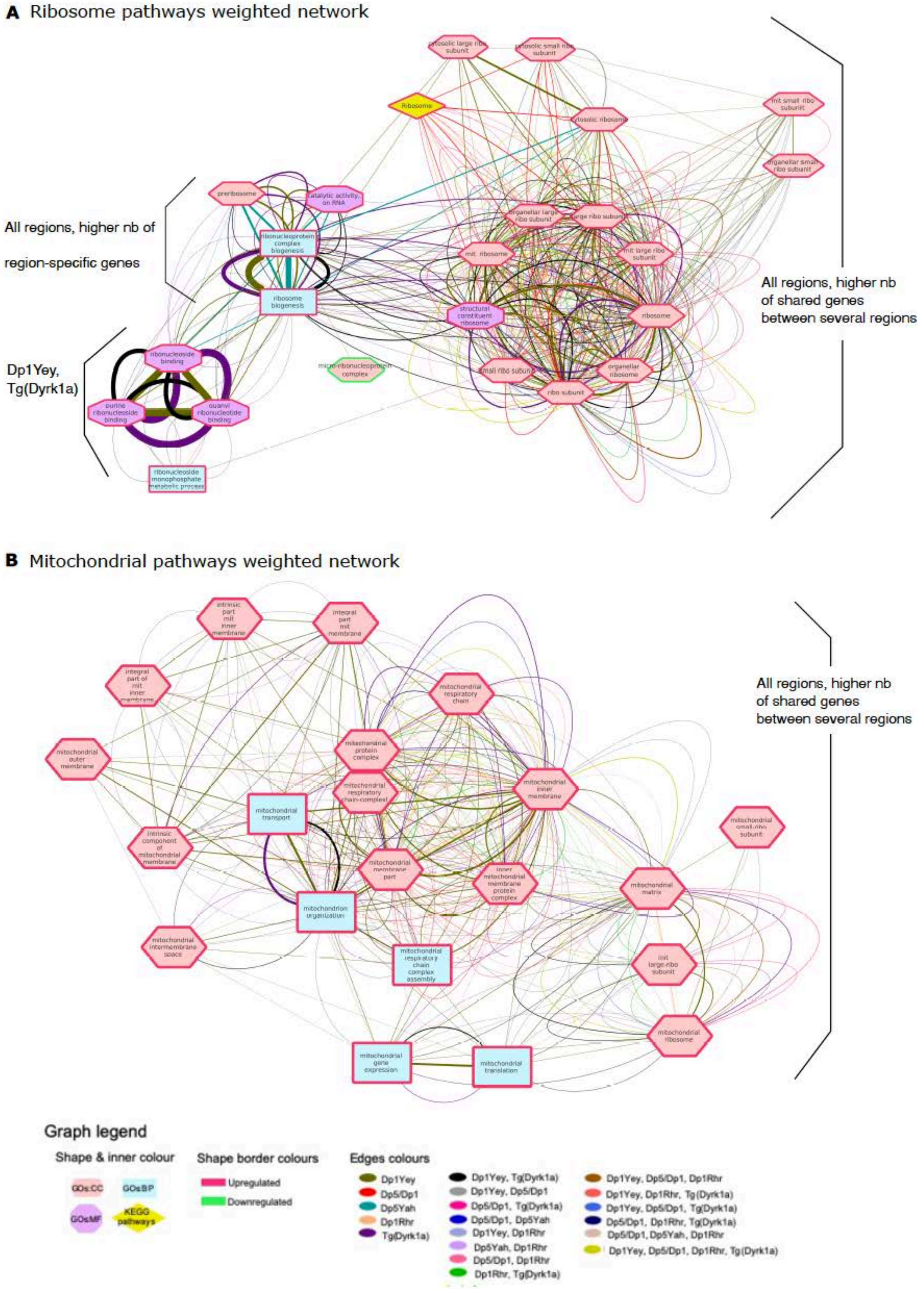


**S11 Fig. Weighted networks representing the connectivity of the pathways included in the Ribosome and Mitochondrial meta-pathways.** (A) Representation of the strength connecting the pathways included on the Ribosome meta-pathway. (B) Representation of the strength connecting the pathways included on the Mitochondria meta-pathway. The pathways incorporated in each meta-pathway were identified in the differential functional analysis (DFA) by GAGE after imposing a q-val <0.1 cut off. The connectivity between the pathways, (the edges weight or strength connecting the pathways) is represented by the number of altered genes identified by GAGE shared by each two pathways and is calculated based on the number of common genes within the pathways shared inter and intra mouse model (by each mouse model multi group defined) also grouped by regulation sense using the following formula* and represented by the thickness of the edges connecting the nodes (pathways) of the network. Each edges colour depends on the mouse models where those genes were identified (mouse model multi groups). Formula* = number of genes shared by each pathway on each model on each sense of regulation * factorRes with factorRes= 0.6

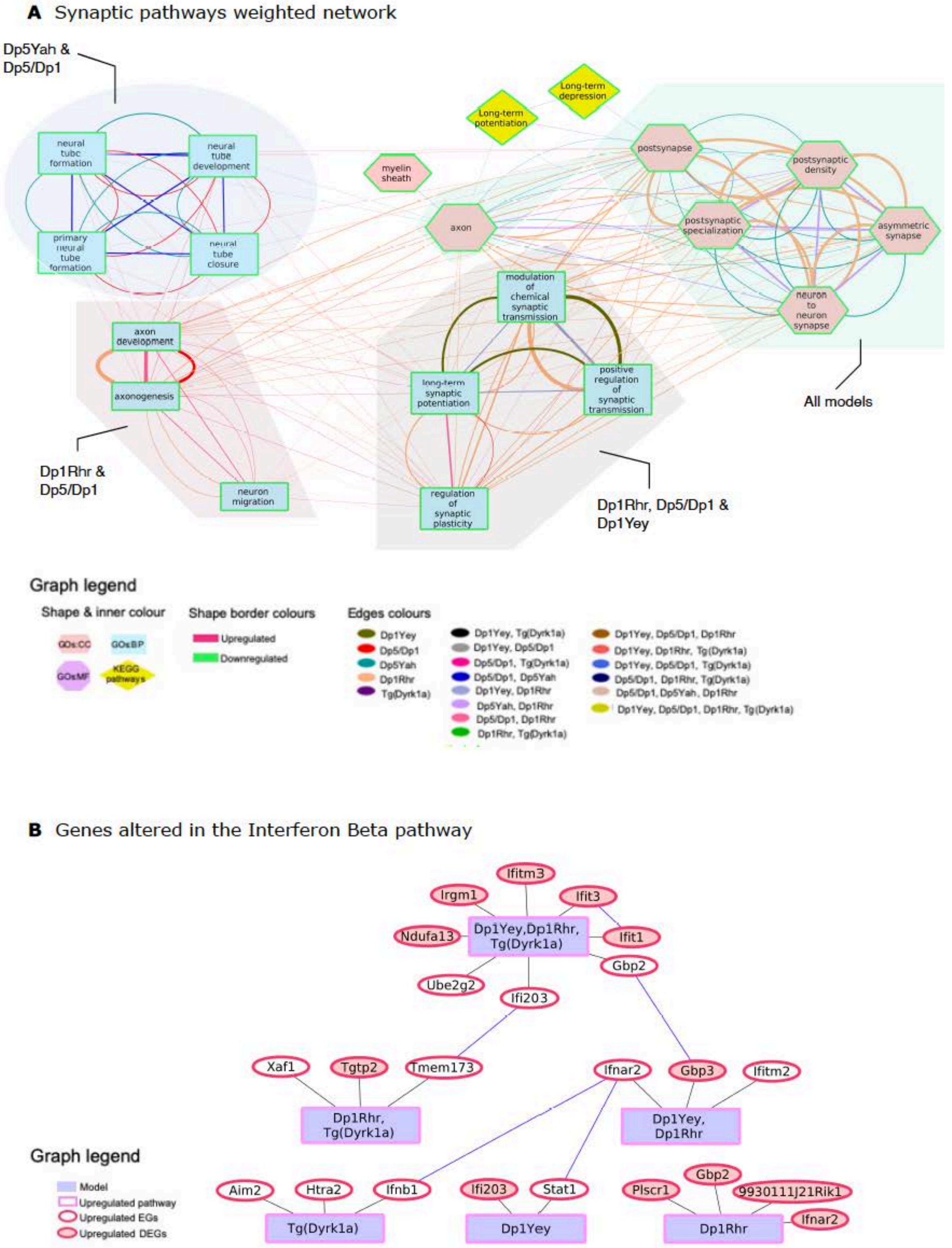


**S12 Fig. Weighted network representing the connectivity of the pathways included in the Synaptic meta-group and the network representation of the genes linked to Interferon Beta (IFN-B) ungrouped pathway.** (A) Representation of the strength connecting the pathways included on the Synaptic meta-pathway. The pathways incorporated in each meta-pathway were identified in the differential functional analysis (DFA) by GAGE after imposing a q-val <0.1 cut off. The connectivity between the pathways, (the edges weight or strength connecting the pathways) is represented by the number of altered genes identified by GAGE shared by each two pathways and is calculated based on the number of common genes within the pathways shared inter and intra mouse model (by each mouse model multi group defined) also grouped by regulation sense using the following formula* and represented by the thickness of the edges connecting the nodes (pathways) of the network. Each edges colour depends on the mouse models where those genes were identified (mouse model multi groups). Formula* = number of genes shared by each pathway on each model on each sense of regulation * factorRes, factorRes= 0.6. (B) Genes linked to Interferon Beta (IFN-B) ungrouped pathway, found altered in the different mouse model multi groups by GAGE. The genes are represented by the ellipse shapes instead the mouse model multi groups by lilac cuadrangles. If the genes were found up regulated then a red boder color was used around the ellipses. The genes also identified by FCROS as differentially expressed (DEG) were identified by a red colored inner ellipse.

A

Shared nodes (Nb)

Shared Seeds (Nb)

| | | | | | |
|---|---|---|---|---|---|
| DYRK1A | 325 | 346 | 28 | 253 | 159 |
| 153 | RHOA | 727 | 86 | 483 | 306 |
| 154 | 154 | GSK3B | 66 | 439 | 316 |
| 22 | 34 | 18 | NPY | 38 | 22 |
| 148 | 158 | 137 | 15 | SNARE | 214 |
| 48 | 57 | 54 | 10 | 51 | NPAS4 |

B

| | | | | | |
|---|---|---|---|---|---|
| DYRK1A | | | | | |
| 47.1 | RHOA | | | | |
| 44.5 | 21.2 | GSK3B | | | |
| 78.5 | 39.5 | 27.2 | NPY | | |
| 58.5 | 32.7 | 31.2 | 39.4 | SNARE | |
| 30.1 | 0.21 | 17.1 | 45.4 | 23.8 | NPAS4 |

Shared Seeds (Nb) /Shared nodes (Nb)

**S13 Fig. Shared interactors between the subnetworks.** (A) Representation of the number of shared seeds (bottom matrix, in soft blue) or shared nodes (seeds plus connecting proteins, upper matrix in dark blue) connecting the subnetworks. (B) Representation of the relevance of the percentage of shared altered genes “seeds” found linked to synaptic function compared to the total number of shared nodes of the subnetworks.

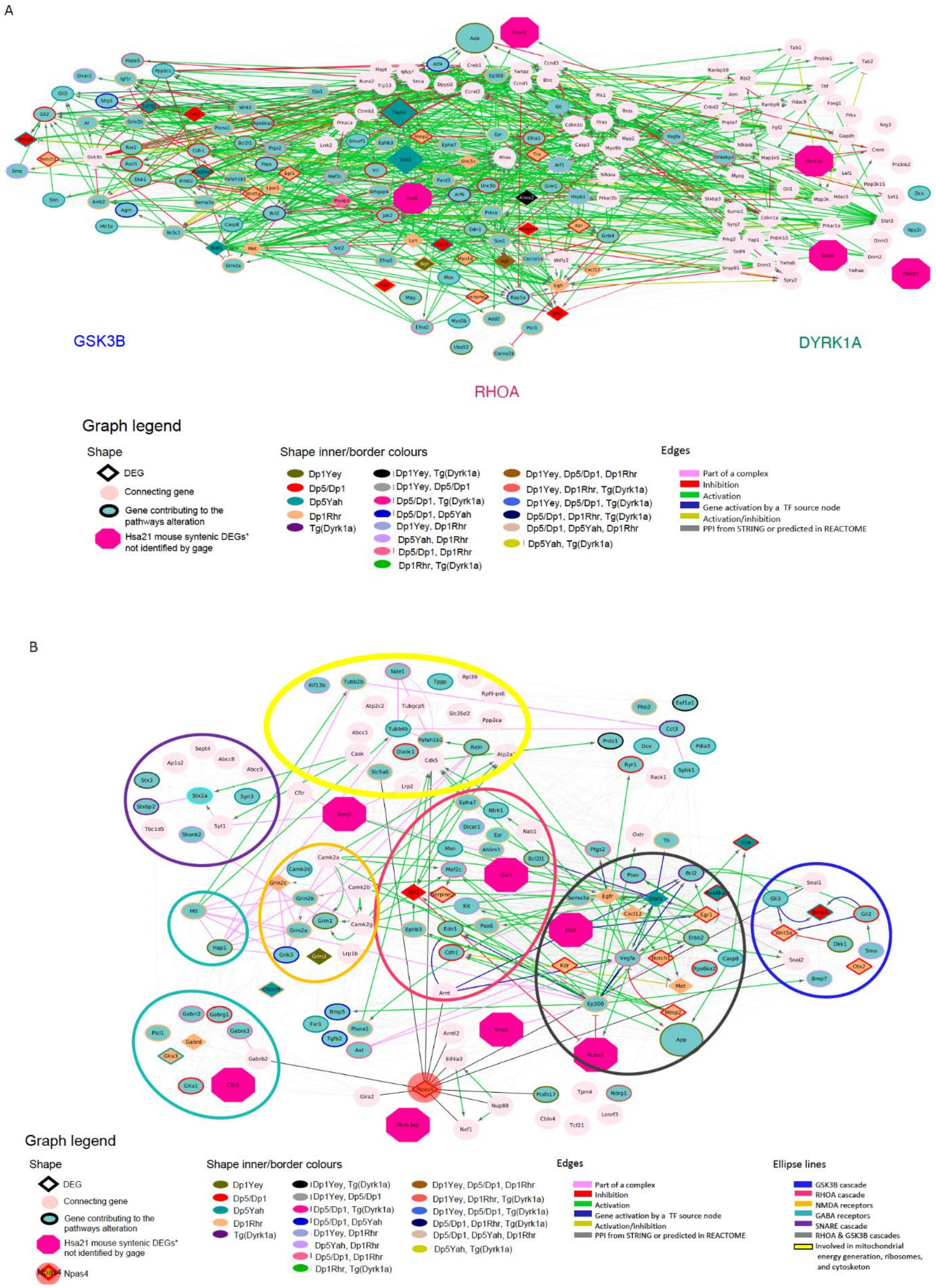
A
GSK3B
DYRK1A
RHOA
Graph legend
Shape
DEG
Connecting gene
Gene contributing to the pathways alteration
Hsa21 mouse syntenic DEGs* not identified by gage
Shape inner/border colours
Dp1Yey
Dp5/Dp1
Dp5Yah
Dp1Rhr
Tg(Dyrk1a)
Dp1Yey, Tg(Dyrk1a)
Dp1Yey, Dp5/Dp1
Dp5/Dp1, Tg(Dyrk1a)
Dp5/Dp1, Dp5Yah
Dp1Yey, Dp1Rhr
Dp5Yah, Dp1Rhr
Dp5/Dp1, Dp1Rhr
Dp1Rhr, Tg(Dyrk1a)
Dp1Yey, Dp5/Dp1, Dp1Rhr
Dp1Yey, Dp1Rhr, Tg(Dyrk1a)
Dp1Yey, Dp5/Dp1, Tg(Dyrk1a)
Dp5/Dp1, Dp1Rhr, Tg(Dyrk1a)
Dp5/Dp1, Dp5Yah, Dp1Rhr
Dp5Yah, Tg(Dyrk1a)
Edges
Part of a complex
Inhibition
Activation
Gene activation by a TF source node
Activation/inhibition
PPI from STRING or predicted in REACTOME
B
Graph legend
Shape
DEG
Connecting gene
Gene contributing to the pathways alteration
Hsa21 mouse syntenic DEGs* not identified by gage
Npas4
Shape inner/border colours
Dp1Yey
Dp5/Dp1
Dp5Yah
Dp1Rhr
Tg(Dyrk1a)
Dp1Yey, Tg(Dyrk1a)
Dp1Yey, Dp5/Dp1
Dp5/Dp1, Tg(Dyrk1a)
Dp5/Dp1, Dp5Yah
Dp1Yey, Dp1Rhr
Dp5Yah, Dp1Rhr
Dp5/Dp1, Dp1Rhr
Dp1Rhr, Tg(Dyrk1a)
Dp1Yey, Dp5/Dp1, Dp1Rhr
Dp1Yey, Dp1Rhr, Tg(Dyrk1a)
Dp1Yey, Dp5/Dp1, Tg(Dyrk1a)
Dp5/Dp1, Dp1Rhr, Tg(Dyrk1a)
Dp5/Dp1, Dp5Yah, Dp1Rhr
Dp5Yah, Tg(Dyrk1a)
Edges
Part of a complex
Inhibition
Activation
Gene activation by a TF source node
Activation/inhibition
PPI from STRING or predicted in REACTOME
Ellipse lines
GSK3B cascade
RHOA cascade
NMDA receptors
GABA receptors
SNARE cascade
RHOA & GSK3B cascades
Involved in mitochondrial energy generation, ribosomes, and cytosketon

**S14 Fig. Protein-protein interaction and regulatory gene connectivity networks (RegPPINets) of genes involved in the synaptic meta-pathway highlighting biological cascades connectivity.** The two sub-networks (A and B) were extracted from the STRING04 MinPPINet. This network was built querying STRING and selecting the PPIs with a medium confidence score (CS), CS=0.4 from all sources of evidence, and was further annotated with regulatory information using REACTOME (See Supplementary material and Methods). The shapes of the nodes represent the following information: Shapes: i)Pallid pink ellipses: represent connecting proteins added to assure the full connectivity of the network; ii) pink octagons, represent HSA21 syntenic genes in mouse not identified as contributing to the meta-pathway dysregulation by GAGE; iii) green inner coloured ellipses, genes identified by GAGE after q-val <0.1 cut off to be contributing even slightly, to any pathway of those found dysregulated inside the meta-pathway. If the size is similar to the octagons, they are also HSA21 syntenic genes in mouse. Additionally, the border colour represents the mouse model multi group where those genes are found altered in; iv) diamonds, genes identified by GAGE after q-val <0.1 cut off and also by FCROS as DEGs. (A) RegPPINet Sub-network extracted from the selection of RHOA 2$^{nd}$ interactors from STRING04 MinPPINet, highlighting RHOA 1$^{st}$ Interactors and DYRK1A and GSK3B interactors annotated with the regulatory information from REACTOME. (B) RegPPINet Sub-network extracted from the selection of NPAS4 2$^{nd}$ interactors from STRING04 MinPPINet. Inside of the ellipsoidal line shapes we highlighted NPAS4 interactors involved in the regulation of GABA receptors (cyan), of NMDA receptors (orange), SNARE complex (purple), RHOA (pink), GSK3B (Blue), both RHOA and GSK3B (black) and the genes linked to mitochondrial, ribosomal and cytoskeleton activity (yellow).

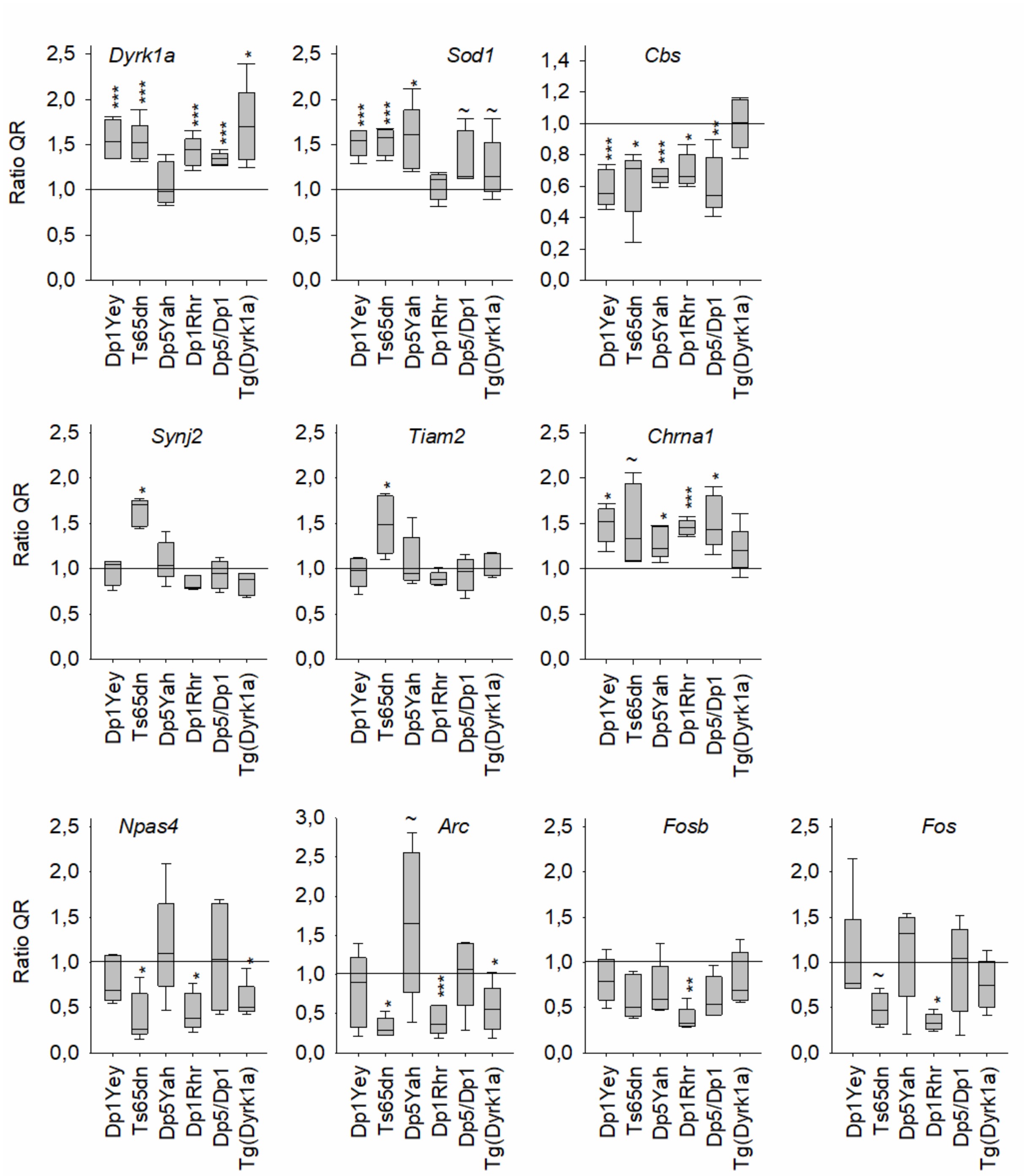


**S15 Fig. Expression level of genes of interest in the different DS models.** Different panels to show the relative ratio (QR) of the expression level of genes of interest in DS models versus wild-type control, located (*Dyrk1a*, *Sod1*) or not in the region homologous to Hsa21. Data are presented as boxplot plots with the median and quartiles (Stastitical analysis was done comparing wt and mutant value with a student t test, * $p<0.05$, ** $p<0.01$, *** $p<0.001$; n=5 per genotype).

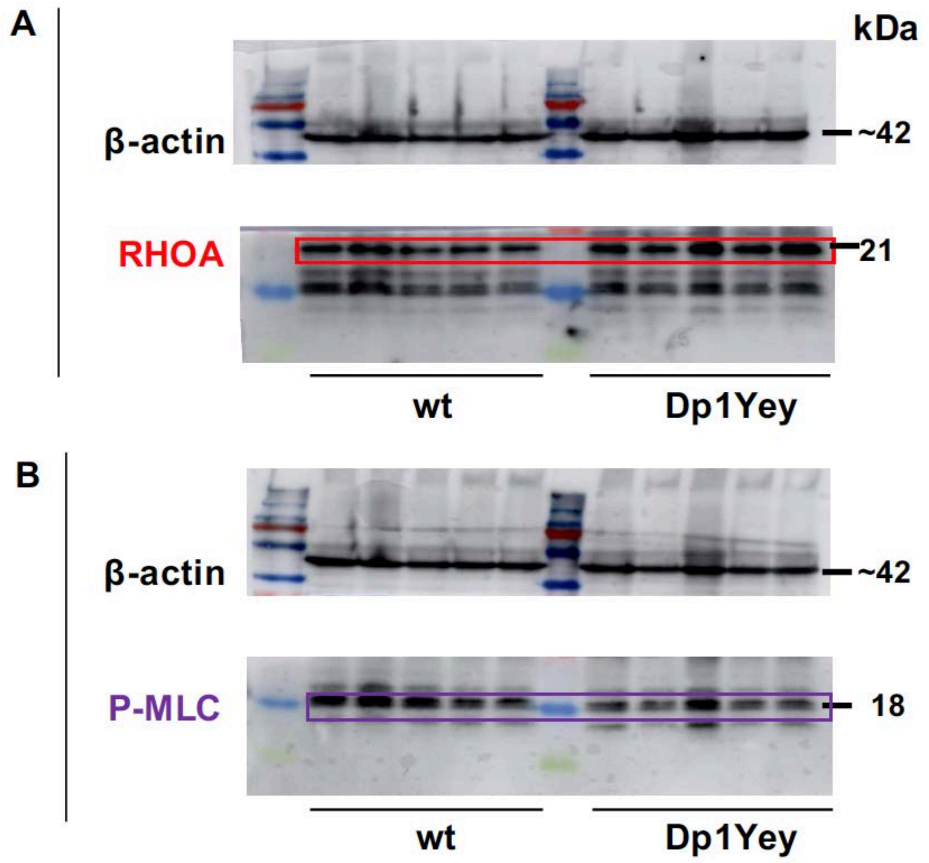


**S16 Fig. Detection of RHOA and the phoshorylated form of Myosin Light Chain (P-MLC) by Western blot in Dp1Yey hippocampal lysates and their control (wt) littermates.** (A) Western blot of RHOA showed no changes in RHOA protein levels in the Dp1Yey line compared to their wt littermates (wt (n=5) and Dp1Yey (n=5)). (B) Western blot of P-MLC revealed a statistically significant decrease (* $p < 0.05$) in the amount of this protein in Dp1Yey mice (n=5 wt and 5 Tg) (Student's t-test: wt vs. Dp1Yey $t(8) = 2{,}392$; $p = 0{,}0437$).

**S1 Table: Summary of behavioural results including statistical assessment.**

**S2 Table. Volumes of each brain structure (in mm3) normalized by total brain volume derived from MRI region-based analysis.**

**S3 Table. Summary of the DEA and DFA results for each trisomic DS model.** We included the number of trisomic genes with compensated or partially compensated expression levels, approximately half of the trisomic genes are compensated and the overall identity of DEGs are different between the models.

**S4 Table. Fold-change of the gene expression in trisomic (Tg) compared to the controls for each mouse model dataset..** The genes included on thoses lists were used to produce the supplementary figs 6 and 7 as follows: i) genes included inside the duplicated genomic regions of chr 16, ii) genes included inside the Ts65Dn chr17 duplicated region iii) genes of the centromeric region of chr17.

**S5 Table. Pathways identified by GAGE as dysregulated including the genes contributing to each altered pathway per each mouse model multi group combination and including the name of the meta-pathway assignated.**

**S6 Table. Number of netork nodes or seeds identified by GAGE as dysregulated included on each main DYRK1A, GSK3B, RHOA, SNARE, NPY, NPAS4 cascades.**

**S7 Table. Full list of R packages used for the bioinformatic pipeline and their citations in R format.**

#### References


1 Duchon, A., Besson, V., Pereira, P., Magnol, L. and Hérault, Y. (2008) Inducing segmental aneuploid mosaicism in the mouse through targeted asymmetric sister chromatid event of recombination. *Genetics*, **180**, 51-59.

2 Hérault, Y., Rassoulzadegan, M., Cuzin, F. and Duboule, D. (1998) Engineering chromosomes in mice through targeted meiotic recombination (TAMERE). *Nat Genet*, **20**, 381-384.

3 Raveau, M., Lignon, J.M., Nalesso, V., Duchon, A., Groner, Y., Sharp, A.J., Dembele, D., Brault, V. and Herault, Y. (2012) The App-Runx1 region is critical for birth defects and electrocardiographic dysfunctions observed in a Down syndrome mouse model. *PLoS genetics*, **8**, e1002724.

4 Michel, D., Chatelain, G., Herault, Y., Harper, F. and Brun, G. (1993) H-DNA can act as a transcriptional insulator. *Cell Mol Biol Res*, **39**, 131-140.

5 Brault, V., Pereira, P., Duchon, A. and Herault, Y. (2006) Modeling chromosomes in mouse to explore the function of genes, genomic disorders, and chromosomal organization. *Plos Genetics*, **2**, 911-919.

6 Olson, L.E., Richtsmeier, J.T., Leszl, J. and Reeves, R.H. (2004) A chromosome 21 critical region does not cause specific Down syndrome phenotypes. *Science*, **306**, 687-690.

7 Li, Z., Yu, T., Morishima, M., Pao, A., LaDuca, J., Conroy, J., Nowak, N., Matsui, S.-I., Shiraishi, I. and Yu, Y.E. (2007) Duplication of the entire 22.9 Mb human chromosome 21 syntenic region on mouse chromosome 16 causes cardiovascular and gastrointestinal abnormalities. *Hum Mol Genet*, **16**, 1359-1366.

8 Brault, V., Duchon, A., Romestaing, C., Sahun, I., Pothion, S., Karout, M., Borel, C., Dembele, D., Bizot, J.-C., Messaddeq, N. *et al.* (2015) Opposite Phenotypes of Muscle Strength and Locomotor Function in Mouse Models of Partial Trisomy and Monosomy 21 for the Proximal Hspa13-App Region. *Plos Genetics*, **11**.

9 Guedj, F., Pereira, P.L., Najas, S., Barallobre, M.J., Chabert, C., Souchet, B., Sebrie, C., Verney, C., Herault, Y., Arbones, M. *et al.* (2012) DYRK1A: A master regulatory protein controlling brain growth. *Neurobiology of Disease*, **46**, 190-203.

10 Hughes, R.N. (2004) The value of spontaneous alternation behavior (SAB) as a test of retention in pharmacological investigations of memory. *Neurosci Biobehav Rev*, **28**, 497-505.

11 Bevins, R.A. and Besheer, J. (2006) Object recognition in rats and mice: a one-trial non-matching-to-sample learning <p>task to study 'recognition memory. *Nat Protoc*, **1**, 1306-1311.

12 Vorhees, C.V. and Williams, M.T. (2006) Morris water maze: procedures for assessing spatial and related forms of learning and memory. *Nat Protoc*, **1**, 848-858.

13 Arbogast, T., Raveau, M., Chevalier, C., Nalesso, V., Dembele, D., Jacobs, H., Wendling, O., Roux, M., Duchon, A. and Herault, Y. (2015) Deletion of the App-Runx1 region in mice models human partial monosomy 21. *Disease Models & Mechanisms*, **8**, 623-634.

14 Crawley, J.N. (1999) Behavioral phenotyping of transgenic and knockout mice: experimental design and evaluation of general health, sensory functions, motor abilities, and specific behavioral tests. *Brain Research*, **835**, 18-26.

15 Goeldner, C., Reiss, D., Wichmann, J., Kieffer, B.L. and Ouagazzal, A.-M. (2009) Activation of nociceptin opioid peptide (NOP) receptor impairs contextual fear learning in mice through glutamatergic mechanisms. *Neurobiology of Learning and Memory*, **91**, 393-401.

16 Ma, Y., Hof, P.R., Grant, S.C., Blackband, S.J., Bennett, R., Slatest, L., McGuigan, M.D. and Benveniste, H. (2005) A three-dimensional digital atlas database of the adult C57BL/6J mouse brain by magnetic resonance microscopy. *Neuroscience*, **135**, 1203-1215.

17 Sawiak, S.J., Wood, N.I., Williams, G.B., Morton, A.J. and Carpenter, T.A. (2009) SPMMouse: A new toolbox for SPM in animal brain. *Proc. Int'l. Soc. Mag. Res. Med.*, 1086.

18 Tustison, N.J., Avants, B.B., Cook, P.A., Zheng, Y.J., Egan, A., Yushkevich, P.A. and Gee, J.C. (2010) N4ITK: Improved N3 Bias Correction. *Ieee Transactions on Medical Imaging*, **29**, 1310-1320.

19 Avants, B.B., Epstein, C.L., Grossman, M. and Gee, J.C. (2008) Symmetric diffeomorphic image registration with cross-correlation: Evaluating automated labeling of elderly and neurodegenerative brain. *Medical Image Analysis*, **12**, 26-41.

20 Warfield, S.K., Zou, K.H. and Wells, W.M. (2004) Simultaneous truth and performance level estimation (STAPLE): An algorithm for the validation of image segmentation. *Ieee Transactions on Medical Imaging*, **23**, 903-921.

21 R_Core_Team. (2019) A language and environment for statistical computing. *R Foundation for Statistical Computing, Vienna, Austria.*

22 Ritchie, M.E., Phipson, B., Wu, D., Hu, Y., Law, C.W., Shi, W. and Smyth, G.K. (2015) limma powers differential expression analyses for RNA-sequencing and microarray studies. *Nucleic Acids Res*, **43**, e47.

23 Carvalho, B. (2015) pd.mogene.1.0.st.v1: Platform Design Info for Affymetrix MoGene-1_0-st-v1. R package version 3.14.1. .

24 Durinck, S., Moreau, Y., Kasprzyk, A., Davis, S., De Moor, B., Brazma, A. and Huber, W. (2005) BioMart and Bioconductor: a powerful link between biological databases and microarray data analysis. *Bioinformatics*, **21**, 3439-3440.

25 Durinck, S., Spellman, P.T., Birney, E. and Huber, W. (2009) Mapping identifiers for the integration of genomic datasets with the R/Bioconductor package biomaRt. *Nat Protoc*, **4**, 1184-1191.

26 Van der Maaten, L. (2014) Accelerating t-SNE using Tree-Based Algorithms. *Journal of Machine Learning Research*, **15**, 3221-3245.

27 Revelle, W. (2019) psych: Procedures for Psychological, Psychometric, and Personality Research. *Northwestern University*.

28 Brault, V., Romestaing, C., Sahun, I., Karout, M., Borel, C., Dembele, D., Bizot, J.-C., Messaddeq, N., Sharp, A.J., Roussell, D. *et al.* (2015) LOCOMOTOR DYSFONCTION AND HYPOTONIA IN DOWN SYNDROME MOUSE MODELS FOR THE HSPA13-APP REGION AS A CONSEQUENCE OF DOSAGE SENSITIVE GENES CONTROLLING MUSCULAR METABOLISM AND MITOCHONDRIAL FUNCTION. *Acta Physiologica*, **214**, 11-12.

29 Dembele, D. and Kastner, P. (2014) Fold change rank ordering statistics: a new method for detecting differentially expressed genes. *Bmc Bioinformatics*, **15**.

30 Perez-Silva, J.G., Araujo-Voces, M. and Quesada, V. (2018) nVenn: generalized, quasi-proportional Venn and Euler diagrams. *Bioinformatics*, **34**, 2322-2324.

31 Swinton, J. Vennerable R package.

32 Subramanian, A., Kuehn, H., Gould, J., Tamayo, P. and Mesirov, J.P. (2007) GSEA-P: a desktop application for Gene Set Enrichment Analysis. *Bioinformatics*, **23**, 3251-3253.

33 Luo, W., Friedman, M.S., Shedden, K., Hankenson, K.D. and Woolf, P.J. (2009) GAGE: generally applicable gene set enrichment for pathway analysis. *BMC Bioinformatics*, **10**, 161.

34 Szklarczyk, D., Morris, J.H., Cook, H., Kuhn, M., Wyder, S., Simonovic, M., Santos, A., Doncheva, N.T., Roth, A., Bork, P. *et al.* (2017) The STRING database in 2017: quality-controlled protein-protein association networks, made broadly accessible. *Nucleic Acids Res*, **45**, D362-D368.

35 Wu, G. and Haw, R. (2017) Functional Interaction Network Construction and Analysis for Disease Discovery. *Methods Mol Biol*, **1558**, 235-253.

36 Xia, J., Gill, E.E. and Hancock, R.E. (2015) NetworkAnalyst for statistical, visual and network-based meta-analysis of gene expression data. *Nat Protoc*, **10**, 823-844.

37 Shannon, P., Markiel, A., Ozier, O., Baliga, N.S., Wang, J.T., Ramage, D., Amin, N., Schwikowski, B. and Ideker, T. (2003) Cytoscape: a software environment for integrated models of biomolecular interaction networks. *Genome Res*, **13**, 2498-2504.

38 Tang, H., Zhong, F., Liu, W., He, F. and Xie, H. (2015) PathPPI: an integrated dataset of human pathways and protein-protein interactions. *Sci China Life Sci*, **58**, 579-589.

39 Vandesompele, J., De Preter, K., Pattyn, F., Poppe, B., Van Roy, N., De Paepe, A. and Speleman, F. (2002) Accurate normalization of real-time quantitative RT-PCR data by geometric averaging of multiple internal control genes. *Genome Biology*, **3**.
